# Fuscimiditide: a RiPP with Ω-Ester and Aspartimide Post-translational Modifications

**DOI:** 10.1101/2021.05.19.444834

**Authors:** Hader E. Elashal, Joseph D. Koos, Wai Ling Cheung-Lee, Brian Choi, Li Cao, Michelle A. Richardson, Heather L. White, A. James Link

## Abstract

Microviridins and other ω−ester linked peptides (OEPs) are characterized by sidechain-sidechain linkages installed by ATP-grasp enzymes. Here we describe the discovery of a new family of OEPs, the gene clusters of which also encode an *O*-methyltransferase with homology to the protein repair catalyst protein L-isoaspartyl methyltransferase (PIMT). We produced the first example of this new ribosomally synthesized and post-translationally modified peptide (RiPP), fuscimiditide, via heterologous expression. NMR analysis of fuscimiditide revealed that the peptide contains two ester crosslinks forming a stem-loop macrocycle. Furthermore, an unusually stable aspartimide moiety is found within the loop macrocycle. We have also fully reconstituted fuscimiditide biosynthesis *in vitro* establishing that ester formation catalyzed by the ATP-grasp enzyme is an obligate, rate-limiting first biosynthetic step. Aspartimide formation from aspartate is catalyzed by the PIMT homolog in the second step. The aspartimide moiety embedded in fuscimiditide hydrolyzes regioselectively to isoaspartate (isoAsp). Surprisingly, this isoAsp-containing protein is also a substrate for the PIMT homolog, thus driving any hydrolysis products back to the aspartimide form. Whereas aspartimide is often considered a nuisance product in protein formulations, our data here suggest that some RiPPs have aspartimide residues intentionally installed via enzymatic activity.

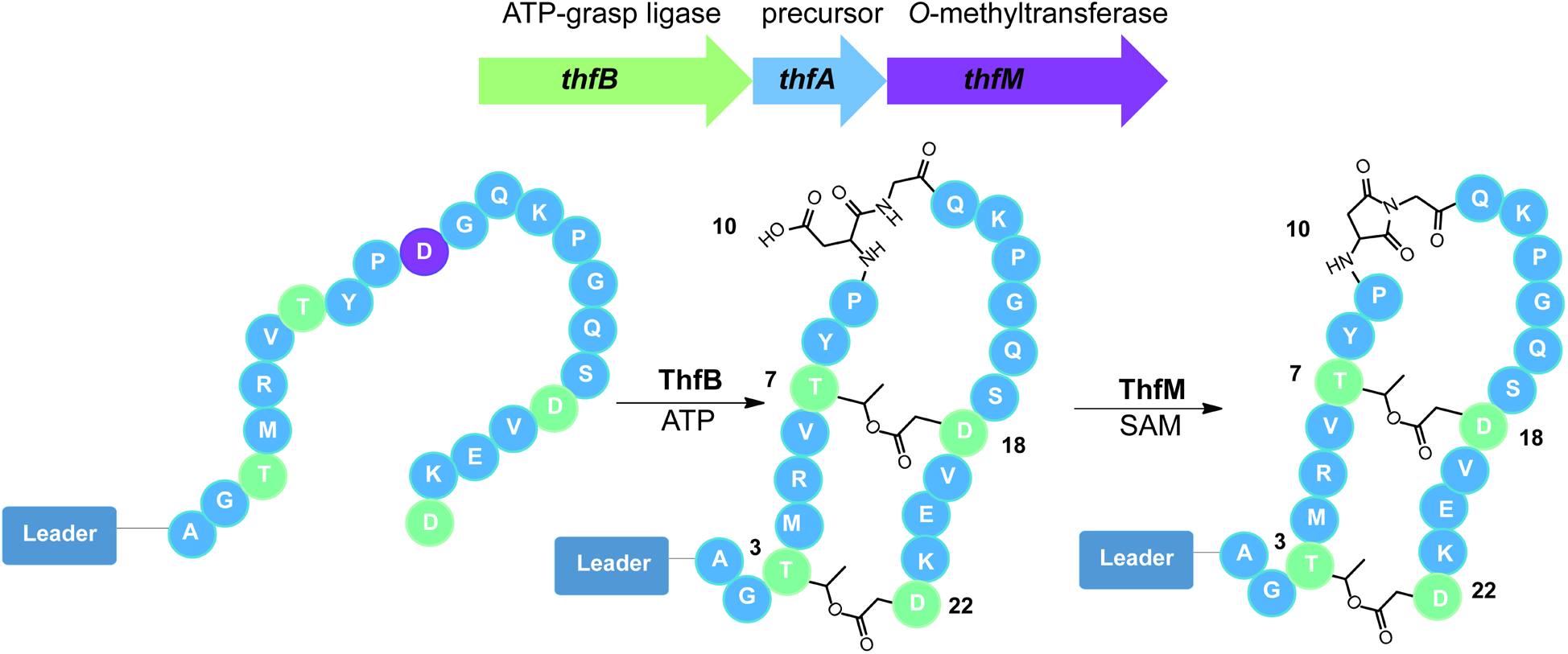

---

Two of the structural hallmarks of ribosomally synthesized and post-translationally modified peptides (RiPPs)^1^ are macrocyclization and backbone modifications. Macrocyclization occurs using backbone-backbone cyclization in the case of cyclotides/orbitides,^2,3^ and cyanobactins,^4^ backbone to sidechain linkages for lasso peptides,^5–8^ sactipeptides,^9,10^ and thiopeptides,^11–13^ and an expanding array of crosslinks between sidechains catalyzed by numerous enzymes.^14–19^ The most well-studied example of backbone modification in RiPPs is the introduction of oxazol(in)e and thiazol(in)e moieties,^20–23^ though other modifications such as epimerization are observed as well.^24^ Both macrocyclization and backbone modification serve to endow RiPPs with improved protease resistance relative to their unmodified counterparts. Many of these products are secreted into hostile, protease-rich environments, and thus require stabilization of their structure. Macrocyclization and backbone modification are also important for the bioactivity of RiPPs. Lasso peptides are comprised of one covalently bonded macrocycle and one mechanically bonded macrocycle,^25^ and this unique structure is required for molecular recognition of targets such as RNA polymerase.^26,27^ Likewise, the thiazoles and oxazoles in linear azol(in)e peptides such as klebsazolicin and phazolicin are involved in base stacking interactions in the ribosome.^28,29^

One well-studied example of macrocyclized RiPPs is the microviridins. The isolation of the first microviridin was published in 1990,^30^ and subsequent work revealed that these peptides are RiPPs.^31–35^ The class-defining modifications for microviridins are two sidechain-sidechain ester linkages and one amide linkage that are installed via the action of two distinct ATP-grasp enzymes.^34,35^ More recent work has uncovered further examples of macrocyclic peptides with only ester crosslinks installed by a single ATP-grasp enzyme, plesiocin^36,37^ and thuringinin.^38^ Given that microviridins and these newer examples all contain sidechain-sidechain ester linkages, Kim and coworkers have named this class of peptides omega ester-containing peptides (OEPs). In a recent bioinformatic survey of OEPs, Kim and coworkers further classified these peptides into 12 distinct groups based on a sequence similarity network constructed from ATP-grasp sequences.^39^ OEP groups 1, 2, and 3 correspond to microviridins, plesiocin, and thuringinin, respectively (Fig. 1a). Characterization of examples from OEP groups 4-6 and 11 has been carried out revealing connectivity patterns different from groups 1-3.^39,40^ There has been a recent call to rename all member of this class graspetides^41^ because of the class-defining ATP-grasp enzyme.

**Fig. 1.**
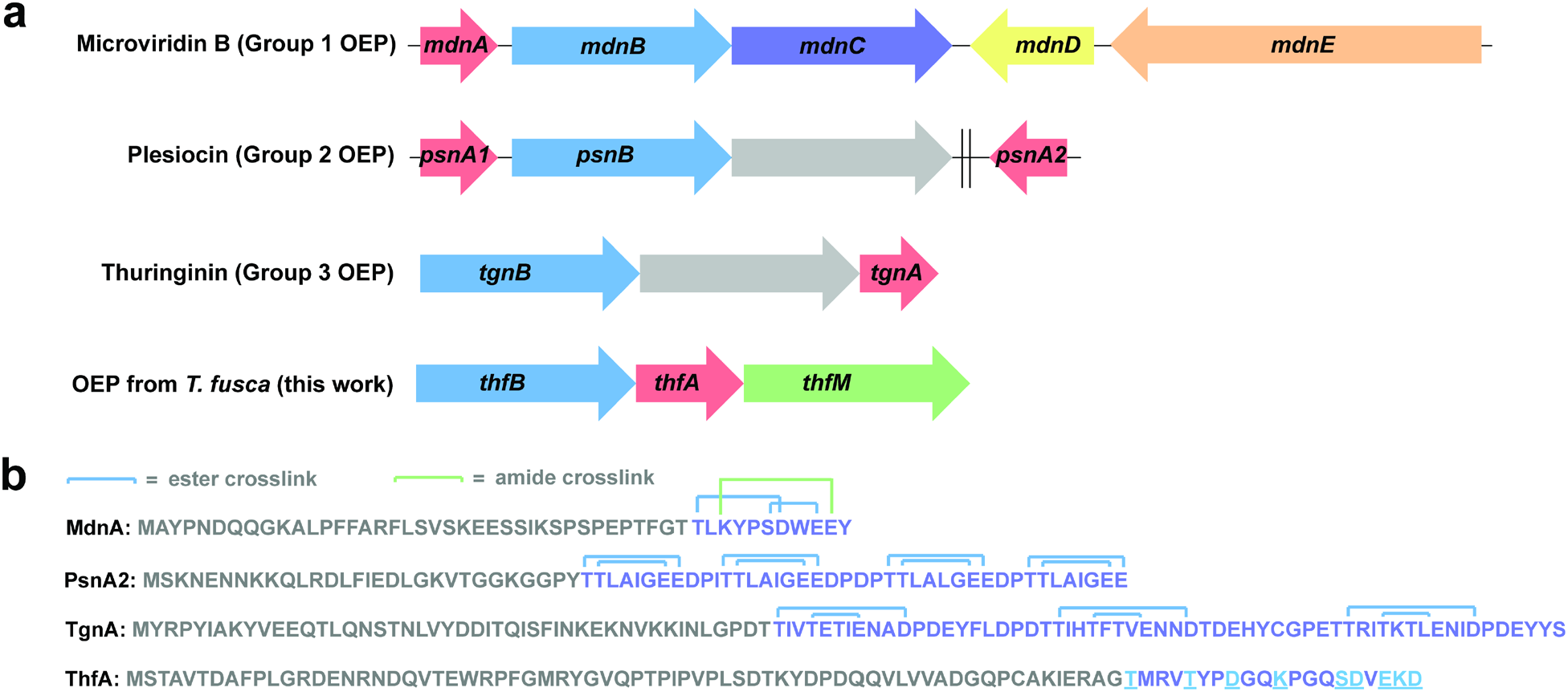
Gene clusters and precursors for microviridins/OEPs. a) Biosynthetic gene clusters (BGCs) for microviridin and other omega ester peptides (OEPs). Red: precursor peptides, light blue: ester-installing ATP-grasp enzymes. In the microviridin B BGC, *mdnC* encodes for an amide-installing ATP-grasp enzyme (purple), *mdnD* encodes for an acetyltransferase (yellow), and *mdnE* encode for a protease/ABC transporter (peach). The plesiocin and thuringinin BGCs harbor a gene encoding a membrane protein of unknown function (gray). The *psnA2* gene is distal from the plesiocin BGC as indicated by the double line. BGC for the putative OEP from *T. fusca* is distinguished by the presence of an *O*-methyltransferase gene (green). b) Precursor protein sequences for the BGCs in part a. Putative leader and core sequences are shown in gray and purple respectively. Ester and amide connectivity is shown in the core peptide for microviridin B, plesiocin, and thuringinin, while potential crosslinkable residues are underlined in the *T. fusca* precursor ThfA.

Here, we describe the discovery and characterization of a new example of an OEP/graspetide, fuscimiditide, from the thermophilic actinobacterium *Thermobifida fusca*. The biosynthetic gene cluster (BGC) for fuscimiditide includes a precursor protein ThfA, a single ATP-grasp enzyme ThfB, and an *O*-methyltransferse ThfM with homology to the protein repair catalyst protein L-isoaspartyl methyltransferase (PIMT). We show through a series of heterologous expression and *in vitro* enzymology experiments coupled with detailed NMR analysis that fuscimiditide is a stem-loop structure peptide macrocycle with two sidechain-sidechain ester linkages and an aspartimide modification within the backbone of the loop macrocycle. ThfM methylates an Asp residue and the methyl ester is subsequently attacked by the adjacent backbone to form an aspartimide. This is in contrast to native PIMTs which have a strong preference for isoaspartate (isoAsp) as a substrate. Whereas aspartimide is typically viewed solely as an intermediate in age-related protein damage and its subsequent repair,^42–45^ our data shows that the aspartimidylated peptide is the desired final product of the BGC *in cellulo*. While the aspartimide residue in fuscimiditide is stable for days in neutral, low ionic strength conditions, it hydrolyzes regioselectively to an isoAsp moiety under higher ionic strength or basic conditions. Remarkably, ThfM is also able to recognize isoAsp and drive the hydrolyzed material back to the aspartimidylated form, showing that ThfM has unprecedented promiscuity among PIMT enzymes.

## Results

### A microviridin/OEP-like BGC in *T. fusca* harbors an uncharacterized methyltransferase

Our lab has previously worked with *Thermobifida fusca*, a thermophilic actinobacterium, in our studies on fuscanodin, a lasso peptide.^46^ In a closely related organism, *Thermobifida cellulosilytica*, there is a lasso peptide biosynthetic gene cluster (BGC) with an *O*-methyltransferase in addition to the minimal biosynthetic genes for lasso peptides.^47^ We searched *T. fusca* for similar *O-*methyltransferases and found one associated with an ATP-grasp enzyme and a putative RiPP precursor. In a previous bioinformatic survey of ATP-grasp enzyme prevalence, the colocalization of an ATP-grasp enzyme, an *O-*methyltransferase, and a precursor protein was noted in actinobacteria.^48^ To our knowledge, there have been no investigations into the products of these BGCs. The putative precursor protein in *T. fusca*, the gene for which will be referred to as *thfA*, is 89 aa long and enriched in Thr, Ser, Asp, and Glu residues at its C-terminus, consistent with it being an OEP-like substrate (Fig. 1b). In this BGC, the precursor is found in between the gene encoding the ATP-grasp enzyme (ThfB) and the methyltransferase (ThfM) (Fig. 1a). The pattern of residues at the C-terminus of ThfA is different from the patterns in any of the 12 groups of OEPs delineated by Kim and coworkers (Fig. 1b).^39^ This fact, coupled with our curiosity about the function of the *O*-methyltransferase, prompted us to study the *thfBAM* BGC in more detail.

### Heterologous expression of Thf BGC

To determine whether ThfB and ThfM were able to modify the ThfA precursor protein, we developed a coexpression system in *E. coli*. An N-terminal His-tag was added to ThfA and the gene was placed under an IPTG-inducible T7 promoter. The biosynthetic enzymes ThfB and ThfM were cloned individually under an IPTG-inducible T5 promoter. Utilizing *E. coli* BL21 (DE3) Δ*slyD* as the host strain, ThfA was heterologously expressed and purified with a yield of ~3.3 mg/L (Fig. S1). Following purification, LC-MS analysis of the precursor revealed an observed mass of 11,361 Da corresponding to the full protein with cleavage of the N-terminal methionine (Fig. 2a). Next, we sought to determine the extent of the PTMs installed by ThfB and ThfM and whether there was a specific order of these modifications. Coexpression of the precursor ThfA and ATP-grasp ThfB yielded an observed average mass of 11,325 Da indicating a loss of 36 Da corresponding to a 2-fold dehydrated product; minor amounts of 11,361 Da (unmodified) and 11,343 Da (one dehydration) were also detected (Fig. 2a). On the other hand, coexpression with the *O*-methyltransferase ThfM did not yield any modification on ThfA (Fig. S2). When all three components of the biosynthetic pathway were coexpressed, we expected to see a product corresponding to doubly dehydrated ThfA with addition of a methyl group (11,339 Da). Surprisingly, the major product of the heterologous expression was a 3-fold dehydrated precursor with an observed mass of 11,307 Da (Fig. 2a). These experiments established that the ATP-grasp ThfB was responsible for installing two dehydrations, while the *O*-methyltransferase ThfM installed a third dehydration (Fig. 2a). Notably, ThfM only recognizes the doubly dehydrated ThfA that has been modified by ThfB, establishing a strict order of modification.

**Fig. 2.**
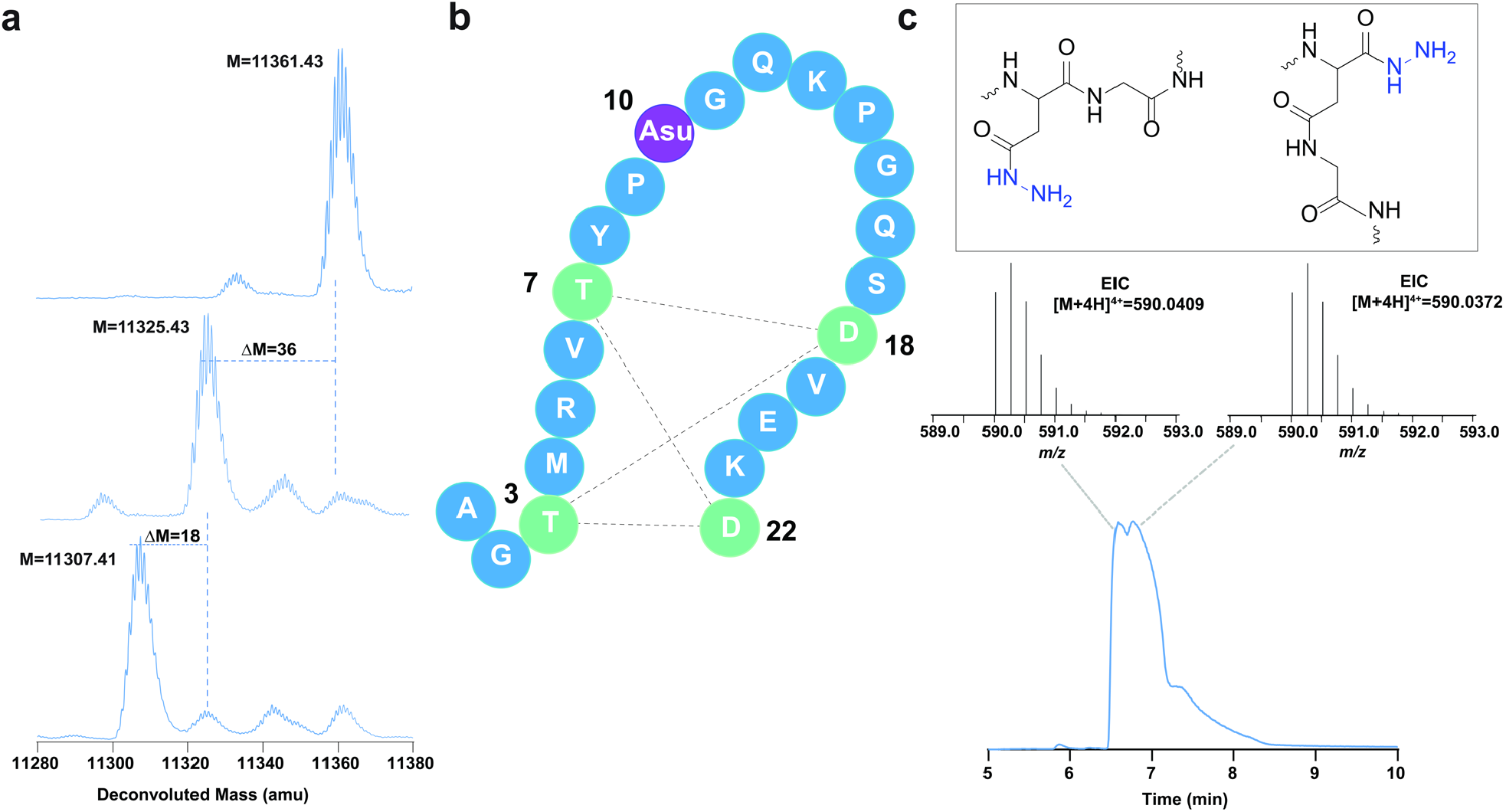
Posttranslational modifications (PTMs) on ThfA. a) Deconvoluted mass spectra of unmodified ThfA (top), ThfA co-expressed with ATP-grasp enzyme ThfB (middle), and ThfA co-expressed with ThfB and methyltransferase ThfM (bottom). ThfB dehydrates ThfA twice, while ThfM adds a third dehydration. b) The three dehydrations on ThfA are localized to the C-terminal 22 aa, the putative core peptide. MS/MS analysis (Fig. S5) indicates a macrocyclic diester structure stretching from T3 to D22 with the third dehydration being an aspartimide at D10 (Asu). c) Products of hydrazine addition to the 22 aa, triply dehydrated ThfA core peptide. Two distinct species with masses corresponding to the peptide plus a single hydrazide are observed, consistent with hydrazine addition to two different locations on aspartimide. Top: extracted ion current (EIC) mass spectra of the two species. Bottom: total ion current (TIC) chromatogram of the hydrazine/triply dehydrated core peptide reaction.

### Characterization of PTMs on ThfA

In order to identify the posttranslational modification (PTMs) in doubly and triply dehydrated ThfA, we examined the ThfA sequence for potential trypsin cleavage sites. Digestion of the 2-fold dehydrated ThfA yielded a C-terminal 22 amino acid peptide containing 2 dehydrations and 3 missed trypsin cleavages (Fig. S3). Similarly, digestion of the 3-fold dehydrated ThfA yielded the same fragment with 3 dehydrations marking the digested product as the putative core peptide of ThfA (Fig. S3). Following identification of the core sequence, we sought to gather more evidence regarding the chemical nature of the installed PTMs. Since ATP-grasp ligases are known to catalyze the formation of esters and/or amides in OEPs, we reasoned that stability studies conducted at varying pHs could differentiate between these two possibilities. Therefore, we incubated doubly and triply dehydrated ThfA in 1x PBS buffer for 21 hours at 25 °C at a range of pHs; pH 3-11 for the doubly dehydrated species and pH 7-11 for the triply dehydrated species. The 2-fold dehydrated ThfA was stable at pH 3-10; partial hydrolysis was only observed at pH 11 indicating that the installed PTMs were likely ester linkages (Fig. S4). Meanwhile, hydrolysis of the third dehydration (installed by ThfM) was observed at pH 7-11 suggesting a more labile modification (Fig. S4).

Following these stability studies, we performed MS/MS analysis of doubly and triply dehydrated ThfA core peptide. In previous work on OEPs, the ester linkages were cleaved during MS/MS analysis, allowing for fragmentation patterns that assist in position determination of the ester forming residues.^36,38^ However, we did not observe cleavage of doubly dehydrated ThfA core peptide under collision-induced dissociation (CID) conditions. Instead, we observed minimal fragmentation yielding the *b*_2_, *y*_20_, and *y*_21_ ions in addition to several fragments corresponding to internal peptide cleavages between Pro9 and Pro14 of the core peptide (Fig. S5).^49^ The lack of fragmentation beyond the observed *b*/*y* ions suggested the involvement of Thr at position 3 and Asp in position 22 (Fig. 2b). Moreover, the peptide fragmentation pattern coupled with the fact that there were 3 missed trypsin cleavages indicated macrocyclization of the peptide (Fig. S3). The triply dehydrated ThfA core exhibited an identical fragmentation pattern except for one internal cleavage fragment between Asp10 and Gly11, which was no longer observed.

Following the line of evidence indicating that the third PTM is labile and the lack of observable MS/MS fragmentation between Asp10 and Gly11 in triply dehydrated core, we hypothesized that the dehydration installed by ThfM could be an aspartyl succinimide (Asu), also known as aspartimide (Fig. 2b). ThfM belongs to a family of *O*-methyltransferases with homology to protein repair catalysts protein L-isoaspartyl methyl transferases (PIMTs). The usual role of PIMTs is salvaging aging proteins that have acquired backbone isoAsp residues via spontaneous asparagine deamidation or aspartate dehydration with subsequent hydrolysis (Fig. S6). The PIMT enzyme specifically methylates isoAsp, which is followed by non-enzymatic formation of an aspartyl succinimide. The succinimide can open to either Asp or isoAsp, with repeated cycles of catalysis leading to reversion of most of the protein to its Asp form. To test whether triply dehydrated ThfA core contained an aspartimide, we incubated the doubly and triply dehydrated core peptides with a 2 M hydrazine solution in water at room temperature for 30 mins. Given the selective reactivity of hydrazine with succinimides under the abovementioned conditions, a mass shift of +32 *m/z* was expected.^50,51^ LC-MS analysis of the reaction products indicated partial hydrolysis of doubly dehydrated core owing to increased basicity of the reaction solution, while triply dehydrated core exhibited a mass change of +32. Hydrazine trapping of the putative aspartimide yielded two distinct peaks corresponding to opening of the succinimide on two different sides (Fig. 2c). MS/MS analysis of hydrazine reacted core yielded the abovementioned fragmentation pattern with an additional +32 mass change corresponding to hydrazide addition on a residue between Pro9 and Lys13 (Fig. S7). Taken together, this MS evidence points to the formation of aspartimide at Asp10 in triply dehydrated ThfA core.

Additionally, the secondary structures of unmodified linear core peptide (obtained by solid-phase peptide synthesis) as well as doubly and triply dehydrated core peptide (obtained by co-expression followed by tryptic digest) were analyzed using CD spectroscopy. Linear core peptide displayed the signature of a random coil, while both dehydrated peptides exhibited β character presumably endowed by the ester crosslinking (Fig. S8a). We also analyzed the entire precursor ThfA using CD and were surprised to see that this protein exhibited a random coil structure (Fig. S8b) given its stability during heterologous expression in *E. coli*.

To further confirm the presence of 2 ester linkages, we executed an alanine scan of all residues with side chains capable of forming amides or esters. Swaps to alanine of Thr3, Thr7, Asp18, and Asp22 in the core peptide led to products that were no longer doubly dehydrated (Fig. S9) strongly suggesting the involvement of these residues in ester cross-linkages. Thus, the macrocycle formed in doubly and triply dehydrated ThfA is predicted to span 20 aa from Thr3 to Asp22; two different linkage patterns are possible (Fig. S10). As discussed above, the linkage pattern observed in this macrocycle differs from any of the known or predicted classes of OEPs.^39^

### NMR structures of doubly and triply dehydrated ThfA core peptide

Following preliminary structural analysis of the doubly and triply dehydrated peptides, TOCSY, NOESY, HSQC, and HMBC spectra were acquired on the doubly dehydrated ThfA core peptide in 98:2 H_2_O:D_2_O at 25°C (Fig. S11-S14). The TOCSY and NOESY spectra were used for sequential assignment of amino acid protons in addition to providing key pieces of evidence as to amino acid interactions in three-dimensional space (Table S2). In the NOESY spectra, strong NOEs could be observed between the amide proton of Asp18 and the amide proton of Thr7, the amide proton of Asp22 and the α-proton of Thr3, and the amide proton of Thr7 and the α-proton of Asp18 (Table S3). Additionally, a pronounced downfield shift was observed for the β-proton of Thr3 and Thr7 as compared to its average statistical value^52^ (Table S2). HMBC correlations were observed between the β-proton of Thr3 and Thr7 and the Asp22 and Asp18 side chain carbonyl carbons, respectively. Strikingly, the Asp18 and Asp22 side chain carbonyls were shifted upfield to 169.0 ppm and 169.5 ppm indicating esterification (Fig. S15, Table S2). Mutagenesis results coupled with NOE and HMBC correlations confirmed that ThfA installed two sidechain-sidechain ester linkages: Thr3-Asp22 and Thr7-Asp18. Following assignment, CYANA 2.1 was used to calculate the 20 lowest energy conformers of the ester linked peptide. The peptide was shown to form a stem-loop structure with anti-parallel β character in the stem, thereby validating the secondary structure observed by CD spectroscopy (Fig. 3a). The modeled CYANA structure shows that the peptide formed a bis-macrocyclic structure, where the stem macrocycle was composed of 34 backbone atoms, while the loop macrocycle contained 32. The Tyr8 to Ser17 residues, corresponding to the loop of the stem-loop structure, exhibited few NOEs to non-adjacent residues, suggesting that this macrocycle did not extensively fold back upon the stem. This structure differs substantially from microviridins, however group 2 and 3 OEPs are predicted to also exhibit stem-loop elements in their structure, albeit with different macrocycle sizes.^36,38^ This work represents the first comprehensive NMR analysis of an ATP-grasp modified stem-loop structure.

**Fig. 3.**
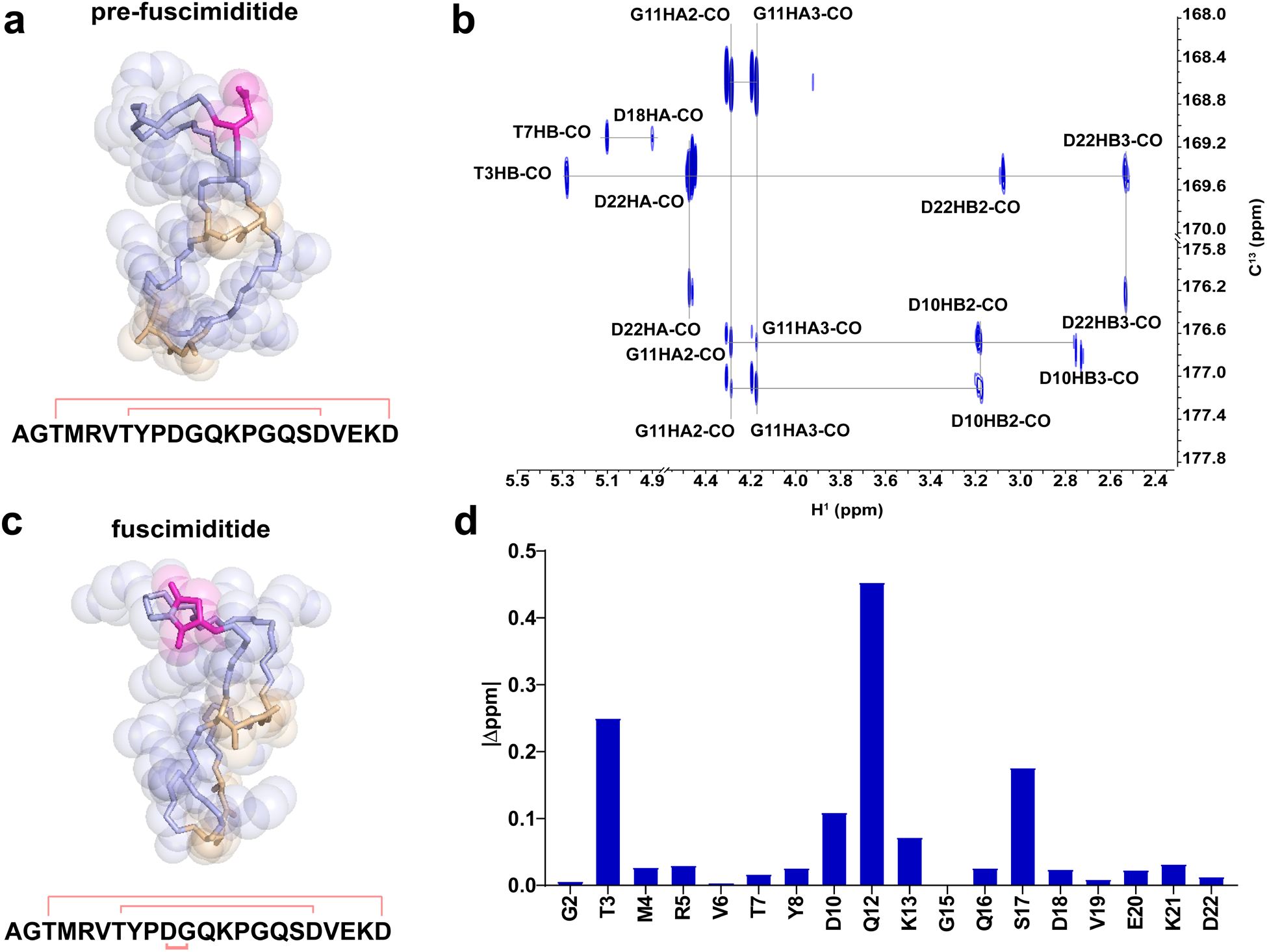
NMR structures of pre-fuscimiditide and fuscimiditide. a) Top structural model of pre-fuscimiditide, the ThfA core peptide with two ester linkages, as computed by CYANA 2.1. Ester linkages (wheat) connect T3 to D22 and T7 to D18 forming a stem-loop structure comprised of two edge-to-edge macrocycles. The peptide backbone is colored light blue; the D10 sidechain is in magenta. An overlay of the top 20 structures is in Fig. S16. b) Portion of the ^1^H-^13^C HMBC spectrum of fuscimiditide, ThfA core peptide with two esters and an aspartimide. Key correlations that show the ester and succinimide connectivity are labeled. c) Top structural model of fuscimiditide. The color scheme is as in part a; the succinimide moiety is now colored magenta. An overlay of the top 20 structures is in Fig. S16. d) Chemical shift deviations of amide protons between pre-fuscimiditide and fuscimiditide. A1, P9, G11, and P14 are not shown because of a lack of amide protons on these residues.

Having established the ester connectivity in doubly dehydrated ThfA core peptide, we sought to use NMR to identify the location of the additional dehydration installed by the *O*-methyltransferase (Fig. S17-S19). Our MS and hydrazine addition data discussed above (Fig. S5 and S7) suggested that the Asp10 side chain was being converted to an aspartimide residue. In the TOCSY and NOESY spectra for triply dehydrated ThfA core peptide, the amide proton of Gly11 could no longer be observed. This suggests the existence of a stable aspartimide linking the side chain of Asp10 and the backbone of Gly11 (Table S2). More directly, HMBC correlations were observed for the Gly11 α-protons and the Asp10 backbone and side chain carbonyls (Fig. 3b) confirming that *O*-methyltransferase catalysis facilitates the formation of the peptide aspartimide. The correlations and chemical shift values we observed for the succinimide moiety are in good agreement with published data on model peptides.^53^ Upon confirmation of the aspartimide residue within the triply dehydrated peptide, we decided to name it fuscimiditide which reflects both the organism that encodes it in its genome as well as its unique aspartimide group. The doubly dehydrated ThfA core peptide containing only esters will be referred to as pre-fuscimiditide (Fig. 3a). Utilizing proton assignments and NOESY constraints, fuscimiditide was modeled to be a stem-loop structure with the aspartimide positioned in the loop macrocycle (Fig. 3c). Notably, the aspartimide lies within a 6 aa stretch, PDGQKP, making this region especially rich in residues containing backbone constrained side chains. When comparing chemical shift deviations between pre-fuscimiditide and fuscimiditide, the largest chemical shift deviations were observed for the Gly11 and Gln12 α-protons and the Gln12 amide proton (Fig. 3d, Table S4). The aspartimide had an especially strong influence on the chemical environment of the Gln12 backbone as indicated by the downfield shift of its amide proton by 0.45 ppm.

### Fuscimiditide contains an unusually stable aspartimide

As discussed above, aspartimides are generally observed as intermediates in protein repair. However, our data strongly suggests that the aspartimidylated peptide may be the final intended product of the fuscimiditide BGC. In fuscimiditide, the aspartimide moiety was able to withstand purification from cells (even under denaturing conditions), 1 h tryptic digests, and acquisition of multiple NMR spectra, prompting us to study its stability in more depth. Incubation of fuscimiditide in water, at RT, for three days resulted in less than 1% hydrolysis indicating that the aspartimide was stable in low ionic strength conditions. However, as discussed above, incubation of the peptide in PBS at varying pH values resulted in more rapid hydrolysis of the aspartimide (Fig. S4), while the ester linkages remained intact. Hydrolysis products were also observed in tryptic digests that were carried out for ~16 hours in 50 mM ammonium bicarbonate (Fig. 4a). During reverse-phase HPLC purification of the hydrolyzed fuscimiditide, two hydrolysis products with different retention times were observed. The minor peak (~5% of the hydrolysate) had a retention time identical to pre-fuscimiditide, whereas the major peak (~95% of the hydrolysate) eluted differently (Fig. S20). The most obvious explanation for these two peaks is the formation of a mixture of Asp and isoAsp in the backbone of fuscimiditide upon hydrolysis (Fig. 4b). Our data suggest that the aspartimide of fuscimiditide regioselectively opens primarily to the isoAsp form.

**Fig. 4.**
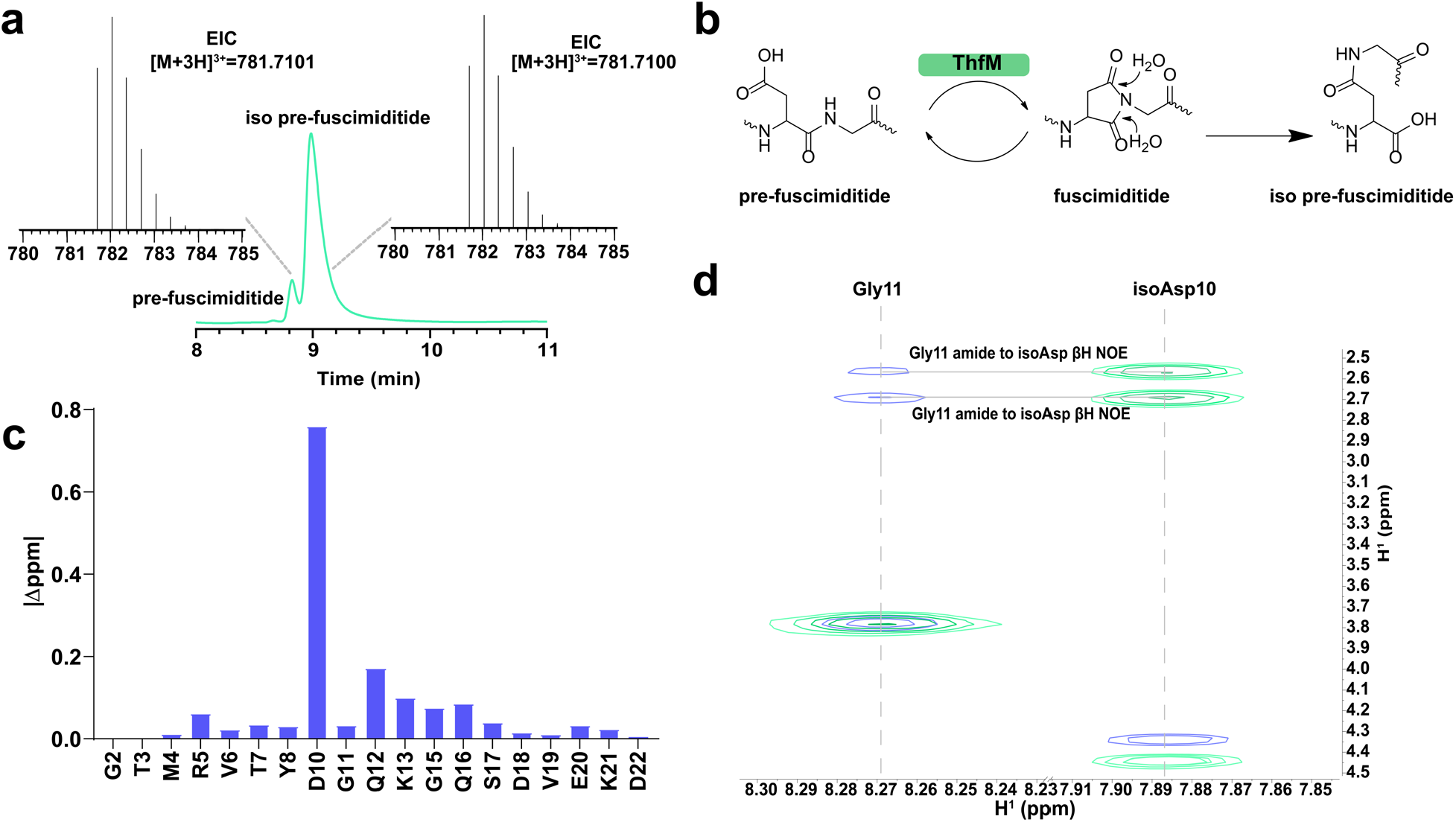
Characterization of the major fuscimiditide hydrolysis product, iso pre-fuscimiditide. a) HPLC trace of the products of fuscimiditide hydrolysis. The small peak at 8.8 min corresponds to the retention time of pre-fuscimiditide, but the major peak elutes later. These two peaks were collected and have identical masses according to LC-MS (insets). b) The aspartimide moiety in fuscimiditide (center) can hydrolyze to either Asp (left) or isoAsp (right). Succinimide formation enabled by ThfM catalysis is also shown. c) Chemical shift deviations of amide protons between pre-fuscimiditide and iso pre-fuscimiditide. d) Overlay of TOCSY (green) and NOESY (blue) spectra for iso pre-fuscimiditide showing connectivity of the D10 sidechain to G11 main chain. NOESY crosspeaks between the β protons of D10 and the amide proton of G11 are observed.

Intrigued by the regioselectivity of fuscimiditide hydrolysis, we sought NMR confirmation that the preferred product had isoAsp in its backbone. In the TOCSY and NOESY spectra (Fig. S21-S22) of the putative isoAsp containing peptide, there was a distinct upfield shift of 0.713 ppm observed for the Asp10 amide proton (Fig. 4c, Table S4). Additionally, strong NOEs were observed between the amide proton of Gly11 and the two distinct β-protons of Asp10 (Fig. 4d). In contrast, a single NOE was observed between the amide proton of Gly11 and the α-proton of Asp10 for the L-Asp pre-fuscimiditide (Fig. S23). These NOEs indicate that the native peptide bond was no longer intact, and it was replaced by a newly formed β-peptidyl linkage. We will refer to this major hydrolysis product of fuscimiditide as iso pre-fuscimiditide to signify the presence of the isoAsp residue in its backbone (Fig. 4b).

### Role of leader peptide in fuscimiditide maturation

Having established the structure of fuscimiditide, pre-fuscimiditide, and the hydrolysis product iso pre-fuscimiditide, we turned our attention to investigating the enzymology of its biosynthesis in more detail. First, we asked whether the leader peptide was required for ThfB activity. In both lanthipeptides and lasso peptides, N-terminal portions of the leader peptide can be removed without affecting modification of the resulting peptide.^54, 55^ In several other RiPPs such as cyanobactins, lanthipeptides, and microviridins, the leader peptide can be supplied in *trans* or can be fused to the N-terminus of the modifying enzymes.^56–58^ Since the fuscimiditide precursor ThfA was observed to be a random coil, we reasoned that large N-terminal truncations of the leader peptide may be tolerated without having an effect on peptide cyclization. To test the role of the lengthy (67 aa, without the added His tag) leader and to identify potential key residues, a series of truncation variants were created by removing 10 amino acid segments from the N-terminus (Fig. S24). Removal of more than 20 N-terminal residues abolished modification by the ATP-grasp ThfB. The intolerance of truncations larger than 20 residues indicated that much of the leader peptide was necessary for installation of the ester linkages by ThfB.

To further probe the importance of the leader peptide, we searched it for conserved motifs. In microviridins, the highly conserved PFFARFL α-helical motif at the N-terminus is key in ATP-grasp activation.^59^ We generated a list of ThfA homologs by BLAST (Fig. S25a), and upon alignment identified a conserved region, RPFG at positions 25-28 of ThfA (Fig. S25b), at the beginning of the minimal leader peptide. We tested the significance of the RPFG motif by executing an alanine scan. However, all variants were successfully modified by both ATP-grasp ThfB and *O*-methyltransferase ThfM (Fig. S25c).

### *In vitro* reconstitution of fuscimiditide biosynthesis

We next turned our attention to *in vitro* reconstitution of fuscimiditide biosynthesis. We purified unmodified N-terminal 6xHis tagged ThfA precursor both by native and denaturing methods. Following the strict order of reaction observed in heterologous expression experiments (Fig. 2a), 10 μM ThfA was first incubated with 1 μM ThfB in the presence of a 1000-fold excess of ATP, MgCl_2_, and DTT in 100 mM sodium phosphate buffer, pH 6.8 at 37 °C. DTT was used to prevent dimerization of ThfA (Fig. S26), which contains one Cys residue in its leader region. The reaction was monitored every hour by LC-MS; after 5 h nearly all the starting material was consumed, and the major product was doubly dehydrated ThfA (Fig. 5a). Overnight incubation of the reaction mixture did not yield an appreciable change in the extent of reaction. ThfA purified under native and denaturing conditions performed equally well in these reactions. These results indicate that ThfB can modify ThfA *in vitro*, albeit at a relatively low turnover rate which could be due to tight binding of the enzyme to the matured product. Additionally, it was previously reported that the reaction efficiency of ATP-grasp catalyzed reactions can be increased ~20-fold in the presence of monovalent cations such as K^+^.^60^ We repeated the reaction in the presence of 100 mM KCl and monitored the formation of the 2-fold dehydrated product every 10 mins; the reaction was ~86% complete in 1 hr (Fig. S27). Under both reaction conditions (with and without KCl) we observed minimal singly dehydrated ThfA (Fig. 5a), suggesting that once one ester forms, the second one forms rapidly. This result is in contrast to the proposed distributive mechanism for ester installation in plesiocin and thuringinin.^36,38^

**Fig. 5.**
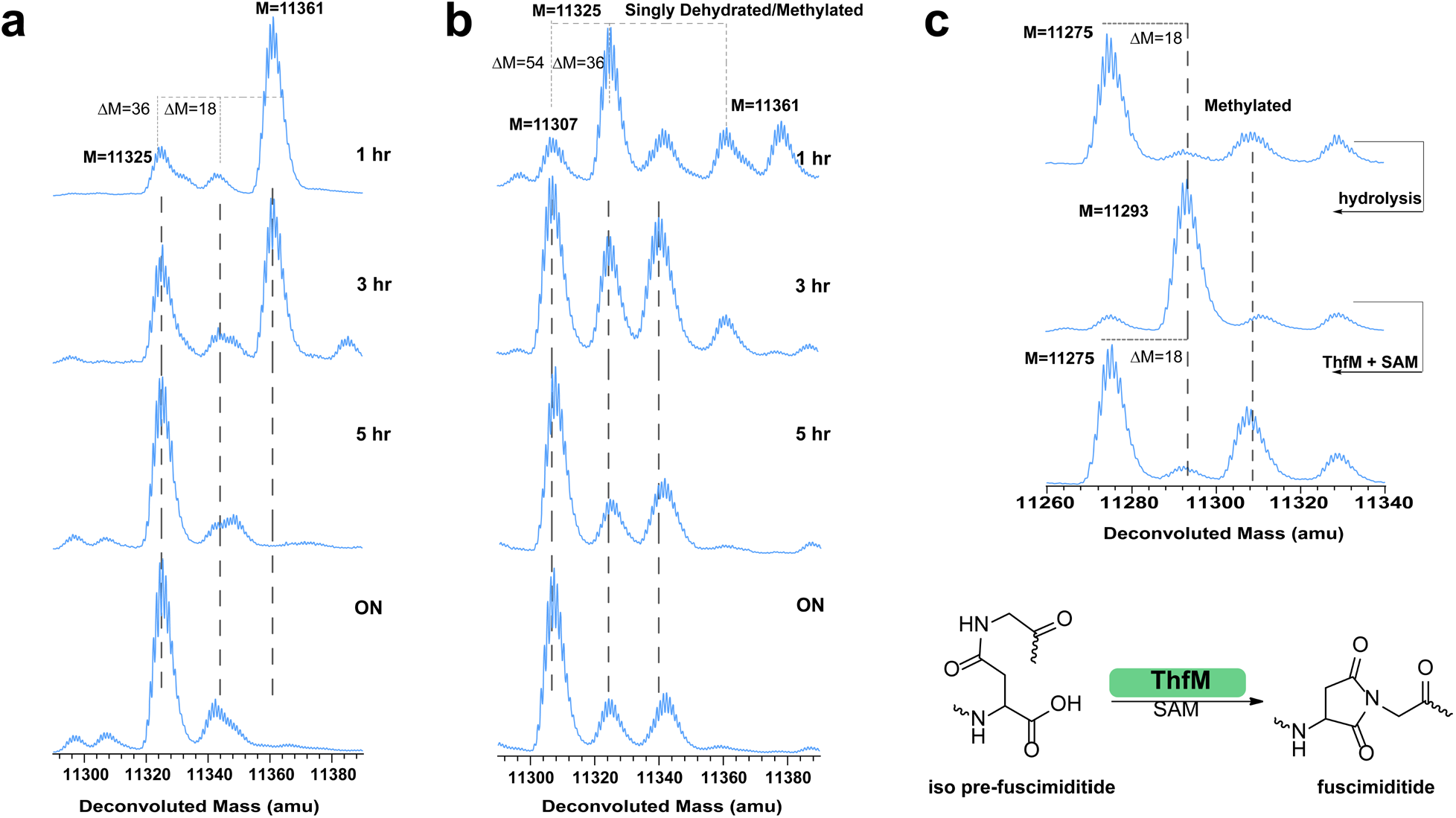
Analysis of ThfB and ThfM activity *in vitro*. a) Time course of the reaction between ThfA and ThfB. Unmodified ThfA (M = 11361) is converted primarily to the doubly esterified product over the course of 5 h. The single ester product does not accumulate, suggesting a model in which the formation of the first ester enables the formation of the second ester. b) Time course of the reaction of ThfA with ThfB and ThfM. A triply dehydrated product is the major species after 5 h indicating that ThfM can facilitate aspartimide formation *in vitro*. The isotopic patterns of singly esterified ThfA and methylated, doubly esterified ThfA overlap. c) Triply dehydrated ThfA C62A (top) was hydrolyzed to its doubly esterified form, comprised primarily of the isoAsp hydrolysate (see Fig. 4). Upon treatment with ThfM and SAM, this protein was dehydrated again to its fully modified form. Some putatively methylated protein is observed as well. These results show that doubly esterified ThfA with isoAsp within its loop macrocycle is a substrate for ThfM (bottom).

We further probed the reconstitution of the full biosynthetic pathway by adding *O*-methyltransferase ThfM (1 μM) simultaneously with ATP-grasp ThfB in the abovementioned reaction mixture (lacking KCl) in addition to SAM (0.4 mM, 40-fold excess relative to ThfA). Conversion to ~75% triply dehydrated product was observed in 5 h with minimal difference upon overnight incubation of the reaction (Fig. 5b). We also set up *in vitro* reactions to test whether ThfM alone could function or if it needed ThfB in the reaction. In accordance with our heterologous expression experiments, a reaction that included unmodified ThfA (10 μM) and ThfM (1 μM) with SAM (0.4 mM) led to no reaction. We also tested whether doubly dehydrated ThfA was a substrate for ThfM. Under native protein purification conditions, however, doubly dehydrated His-tagged ThfA co-elutes with a roughly equimolar amount of untagged ThfB (Fig. S26a) Therefore, we repurified doubly dehydrated ThfA with an additional denaturing step to isolate it from ThfB (Fig. S26b). When doubly dehydrated ThfA was incubated with ThfM in the presence and absence of SAM, formation of the 3-fold dehydrated product was strictly dependent on the presence of SAM and proceed to ~50% completion within 1 h (Fig. S28). This experiment shows that the ThfM enzyme can function on its own, as long as it provided a doubly dehydrated ThfA substrate, and that ThfM acts more rapidly than does ThfB.

Heterologous expression experiments demonstrated that most of the leader peptide of ThfA was needed for the ATP-grasp ThfB to function, while the *O*-methyltransferase ThfM only functioned on the modified precursor. With a functioning *in vitro* system, we could ask whether the leader peptide was required for the modification of the core peptide by ThfM. This experiment is not possible with heterologous expression due to the requirement for the leader peptide in the ThfB reaction. Reactions were set up with pre-fuscimiditide (ie doubly dehydrated core peptide without a leader, 10 μM), ThfM (1 μM) and SAM (0.4 mM) in the same buffer described above. No additional dehydration was observed on pre-fuscimiditide in this reaction, suggesting that at least some portion of the leader peptide is required for the proper function of ThfM (Fig. S29).

Further probing ThfM activity *in vitro*, we attempted modification of ThfA with a D77A substitution. This position corresponds to Asp10 of fuscimiditide, the site of aspartimidylation. We set up a reaction with ThfA D77A, ThfB, and ThfM with ATP and SAM in the same concentrations used above. As expected, overnight incubation of this reaction led to a product with only two dehydrations, further validating Asp10 as the site of aspartimidylation (Fig. S30). Additionally, we noted above that fuscimiditide can hydrolyze into two distinct forms (Fig. 4a) with either Asp or isoAsp in the backbone. To explore the possibility of dual substrate recognition by ThfM, we hydrolyzed 3-fold dehydrated ThfA precursor to yield a mixture of full-length precursors corresponding to pre-fuscimiditide and iso pre-fuscimiditide. A C62A variant of ThfA incapable of disulfide bond formation (discussed below in more detail) was used for these experiments. Following hydrolysis, we incubated ThfM with this mixture under the abovementioned conditions. Intriguingly, overnight incubation yielded >90% conversion to the 3-fold dehydrated product (Fig. 5c). This result suggests the exciting possibility that ThfM is a promiscuous PIMT homolog that can drive both hydrolysis products, containing either Asp or isoAsp, back to the aspartimide.

Under the conditions established for the *in vitro* reconstitution of fuscimiditide, the rate of the ATP-grasp catalyzed reaction was fairly slow; it took several hours for ThfB to execute ~10 turnovers of ThfA (Fig. 5a). As discussed above, heterologous expression of doubly dehydrated ThfA purified under native conditions pulled down an equimolar amount of untagged ATP-grasp ThfB. Our initial thought was that this pulldown could be caused by disulfide bond formation between a Cys residue in ThfB with a Cys residue in the precursor ThfA. Therefore, we generated a C62A substitution in ThfA. However, ATP-grasp pulldown was still observed in the absence of Cys (Fig. S31). This type of association between a RiPP product and its maturation enzyme has been observed before between microcin B17 and its maturation machinery.^61^ Ultimately, we were able to separate the 2-fold dehydrated precursor from ThfB by repurifying the complexes in the presence of urea (Fig. S26b). Having the C62A ThfA variant allowed us to probe its interaction with the ATP-grasp ThfB via biolayer interferometry (BLI). To facilitate this experiment, a maltose binding protein (MBP) fusion to ThfB was purified to use as the analyte in the BLI experiments (Fig. S32). Upon testing the unmodified and doubly dehydrated versions of C62A ThfA, strong affinity to ATP-grasp ThfB was observed for both versions. The K_d_ value for unmodified ThfA binding to ThfB was 28 nM, while doubly dehydrated ThfA exhibited a similar value of 18 nM indicating strong affinity to the substrate as well as the matured product (Fig. S33). This tight binding between the enzyme and its product likely accounts for the low turnover numbers observed in the *in vitro* enzymology experiments described above.

## Discussion

Here, we have described the discovery, biosynthesis, and NMR structure of fuscimiditide, a stem-loop bicyclic peptide and a novel example of microviridin/OEP-like RiPPs. The fuscimiditide BGC includes an *O*-methyltransferase gene that installs a stable aspartimide residue within the loop macrocycle. Though we have focused on this single example of a novel aspartmide-modified OEP, bioinformatic analyses reveal that this class of RiPP BGCs is widespread, though primarily found in actinobacteria. Fig. S34 shows a sampling of 50 putative precursors with homology to the fuscimiditide precursor ThfA. The BGC for each of these precursors also includes ATP-grasp and *O*-methyltransferase enzymes. These precursors are all enriched in Ser, Thr, Asp, and Glu residues near their C-termini, and all of them have an Asp residue that is the putative site for aspartimidylation. Of interest is that this Asp residue is mostly followed by either Gly or Ser. The architecture of these BGCs is also well-conserved with all examples exhibiting an A-B-M gene order except for fuscimiditde, in which the gene order is B-A-M as discussed above.

The aspartimide installed by *O*-methyltransferase ThfM differentiates fuscimiditide from other characterized microviridins/OEPs. Normally aspartimide functions as either an intermediate in protein repair or as a nuisance product in solid-phase peptide synthesis or protein pharmaceutical formulation.^42,43,62^ Our data strongly supports the idea that, at least *in cellulo*, the aspartimidylated product fuscimiditide, is the final, intended product of the BGC. The three PTMs of fuscimiditide (two esters and one aspartimide) are all confined within the final 20 aa of the precursor protein ThfA, so it is logical to think that there may be a protease that removes the leader peptide of ThfA and releases the macrocycle. We did not observe any obvious candidates for proteases in the immediate vicinity of the fuscimiditide BGC, so it seems unlikely that there is a dedicated protease for leader removal. Recent work on class IV lanthipeptides has shown that multiple proteases not colocalized with the BGC can function in leader peptide removal.^63,64^ In general, the proteolysis of leader peptides is understudied in the microviridin/OEP field; a C39 peptidase-ABC transporter fusion protein has been hypothesized to serve this role for microviridins.^18,31,59^ If a similar system for leader peptide cleavage and secretion of fuscimiditide is present in the *T. fusca* genome, the final state of its aspartimide residue will be dictated by the environment into which it is secreted. While we expect the aspartimide to be long-lived in acidic or neutral conditions, it will rapidly hydrolyze to the isoAsp form, iso pre-fuscimitide, in basic conditions. Another intriguing possibility is that posttranslationally modified ThfA remains in the cell. In this case, even if the aspartimide hydrolyzes, it will be dehydrated again due to the promiscuous activity of ThfM.

While this paper has focused on detailed characterization of the structure and biosynthesis of fuscimiditide, we have not studied its bioactivity in great depth. One possible speculation about the bioactivity of fuscimiditide and other aspartimide-modified OEPs is based on the observation that many members of this class serve as protease inhibitors.^32,37^ The addition of an electrophilic moiety like the succinimide to such inhibitors could convert them into suicide inhibitors that covalently attach to a protease target. In addition to OEPs as described here, this novel class of Asp-modifying *O-*methyltransferases has also been found in lasso peptide BGCs (see companion paper by Cao et al.)^47^ as well as in lanthipeptide BGCs.^65^ While all three examples of these RiPP modifying methyltransferases start off with Asp methylation and formation of aspartimide, the fate of the aspartimide residue differs. In the lanthipeptide case, the aspartimide moiety hydrolyzes rapidly to give a mixture of Asp and isoAsp residues in the lanthipeptide backbone. For the fuscimiditide case described here and the lasso peptide case, the macrocycle-embedded aspartimide moieties are more stable. When fuscimiditide hydrolyzes, it does so in a regioselective fashion yielding ~95% of the isoAsp backbone peptide. The opposite result is observed for the lasso peptides cellulonodin-2 and lihuanodin, both of which open exclusively to the Asp backbone form. Taken together, these works demonstrate a novel function for the well-studied PIMT family of enzymes. This work also suggests further studies into the stability, bioactivity, and structural implications of aspartimide residues found in macrocycles.

## Methods

### Expression and purification of ThfA, ThfB, and ThfM

His-tagged versions of ThfA, ThfB, and ThfM were generated using standard recombinant DNA techniques (see SI Methods for full details). Non His-tagged variants of ThfB and ThfM were also generated for coexpression with His-tagged ThfA (see SI Methods for cloning details). All proteins (ThfA coexpressed with ThfB, ThfM, or both, as well as ThfA, ThfB, and ThfM alone) were expressed in *E. coli* BL21(DE3) Δ*slyD. E. coli* harboring the plasmid of interest were cultured at 37 °C until the culture reached an OD_600_ of 0.5 at which point the culture was induced with 1 mM IPTG. The culture was further incubated at 37 °C for 4 h. The cells were collected by centrifugation, lysed by sonication, and His-tagged proteins were purified using Ni-NTA resin under either native or denaturing conditions (see SI methods for details). Purified proteins were desalted into phosphate buffered saline (PBS) with 10% glycerol and frozen at −80 °C until needed.

### HPLC and mass spectrometry

An HPLC system (Agilent 1200 series) was used for purification of fuscimiditide, pre-fuscimiditide, and iso pre-fuscimiditide, the 22 aa products of tryptic digests of modified ThfA. A semi-prep column (Zorbax 300SB-C18, 9.5 mm x 250 mm, 5 μm particle size, Agilent) was used for all purification steps; gradient details are in the SI Methods. For LC-MS analysis, an Agilent 6530 QTOF mass spectrometer was used downstream of an Agilent 1260 HPLC system. Two different analytical columns were used: for analysis of peptides, a Zorbax 300SB-C18, 2.1 mm x 50 mm, 3.5 μm particle size (Agilent) was used. For analysis of intact proteins such as ThfA and its modified variants, a XBridge Protein BEH C4 (2.1 mm x 50 mm, 3.5 μm particle size, Waters) was used. Mass spectra were analyzed using MassHunter Bioconfirm software.

### NMR and structure determination

NMR spectra were collected on a Bruker Avance III 800 MHz spectrometer. TOCSY spectra were acquired at 80 ms mixing time while NOESY spectra were acquired at both 150 ms and 700 ms mixing times. Proton-carbon HSQC and HMBC experiments were carried out at natural carbon abundance. The HMBC experiments were optimized at both 5 Hz and 10 Hz coupling constants. All spectra were assigned manually and integrated using MestReNova software; the 150 ms NOESY spectra were used to generate distance restraints. CYANA 2.1 was used to calculate 20 structural models. The structures were energy-minimized using Avogadro. The structural ensembles of fuscimiditide and pre-fuscimiditide have been deposited (PDB codes 7LIF and 7LI2, respectively).

### Reconstitution of fuscimiditide biosynthesis *in vitro*

Purified His-tagged ThfA, ThfB, and ThfM were used directly in *in vitro* assays. Assays with ThfA and ThfB were set up as follows: 10 μM ThfA, 1 μM ThfB, 10 mM ATP, 10 mM MgCl_2_, 10 mM DTT in 100 mM sodium phosphate buffer. Assays with ThfA, ThfB, and ThfM contained the above plus 1 μM ThfM and 400 μM SAM. In addition, assays were carried out with ThfA/modified ThfA and ThfM. These assays contained 10 μM ThfA/modified ThfA, 1 μM ThfM, 400 μM SAM, 10 mM MgCl_2_, 10 mM DTT in 100 mM sodium phosphate buffer. Reactions were carried out on a 50 μL scale in PCR tubes and incubated at 37 °C in a thermocycler. For time course analysis, a 10 μL was removed and quenched by the addition 1 μL of 10% formic acid. These samples were directly analyzed by LC-MS.

## Acknowledgments

We thank I. Pelczer (Princeton University NMR Facility) for help with acquiring NMR spectra and Hendrik Schroeder for assistance with tandem mass spectrometry. This work was supported by National Institutes of Health Grant GM107036 and a grant from Princeton University School of Engineering and Applied Sciences (Focused Research Team on Precision Antibiotics). L.C. was supported by an NSF Graduate Research Fellowship Program under Grant DGE-1656466. J.D.K. was supported in part by training grant T32 GM7388.

## Author Contributions

H.E.E., J.D.K., and A.J.L. conceived of the project. H.E.E, J.D.K., W.L.C-L., B.C., L.C., M.A.R., and H.L.W. carried out experiments. H.E.E. and A.J.L wrote the initial draft of the paper, all authors participated in revision of the paper. A.J.L. acquired funding and supervised the project.

## Data Availability Statement

The coordinates for fuscimiditide and pre-fuscimiditide has been deposited in the Protein Data Bank (PDB) with accession codes 7LIF and 7LI2, respectively. Coordinates has also been deposited to the Biological Magnetic Resonance Bank (BMRB); the accession numbers are 30851 for fuscimiditide and 30849 for pre-fuscimiditide. Raw mass spectrometry data underlying figure will be provided upon request. All other data is present in the main text or the supplementary information.

