## Supplementary Information for "Fuscimiditide: a RiPP with Ω-Ester and Aspartimide Post-translational Modifications"

**Table of Contents**

Table of Contents ········ 2
Supplementary Methods and Materials ........ 3
Supplementary Figures ········ 8
Supplementary Tables ········ 46
References for Supplemental Information ········ 61

### Methods and Materials

#### Strains and Reagents

*Thermobifida fusca* (DSM 43792) was obtained from the German Collection of Microorganisms and Cell Cultures (DSMZ). Genomic DNA (gDNA) was isolated utilizing the Qiagen DNeasy Blood and Tissue Kit standard protocol. PCR amplification was performed using Q5 polymerase purchased from New England Biolabs (NEB) and all oligonucleotides were purchased from Integrated DNA Technologies (IDT). All primers used in this study are listed in Table S1. Restriction enzymes and T4 DNA ligase were purchased from New England Biolabs. XL-1 Blue *E. coli* were used for all recombinant DNA procedures. Synthetic linear core peptide was purchased from GenScript and re-purified by HPLC. All sequencing results are confirmed by Sanger sequencing (Genewiz).

#### Plasmid Construction

The genes from the fuscimiditide cluster were individually cloned from genomic DNA. The gene for the fuscimiditide precursor *thfA* was amplified using primers 1 and 2, digested with *Bam*HI and *Hin*dIII, and ligated into the multiple cloning site MCS1 of pRSF Duet. All 6xHis tagged precursor mutants and truncated versions were amplified with their respective primers from Table S1, digested with *Bam*HI and *Hin*dIII, and ligated into MCS1 of pRSF Duet. This resulted in a N-terminally 6xHis tagged version of the native, mutant, or truncated precursor. Both an untagged and His-tagged version of the ATP-grasp gene *thfB* were created. Using primers 3 and 5, the gene was amplified from gDNA, digested with *Eco*RI and *Hin*dIII, and ligated into pQE80 to create the untagged version of ThfB. Amplification with primers 4 and 5 followed by digestion with *Bam*HI and *Hin*dIII and ligation into pQE80 yielded the N-terminally 6xHis tagged version of the ThfB. Primer 3 introduces a ribosome binding site (AGGAGA) 10 bases upstream of the start codon. Methyltransferase ThfM was also amplified twice to create both an N-terminally 6xHis tagged and untagged protein. The N-terminally 6xHis tagged version was constructed by amplifying the *thfM* gene from gDNA using primers 6 and 7, digestion with *Bam*HI and *Hin*dIII, and ligation into pQE80. Amplification with primers and digestion with *Eco*RI and *Hin*dIII yielded the untagged version in pQE80. Golden Gate Cloning was used to generate a bicistronic construct in pQE80 that contained untagged versions of both the *thfB* and *thfM* genes. Briefly, cassettes with each gene and a ribosome binding site upstream of the gene were designed with flanking, non-complementary *Bsa*I sites. These cassettes were digested with *Bsa*I and ligated simultaneously into pQE80 digested with *Eco*RI and *Hin*dIII.

#### Expression and Native Purification of ThfA and Modified ThfA

N-terminally His-tagged ThfA, ThfB, and ThfM as well as ThfA coexpressed with ThfB and/or ThfM were all purified in a similar fashion. The plasmid(s) containing the gene(s) of interest were transformed into *E. coli* BL21 Δ*slyD* cells and grown up overnight in Luria-Bertani (LB) + 50 mg/L kanamycin for pRSF Duet plasmids and 100 mg/L ampicillin for pQE80 plasmids. The cells were subcultured at an $OD_{600}$ = 0.02 into LB kanamycin/ampicillin as needed and allowed to grow at 37 °C. Once the $OD_{600}$= 0.5, expression was induced using 1 mM IPTG. The cells were grown for 4 hours at 37 °C before being spun down at 4000 x *g* for 15 min. The pellet was resuspended in 50 mM $NaH_2PO_4$, 300 mM NaCl, 10 mM imidazole, pH 8.0 before freezing at -80 °C. For purification, the resuspended cell pellet was thawed in ice water and 1 mg/mL lysozyme was added, followed by incubation at 4 °C for 20 minutes. The sample was then sonicated on ice for 12 cycles of 10 s on and 20 s off to lyse the cells and spun down at 4000 x *g* for 15 minutes at 4 °C. The supernatant was removed and spun down an additional time at 8000 x *g* for 10 minutes. The clarified lysate was then incubated with 1 mL of Ni-NTA resin (Qiagen) per liter of culture while rotating for 1 hour at 4 °C. The mixture was added to an empty gravity column, and the flow-through was passed over the resin an additional time. The resin was then washed with 10 mL of 50 mM $NaH_2PO_4$, 300 mM NaCl, 20 mM imidazole pH 8.0 and twice with 10 mL of 50 mM $NaH_2PO_4$, 300 mM NaCl, 50 mM imidazole pH 8.0. The protein was then eluted with 50 mM $NaH_2PO_4$, 300 mM NaCl, 250 mM imidazole pH 8.0. The samples were run on SDS-PAGE and the most concentrated samples were pooled and buffer exchanged using a PD-10 desalting column (Biorad) into 1X PBS pH 7.4 containing 10% glycerol. ThfA and ThfA coexpressed with ThfB and/or ThfM was then concentrated using a 3 kDa Amicon concentrator and frozen at -80 °C until needed. His-tagged ThfB and ThfM were concentrated with a 10 kDa Amicon concentrator and frozen at -80 °C until needed. These proteins were used for *in vitro* studies of fuscimiditide maturation.

**Denaturing Purification of Modified ThfA**

His-tagged ThfA coexpressed with ThfB or ThfA coexpressed with both ThfB and ThfM were purified under denaturing conditions. Protein expression was carried out as described in the previous paragraph. The cell pellets were resuspended in 100 mM $NaH_2PO_4$, 10 mM Tris, 8 M urea, pH 8.0 (Buffer B) and stored at -80 °C. For purification, the cells were thawed and spun down at 4000 x *g* for 30 minutes at 4 °C. The supernatant was removed and spun down again at 8000 x *g* for 20 minutes. The clarified supernatant was mixed with 1 mL of Ni-NTA resin per liter of culture and incubated while rotating for 1 hour. The mixture was then added to an empty gravity column and allowed to flow through. The resin was washed with 10 mL of the lysis buffer adjusted to pH 6.3 (Buffer C) and pH 5.9 (Buffer D), before the final elution of 1 mL fractions at pH 4.5. The samples were run on SDS-PAGE and the most concentrated fractions were pooled and buffer exchanged using a PD-10 column and concentrated using a 3 kDa Amicon concentrator. These proteins were used for mass spectrometry analysis as well as substrates for trypsin digestion.

**Trypsin Digestion**

Tryptic digests of doubly and triply dehydrated ThfA were used both on an analytical scale mass spectrometry analysis and on a preparative scale (i.e. multiple mg) to generate samples of pre-fuscimiditide, fuscimiditide, and iso pre-fuscimiditide for NMR. For small scale analysis, purified ThfA or modified ThfA and sequence grade Trypsin (Promega) was mixed at a 1:100 w/w protease:protein ratio in 50 mM $NH_4CO_3$ buffer at 37 ºC for 20 hours before being quenched by addition of 1% formic acid. To generate the pre-fuscimiditide sample, the urea eluent from denaturing purification of 16 L of culture (see Denaturing Purification of Modified ThfA above) was diluted with 50 mM $NH_4CO_3$ buffer until the urea concentration was less than 1M. Trypsin was added to a final protease:protein ratio of 1:200 w/w and the sample was incubated at 37 °C for 20 hours before being stopped by addition of formic acid to a final concentration of 1%. The sample was lyophilized and the resulting powder stored at -80 °C until purification by HPLC. The fuscimiditide NMR sample was generated in a similar fashion, but with some important changes due to the lability of the aspartimide moiety. The urea eluent from denaturing purification (see previous section) was immediately buffer exchanged using a PD-10 column into 50 mM ammonium bicarbonate, adjusted to pH 7 using HCl. Trypsin was added at a 1:200 protease:protein mass ratio as above, but the digestion proceeded for only 1 hr at 37 °C. At this time the reaction was quenched with formic acid to a final concentration of 1%. The sample was immediately frozen, lyophilized, and stored until HPLC purification.

**HPLC**

Semi-preparative reverse-phase HPLC was performed using an Agilent 1200 series instrument equipped with a Zorbax 300SB-C18 (9.4 mm x 250 mm, 5 μm) column and a 215 nm UV detector. The elution gradient was run at a flow rate of 4 mL/min. The mobile phase A contained water and 0.1% trifluoroacetic acid and phase B contained acetonitrile with 0.1% trifluoroacetic acid. The gradient was as follows: 10% acetonitrile for 1 min, 10-50% acetonitrile over 19 min, 50-90% acetonitrile over 5 min, 90% acetonitrile for 5 min, and 90-10% acetonitrile over 2 min. Peaks at 8.82 min, 9.23 min, and 8.93 min were collected for pre-fuscimiditide, fuscimiditide, and iso pre-fuscimiditide respectively. Purified fractions were pooled, frozen at -80 °C for several hours, and then lyophilized (Labconco FreeZone Freeze Dry System). The peptide was resuspended in $H_2O$ or buffer as needed.

**Mass Spectrometry**

All mass spectra were collected using electrospray ionization (ESI) on an Agilent 6530 QTOF equipped with an Agilent 1260 LC system. Peptides from digested proteins were chromatographed on a Zorbax 300SB-C18 (2.1 mm x 50 mm, 3.5 μm particle size). Intact proteins were chromatographed on an XBridge Protein BEH C4 (Waters, 2.1 mm x 50 mm, 3.5 μm particle size). The instrument was calibrated using "Mass Calibration/Check" daily. Data were analyzed using MassHunter (Agilent).

**Hydrazine Reaction**

Hydrazine addition to fuscimiditide was carried to confirm the presence of aspartimide. In the reaction mixture, fuscimiditide in ultrapure water was added to a final concentration of 0.1 mM and hydrazine (35% solution in water) was added to a final concentration of 2M with a total volume of either 25 or 50 μL. The reaction mixture was incubated at room temperature for 30 mins and analyzed by LC-MS.

**NMR and Structure Determination**

Samples of pre-fuscimiditide (ie doubly dehydrated ThfA core peptide, 1.7 mM), fuscimiditide (triply dehydrated ThfA core peptide, 3.3 mM), and iso-pre fuscimiditide (the major product of fuscimiditide hydrolysis, 1.5 mM) were prepared 98/2 $H_2O/D_2O$. Peptide concentrations were determined using HPLC absorbance at 215 nm. 2D NMR spectra (TOCSY, NOESY, $^1H$-$^{13}C$ HSQC, $^1H$-$^{13}C$ HMBC) of pre-fuscimiditide and fuscimiditide were collected on a Bruker Avance III 800 MHz spectrometer at 295 K. The mixing time for the TOCSY was 80 ms while two NOESY spectra were acquired with 150 ms and 700 ms mixing times. For pre-fuscimiditide, two HMBC spectra were acquired targeting 5 Hz and 10 Hz proton-carbon coupling. For fuscimiditide, only a 10 Hz spectrum was acquired. Finally, for iso pre-fuscimiditide, only TOCSY and two NOESY spectra (150 ms and 700 ms mixing times) were acquired. Spectra were processed and analyzed using MestReNova. Proton and carbon resonances were manually assigned to the structure (Table S2). The 150 ms mixing time NOESY spectra were used to determine through space distance restraints, which were used to build structural models in CYANA 2.1 for pre-fuscimiditide and fuscimiditide. Additional geometrical restraints corresponding to the Thr-Asp esters (PDB 4KTU)[1] and aspartimide (PDB 1AT5)[2] were added to CYANA 2.1. The top 20 structures for both of these peptides were energy-minimized using Avogadro's MMFF94 forcefield. The coordinates for pre-fuscimiditide (PDB: 7LI2, BMRB: 30849) and fuscimiditide (PDB: 7LIF, BMRB: 30851) have been deposited in the PDB and BMRB.

**Circular Dichroism**

Circular dichroism (CD) studies were carried out on 4 samples: unmodified full-length ThfA (103 aa including the His tag), linear ThfA core peptide (22 aa), pre-fuscimiditide (22 aa, doubly dehydrated ThfA core peptide), and fuscimiditide (22 aa, triply dehydrated ThfA core peptide. Full-length ThfA was expressed recombinantly and purified under native conditions as described above. The unmodified ThfA core peptide was generated by solid-phase peptide synthesis (Genscript) and purified by HPLC. Fuscimiditide and pre-fuscimiditide were prepared in the same fashion as the NMR samples described above (ie recombinant expression, tryptic digestion, and HPLC purification). All samples were dissolved in ultrapure water. The concentration of each sample was estimated using $A_{280}$ measurement on a Nanodrop instrument. The concentrations of ThfA, ThfA core, pre-fuscimiditide, and fuscimiditide were unmodified precursor, modified core, and unmodified core were diluted to 0.1 mg/mL, 0.1 mg/mL, 0.08 mg/mL, and 0.08 mg/mL. CD measurements were collected on an Applied Photophysics Chirascan instrument. The blank (ultrapure water) and samples were each measured three times at 20 °C scanning from 180 nm to 280 nm in a 1 mm pathlength cuvette (Hellma Analytics). The final CD spectrum was obtained by averaging the spectra and subtracting the average background spectrum.

**Binding Assays**

Binding measurements were conducted using biolayer interferometry with the BLItz system and Ni-NTA biosensors from ForteBio. His-tagged ThfA and His-tagged doubly dehydrated ThfA were adsorbed onto the biosensors. A maltose binding protein (MBP)-ThfB fusion protein was generated as the analyte. Briefly, the MBP-ThfB fusion protein was expressed in *E. coli* BL21(DE3) Δ*slyD* and purified using amylose resin (New England Biolabs) according to the manufacturers' suggestions. ThfA and MBP-ThfB samples were buffer exchanged into PBS buffer pH 7.4 and 10% glycerol. Subsequent dilutions were done in 1x Kinetics Buffer composed of 1x PBS pH 7.4 with 0.13% BSA and 0.013% Tween-20. All biosensors were rehydrated with 200 µL 1x Kinetics buffer for at least 10 minutes and were kept hydrated until use; 1 biosensor was used for each measurement. For each measurement, the biosensor was attached to the Blitz system and the following binding kinetics program was used: 40 s baseline measurement where the biosensor was dipped into a fresh tube with 250 µL 1x Kinetics Buffer, 120 s loading of 6xHis precursor where the biosensor was dipped into 4 µL of 19 µg/mL His-tagged ThfA or 18 µg/mL His-tagged doubly dehydrated ThfA on a magnetic drop holder, 40 s baseline measurement where the biosensor was dipped into 1x Kinetics buffer, 120 s binding where the biosensor was dipped into 4 µL of MBP-ThfB on the magnetic drop holder, and a 300 s dissociation step where the biosensor was dipped into the tube with 1x Kinetics Buffer. For each new measurement, a new biosensor and buffer tube was used. Additionally, between measurements, the drop holders were washed 3 times with 1x Kinetics Buffer and dried with a kimwipe. Curves were corrected for the start of association and dissociation, then fitted globally to obtain values for the association rate constant ($k_a$), disassociation rate constant ($k_d$), and the binding affinity $K_d$.

***In vitro* Enzymology of Fuscimiditide Biosynthesis**

For reactions consisting of ThfA and ThfB the following was added to the reaction mixture: 1 μM ThfB, 10 μM ThfA, 10 mM ATP, 10 mM DTT, 10 mM $MgCl_2$ in 100 mM $NaH_2PO_4$ pH 6.87; 50 μL total. For reactions consisting of ThfA, ThfB, and ThfM, the abovementioned conditions were used with the addition of 400 μM SAM and 1 μM ThfM. Reactions with only ThfA and ThfM consisted of 10 μM ThfA, 1 μM ThfM, 400 μM SAM, 10 mM DTT, 10 mM $MgCl_2$ in 100 mM $NaH_2PO_4$ pH 6.87. Finally, the reaction between pre-fuscimiditide and ThfM was set up in a similar fashion: 10 μM pre-fuscimiditide, 1 μM ThfM, 400 μM SAM, 10 mM DTT, 10 mM $MgCl_2$ in 100 mM $NaH_2PO_4$ pH 6.87. All reactions were incubated at 37 ºC in a thermocycler. For time course analysis, 10 μL aliquots were withdrawn, quenched with formic acid (added to a final concentration of 1%) and stored until analysis. The reactions were then analyzed by LC-MS.

**Supplementary Figures**

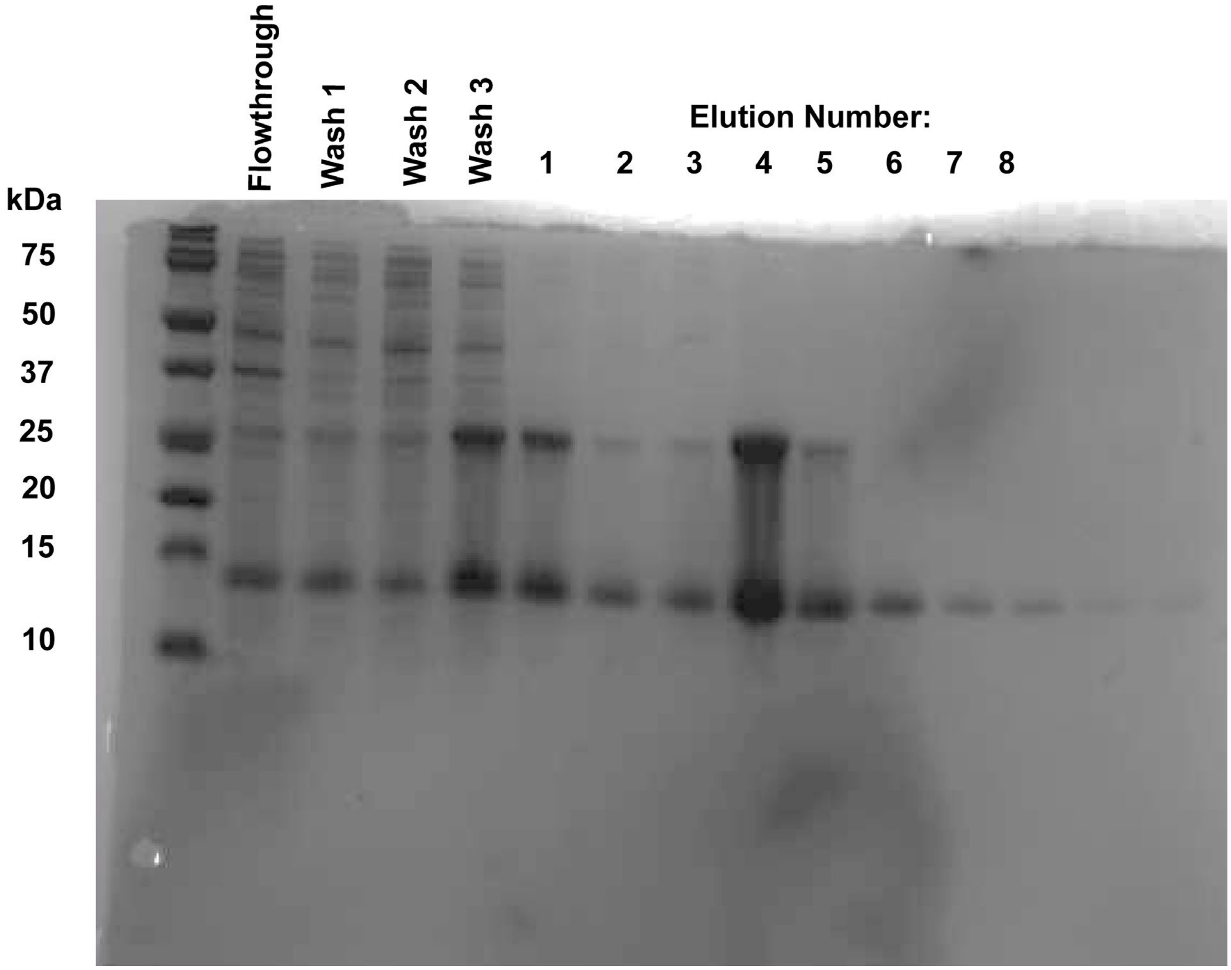


**Fig. S1** SDS-PAGE gel of native Ni-NTA purification of unmodified ThfA precursor protein. Flowthrough, washes, and elutions are labeled. The ThfA sequence includes a single cysteine residue, C62, which leads to dimerization in air, resulting in the bands observed at ~25 kDa.

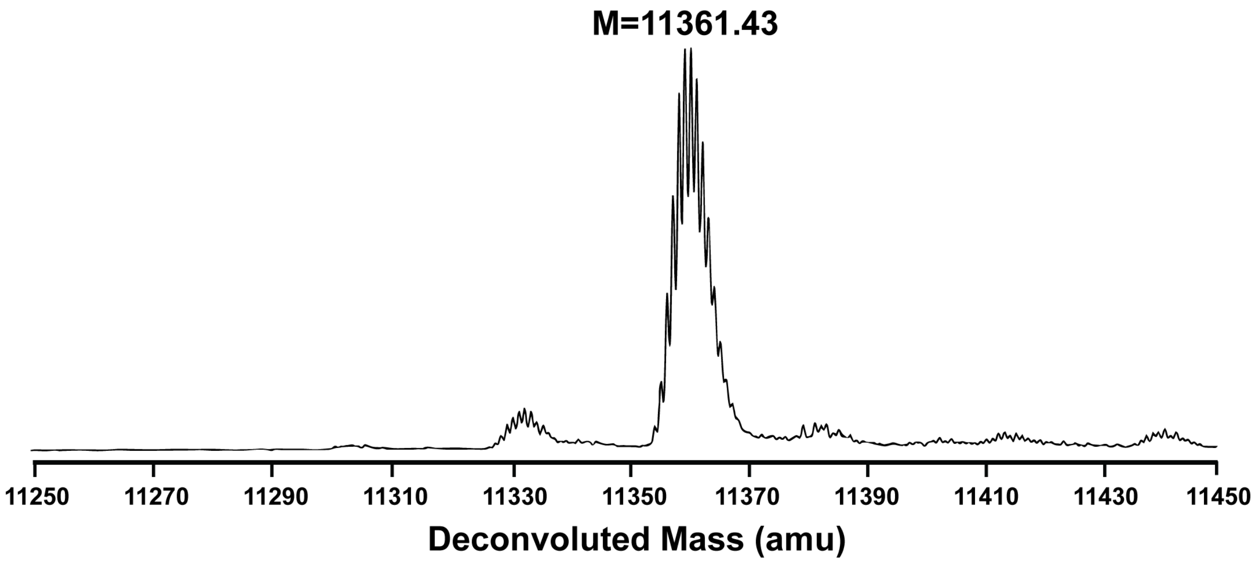


**Fig. S2** Deconvoluted mass spectrum of purified ThfA precursor coexpressed with ThfM methyltransferase. The major peak corresponds to unmodified precursor (refer to Figure 2a), showing that ThfM does not dehydrate this protein.

MGSSHHHHHHSQDPMSTAVTDAFPLGRDENRNDQVTEWRP
FGMRYGVQPTPIPVPLSDTKYDPDQQVLVVADGQPCAKIER
AGTMRVTYPDGQKPGQSDVEKD

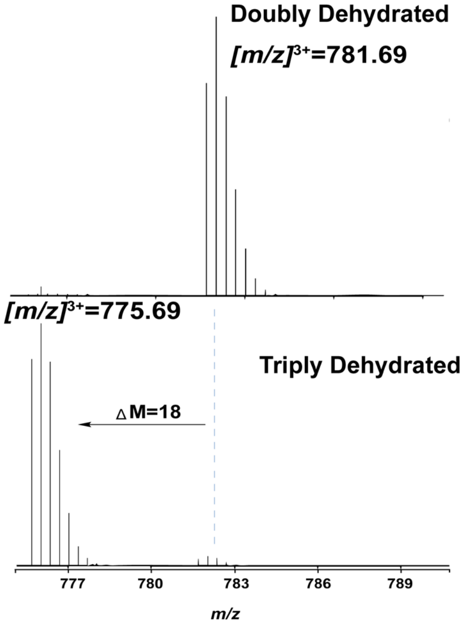


**Fig. S3** Tryptic digest of doubly and triply dehydrated ThfA to identify the location of PTMs. Top: sequence of the ThfA precursor with trypsin cleavage sites underlined. Tryptic digest of doubly- and triply dehydrated ThfA leads to a prominent 22 aa C-terminal fragment with three missed cleavages (sequence in blue), suggesting that the dehydrations are located in the C-terminus. Bottom: Mass spectra of the 22 aa C-terminal fragment arising from trypsin digestion of doubly and triply dehydrated ThfA.

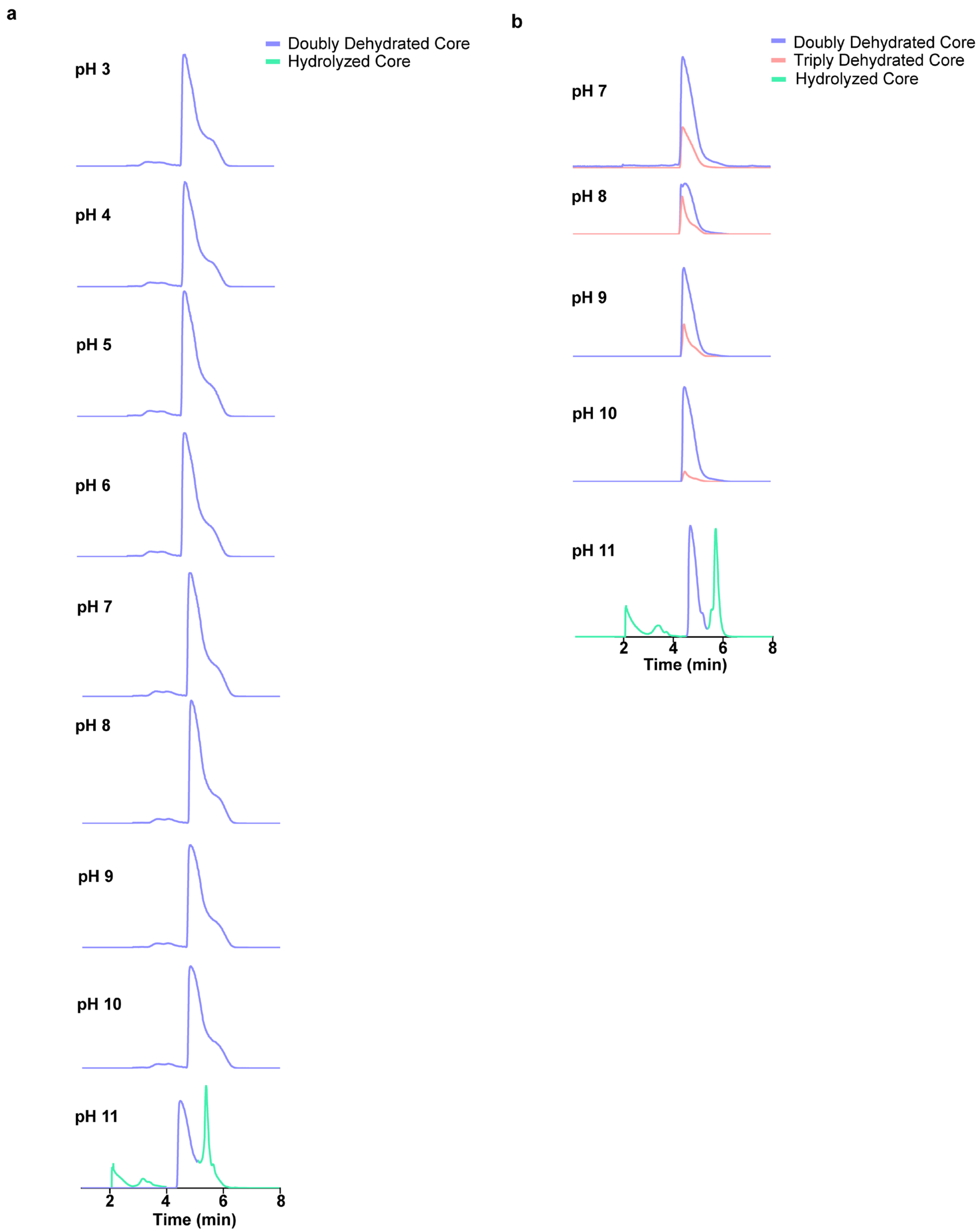


**Fig. S4** Total ion current (TIC) chromatograms of trypsin-digested C-terminal fragment of doubly and triply dehydrated ThfA incubated at a range of pH values. Samples were incubated in phosphate buffered saline, overnight at room temperature. Left: the doubly dehydrated species begins hydrolyzing only at pH 11. Right: the triply dehydrated species hydrolyzes readily to doubly dehydrated species starting at pH 7.

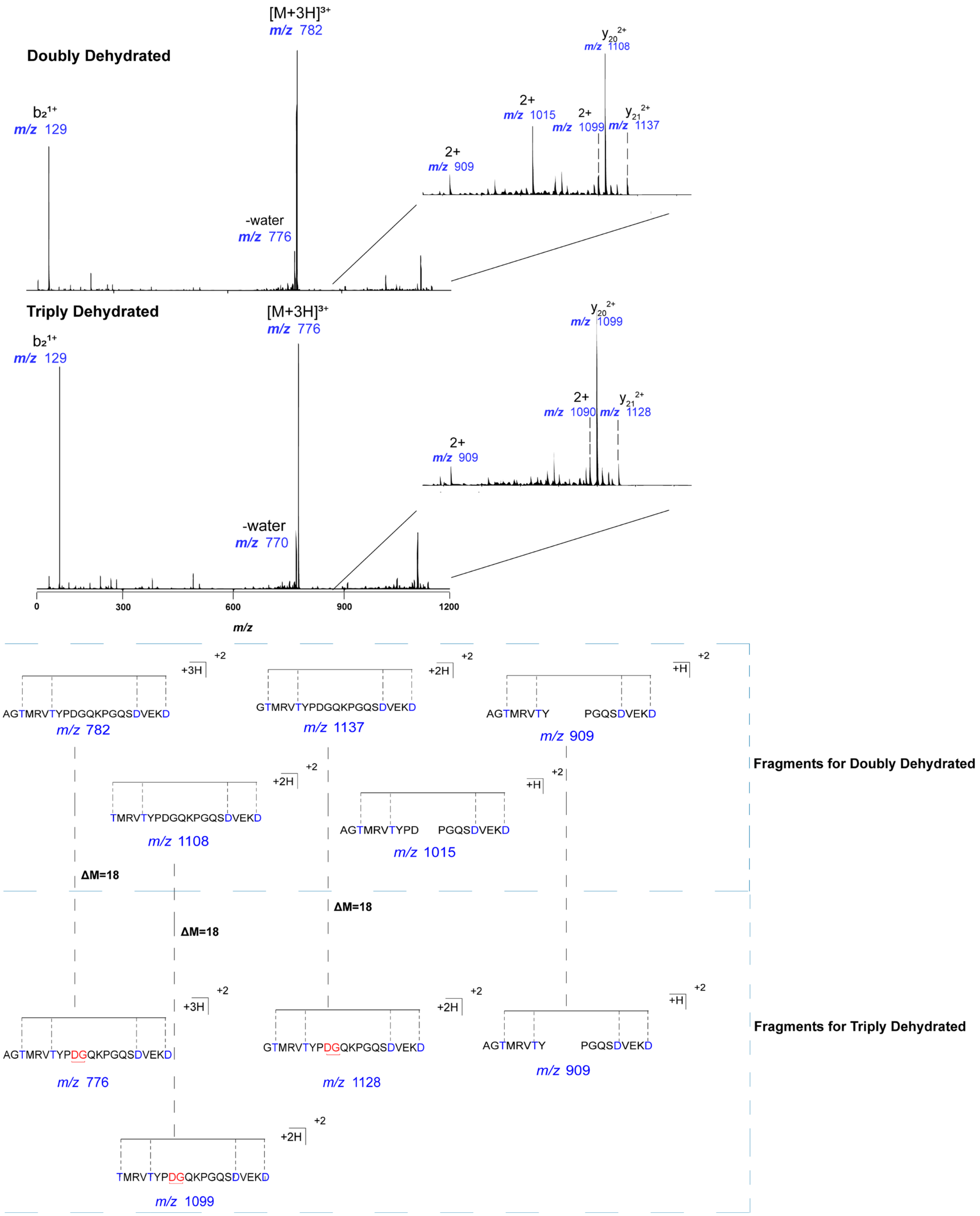


**Fig. S5** Tandem mass spectra of doubly and triply dehydrated core peptides. The major peak after CID fragmentation is still the parent ion ($[M+3H]^{3+}$ = 782 for the doubly dehydrated species and $[M+3H]^{3+}$ = 776 for triply dehydrated). The presence of a $b_2$, $y_{20}$, and $y_{21}$ ions indicate that the peptide is a macrocycle stretching from T3 to D22. Cleavage N-terminal to P9 and P14 was also observed in the CID experiment. Species generated from these cleavages are shown schematically for the doubly dehydrated and triply dehydrated species. The species at *m/z* = 909 corresponds to cleavage at both P9 and P14. This species is present upon fragmentation of both the doubly- and triply dehydrated peptides, showing that the third dehydration is located between these two residues. The ion observed at *m/z* = 1015 for the doubly dehydrated species is not observed in the triply dehydrated species, suggesting modification at the D10 residue.

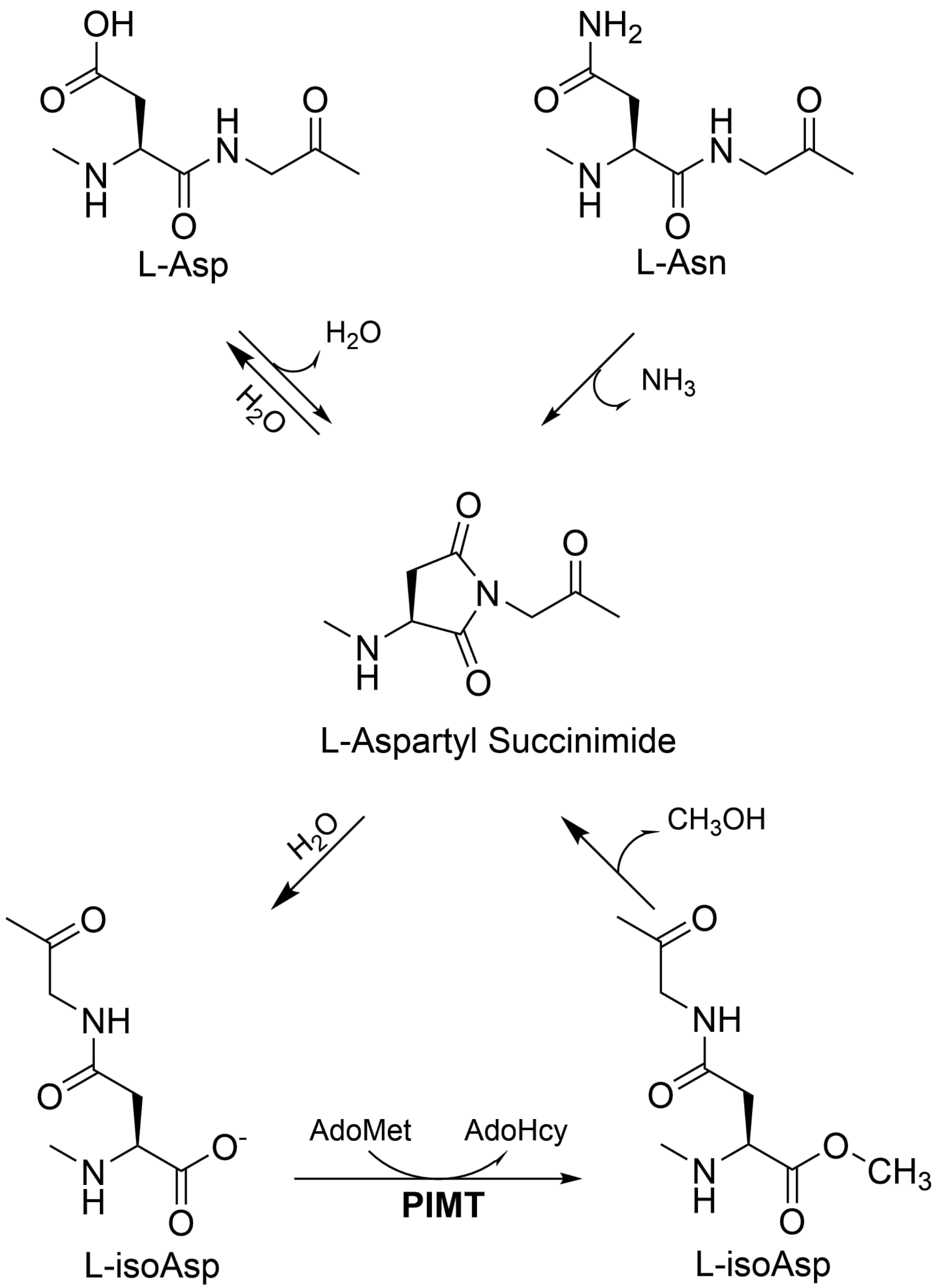


**Fig. S6** Reactions in protein aging and subsequent repair by canonical protein L-isoaspartyl methyltransferase (PIMT). Over time, asparagine (Asn) can spontaneously deamidate and aspartate (Asp) can dehydrate to form an aspartyl succinimide (center). Hydrolysis of the succinimide can generate either Asp or isoaspartate (isoAsp). PIMT specifically recognizes isoAsp over Asp and generates a methyl ester, which is rapidly converted back to the succinimide. Multiple turnovers of PIMT result in restoration of most of the isoAsp residues to Asp.

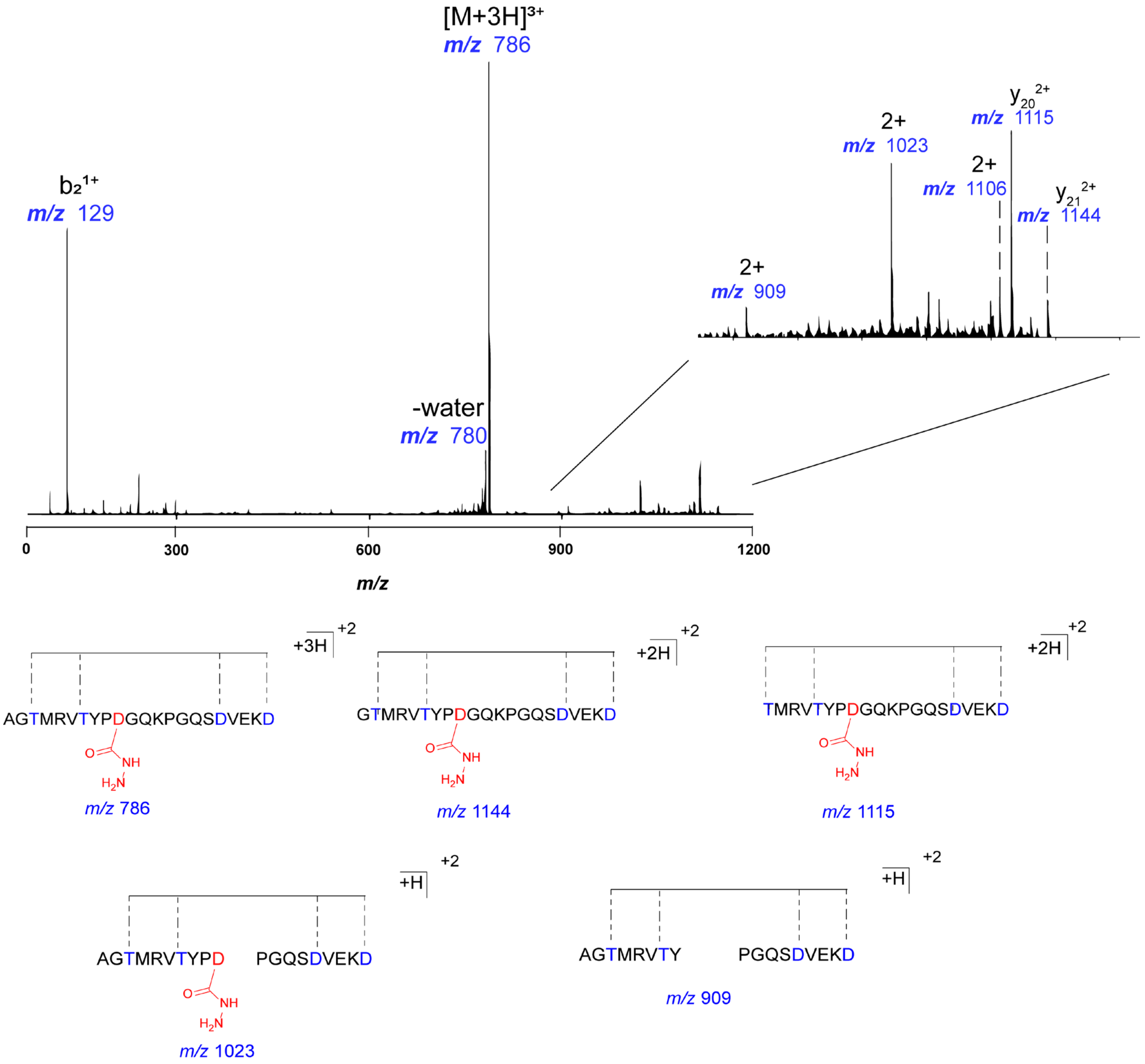


**Fig. S7** MS/MS of hydrazine reacted triply dehydrated core. As in Fig. S5, the major ion upon fragmentation is still the intact peptide. Minor fragmentation products, such as the ions at *m/z* = 909 and *m/z* = 1023 pinpoint the location of hydrazine addition to the D10 residue.

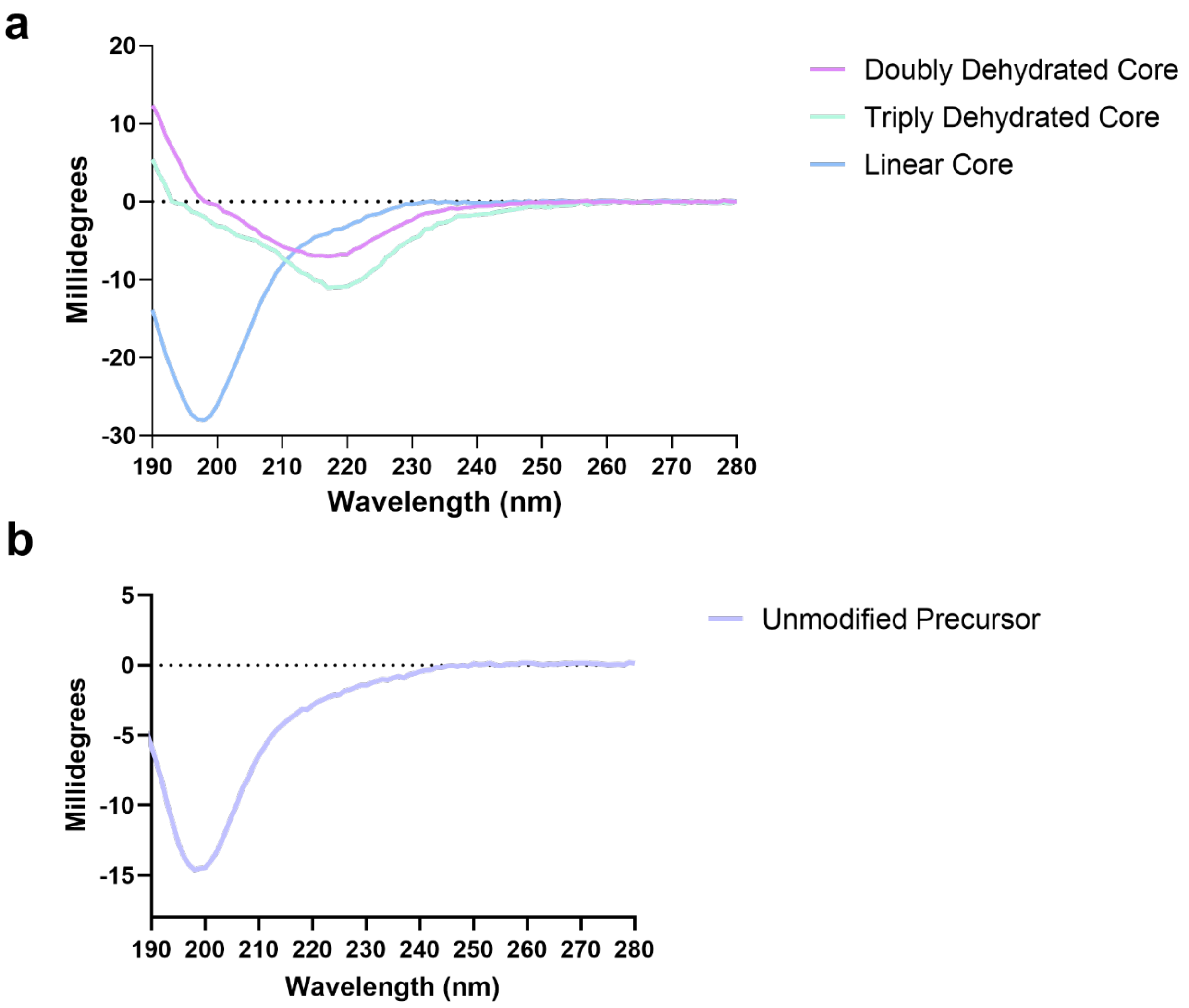


**Fig. S8** Circular dichroism spectra of putative ThfA core peptide and its dehydrated variants and full-length unmodified ThfA. a) Comparison of unmodified linear core peptide to doubly- and triply dehydrated core peptide. The introduction of esters in the doubly and triply dehydrated species endows the peptides with $\beta$ character whereas the unmodified core is a random coil. b) The full-length unmodified ThfA precursor is also a random coil.

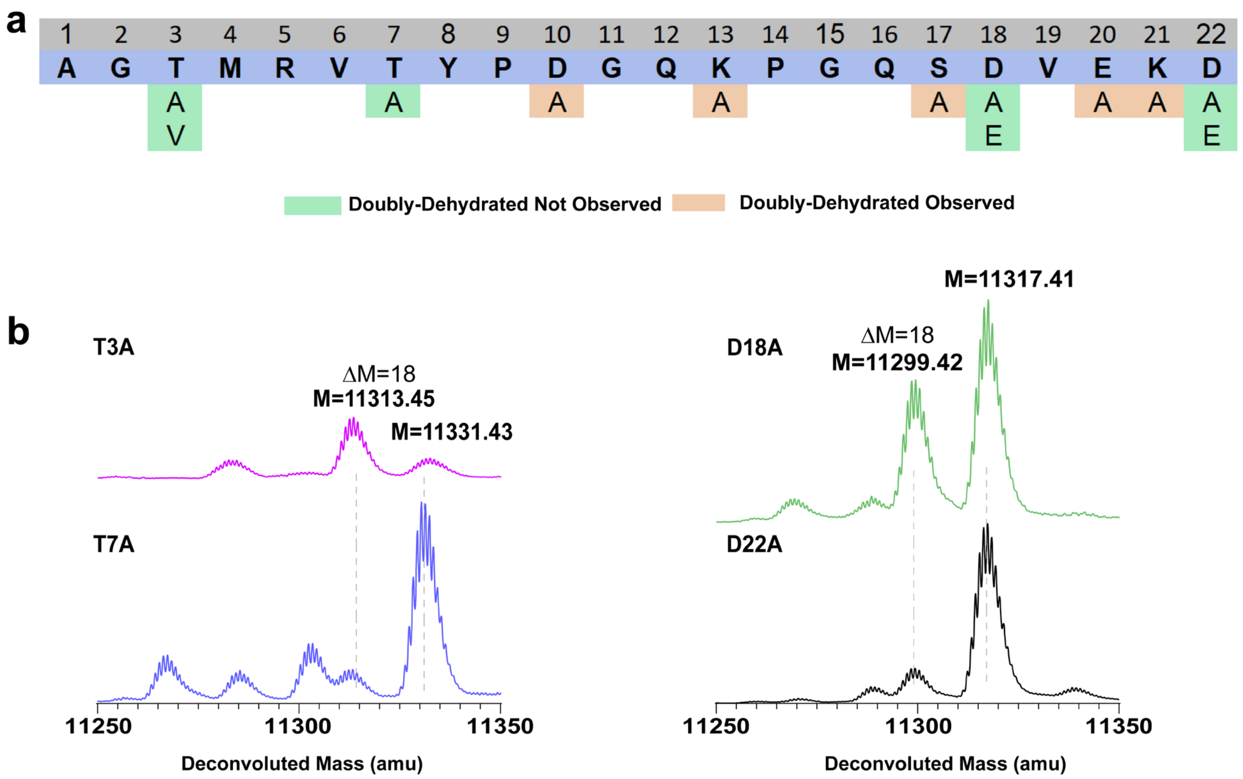


**Fig. S9** Mutagenesis studies on recombinant doubly dehydrated ThfA. a) An alanine scan was performed on all possible nucleophilic and electrophilic sidechains within the C-terminal 22 aa corresponding to the core peptide. The T3A, T7A, D18A, and D22A substitutions resulted in a complete loss of the doubly dehydrated product, whereas all other variants were still able to form the doubly dehydrated species. b) Deconvoluted mass spectra for the T3A, T7A, D18A, and D22A variants of ThfA.

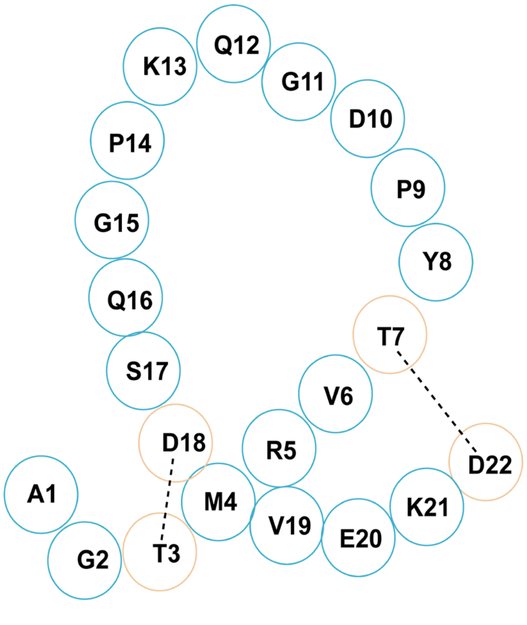


**Fig. S10** Two possible linkages for the ThFA core peptide. In the left "pretzel" conformer (left), T3 connects to D18 and T7 connects to D22. In the stem-loop conformer (right), T3-D22 and T7-D18 linkages are present instead.

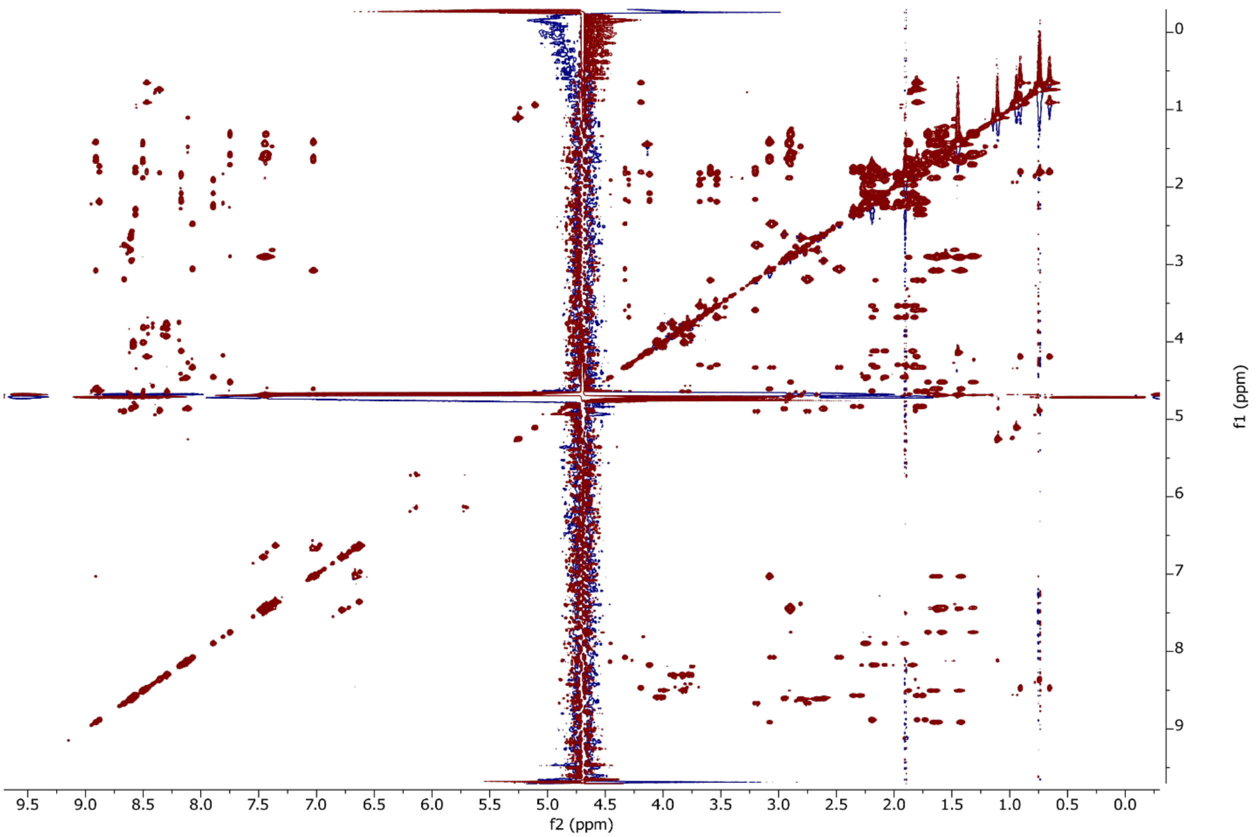


**Fig. S11** TOCSY spectrum of doubly dehydrated ThfA core peptide. Peak assignments are in Table S2.

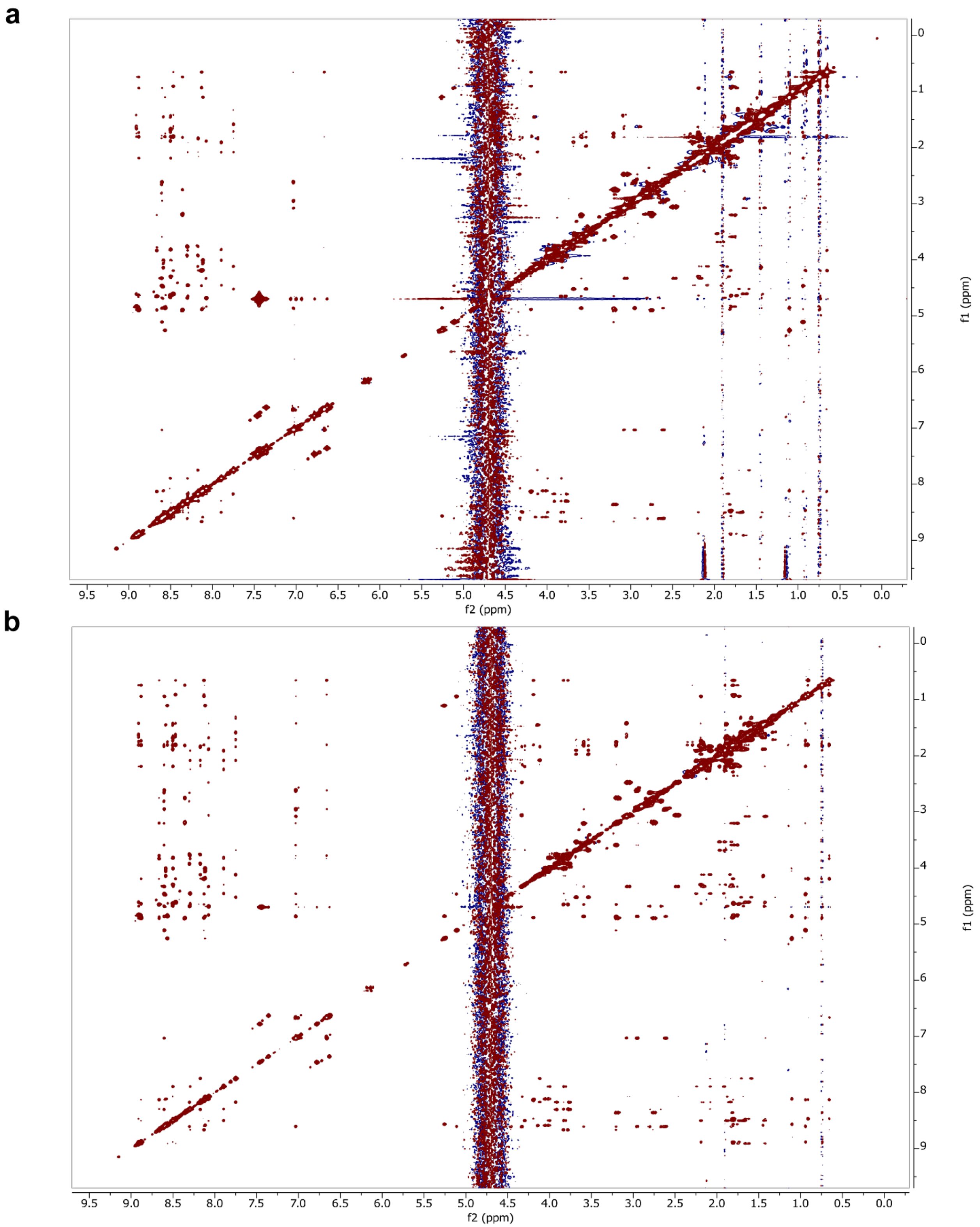


**Fig. S12** NOESY spectra of doubly dehydrated ThfA core peptide. a) mixing time of 150 ms b) mixing time of 700 ms. Peak assignments are in Table S2.

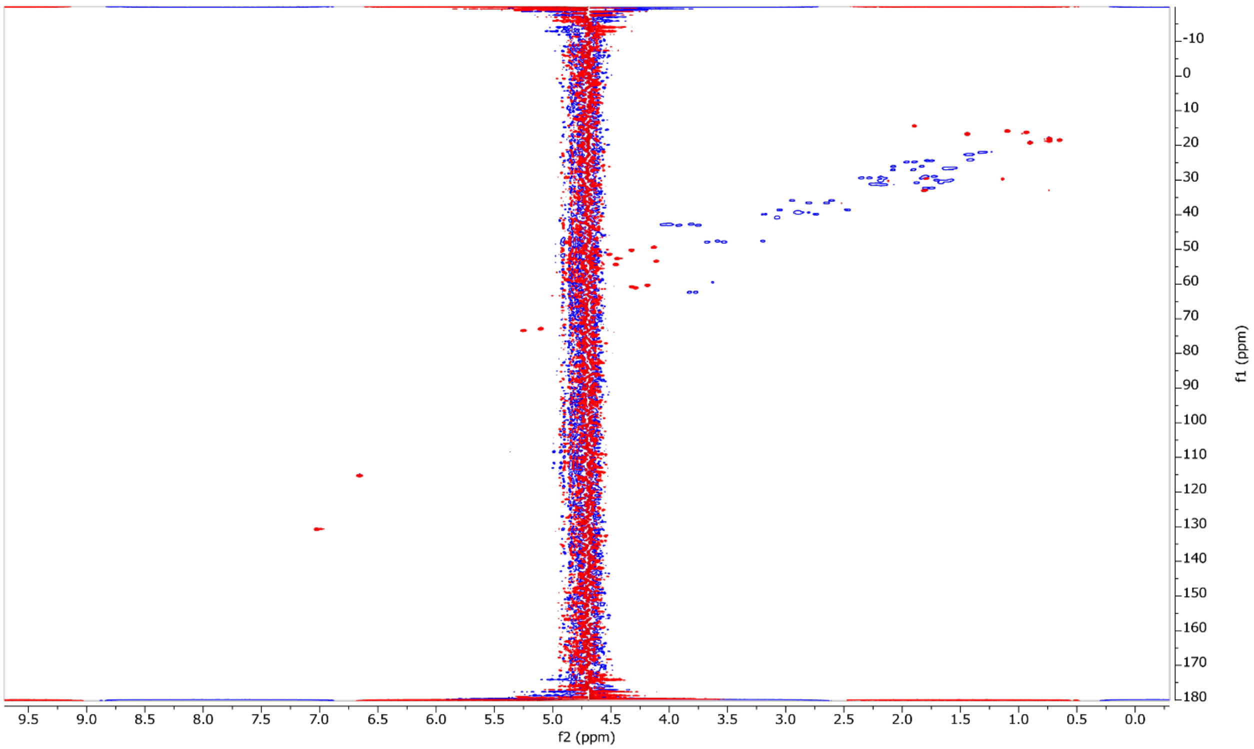


**Fig. S13** $^{1}H$-$^{13}C$ HSQC of doubly dehydrated ThfA core. Peak assignments are in Table S2.

**a**

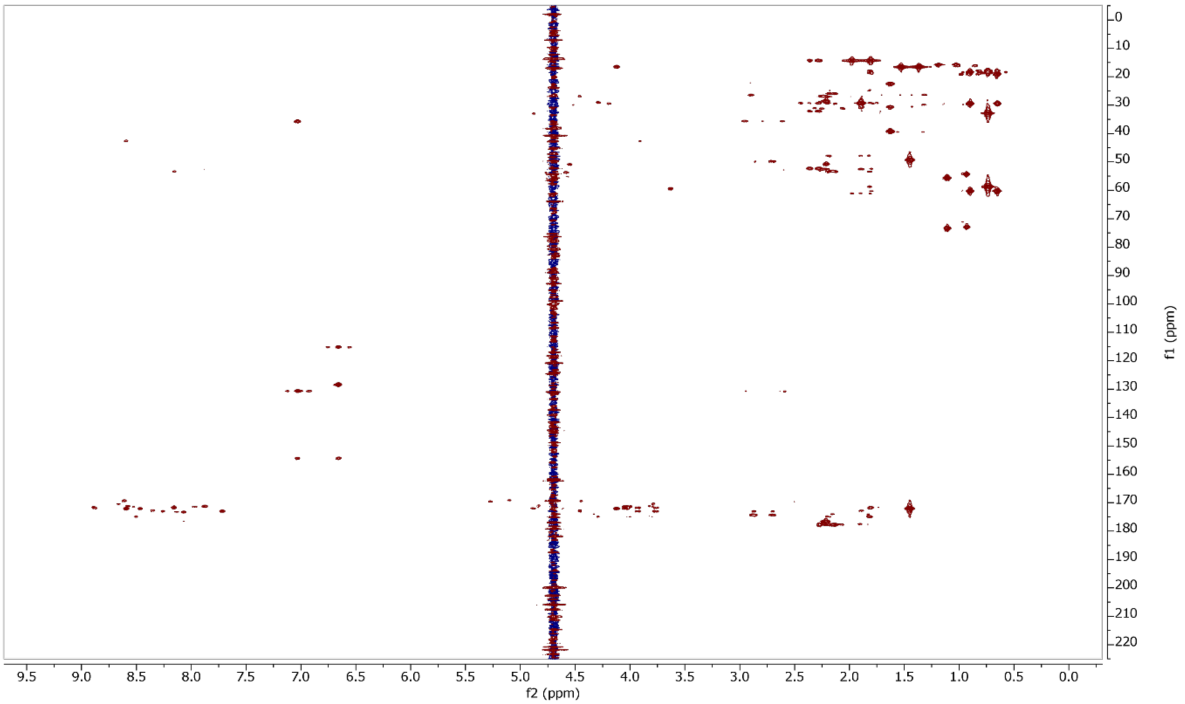


**b**

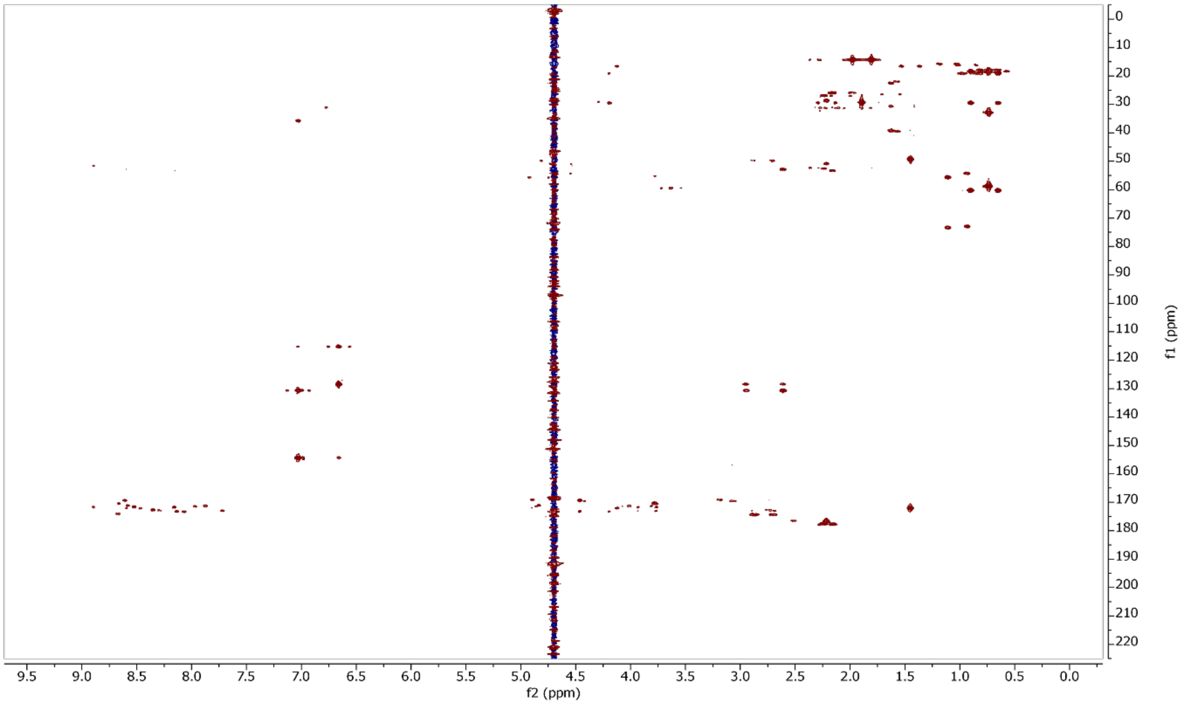


**Fig. S14** $^{1}H$-$^{13}C$ HMBC spectra of doubly dehydrated ThfA core. a) Spectrum optimized for 5 Hz couplings, b) spectrum optimized for 10 Hz couplings.

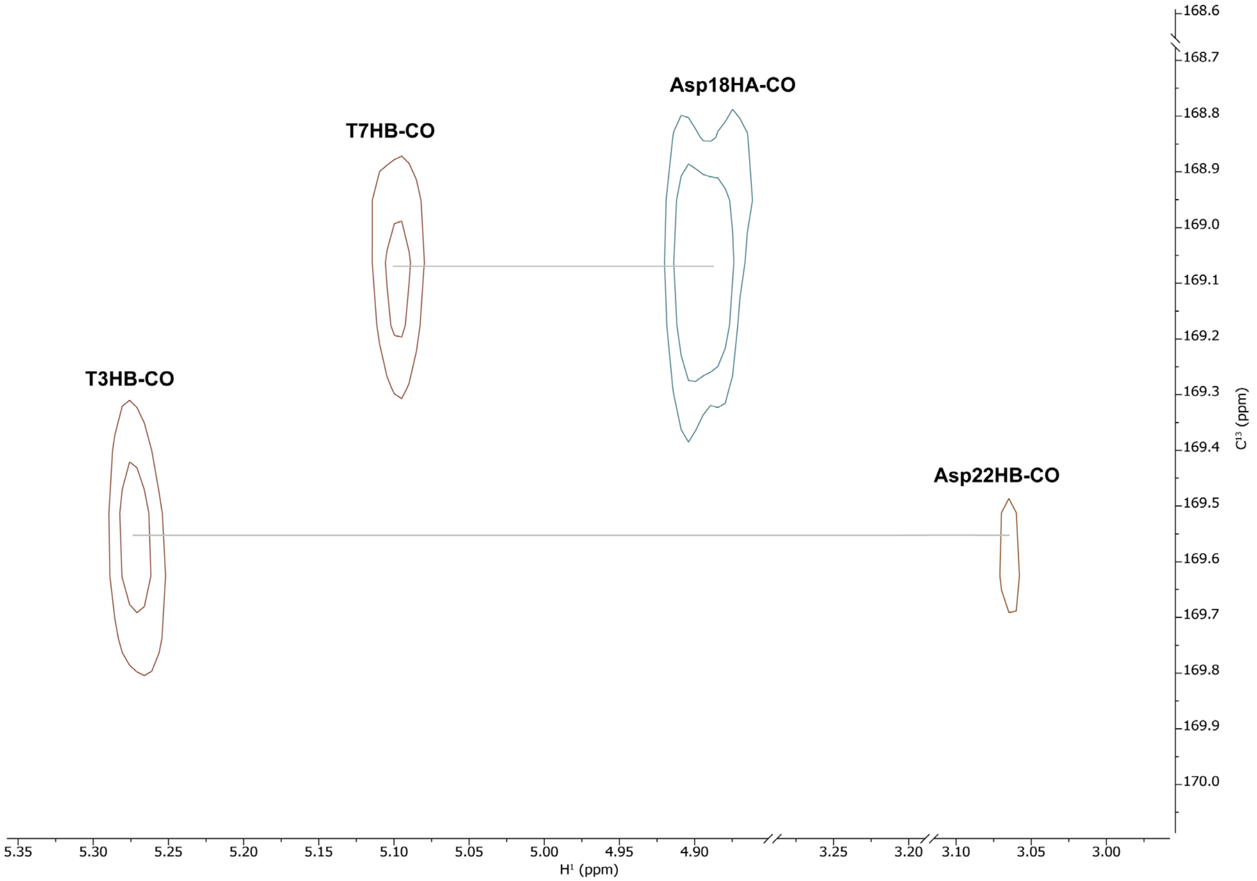


**Fig. S15** Zoomed view of the ThfA doubly dehydrated core $^{1}H$-$^{13}C$ HMBC spectrum (Fig. S14) with 5 Hz (red) and 10 Hz (blue) spectra overlaid. Notably, the signals for the carbonyl carbons are shifted upfield relative to carbonyl carbons in the aspartic acid sidechain (average chemical shift of 177 ppm). A crosspeak is observed between the D18 carbonyl and the β proton of T7. A similar crosspeak is observed between the D22 carbonyl and the β proton of T3.

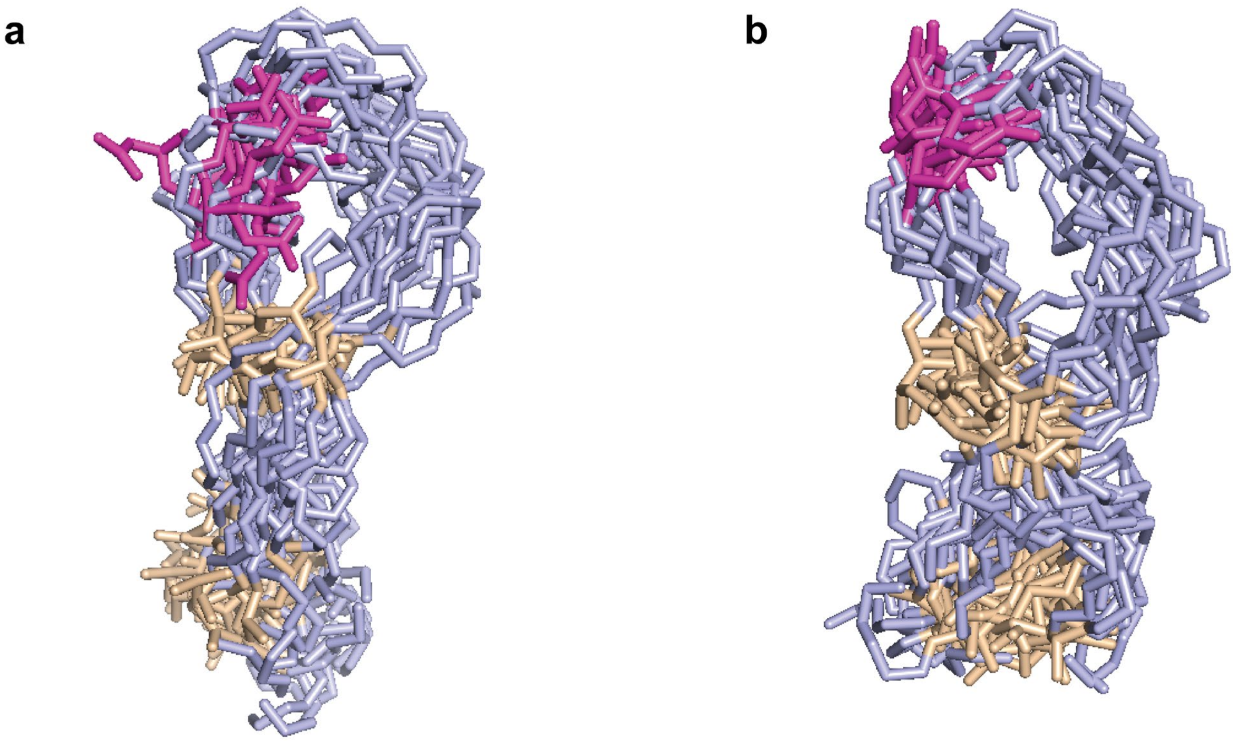


**Fig. S16** Structural models of pre-fuscimiditide and fuscimiditide. a) Overlay of the top 20 structures of pre-fuscimiditide; the D10 residue is in magenta. b) Overlay of the top 20 structures of fuscimiditide; the aspartimide formed from D10 is colored magenta. Structures were generated in CYANA 2.1 and energy minimized using Avogadro.

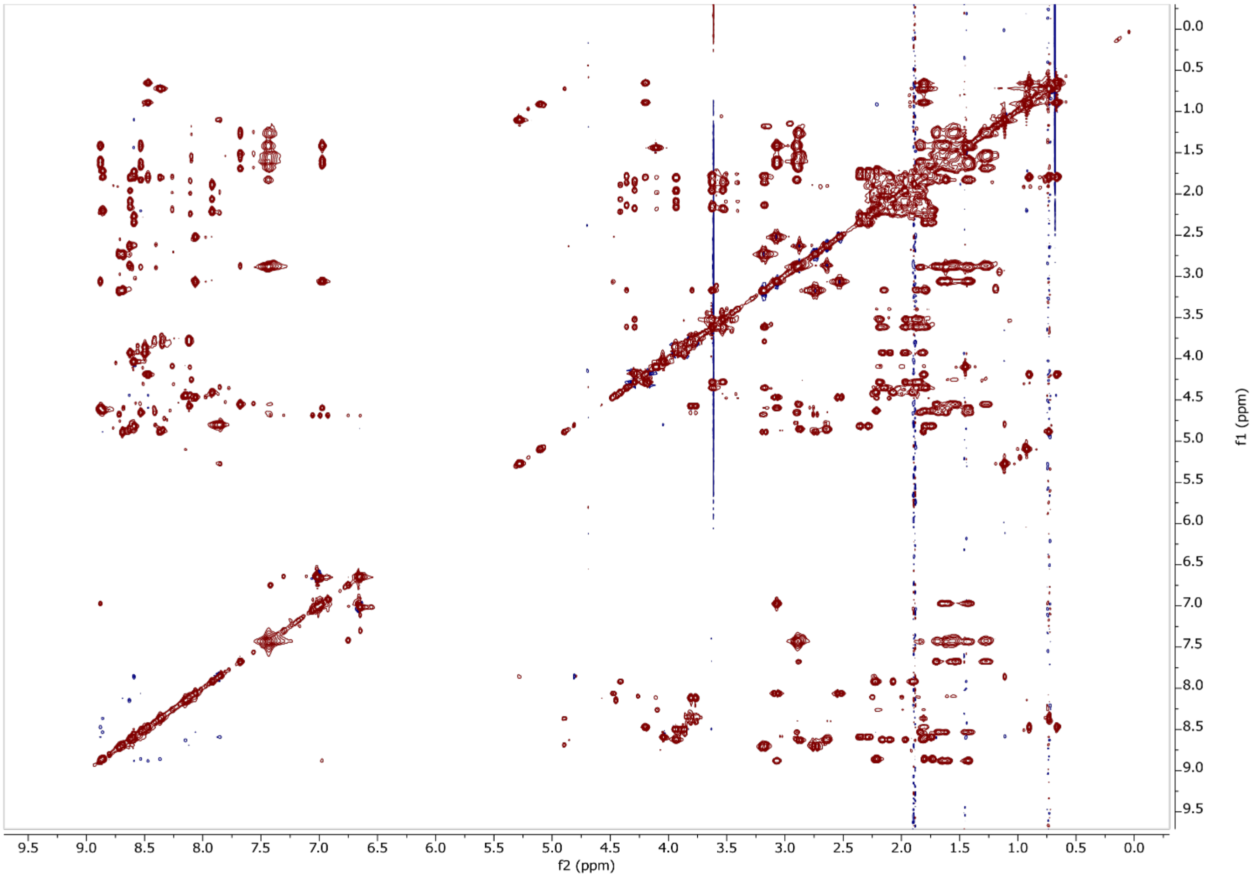


**Fig. S17** TOCSY spectrum of triply dehydrated ThfA core peptide. Peak assignments are in Table S2.

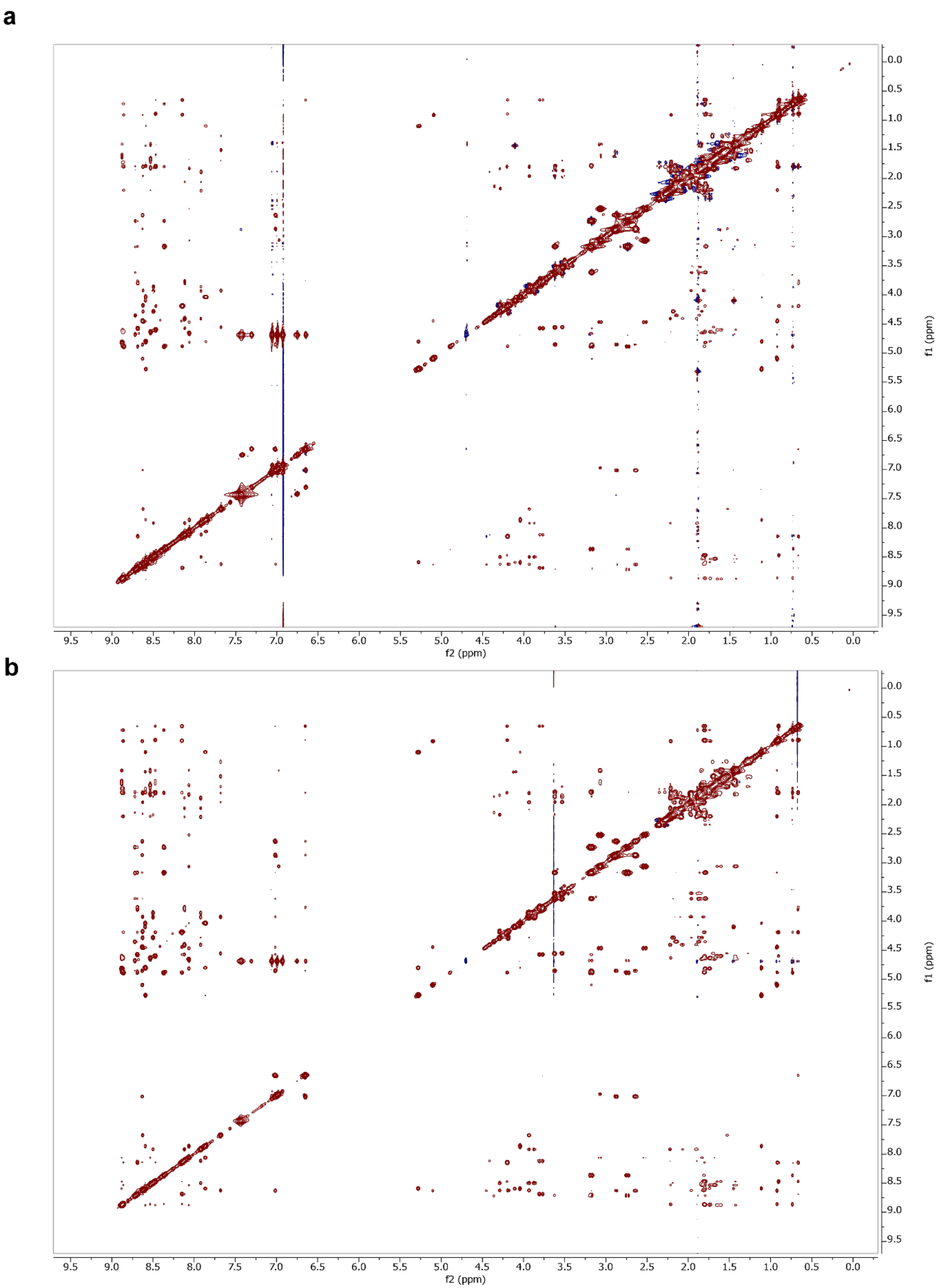


**Fig. S18** NOESY spectra of triply dehydrated ThfA core peptide. a) mixing time of 150 ms b) mixing time of 700 ms. Peak assignments are in Table S2.

a

b

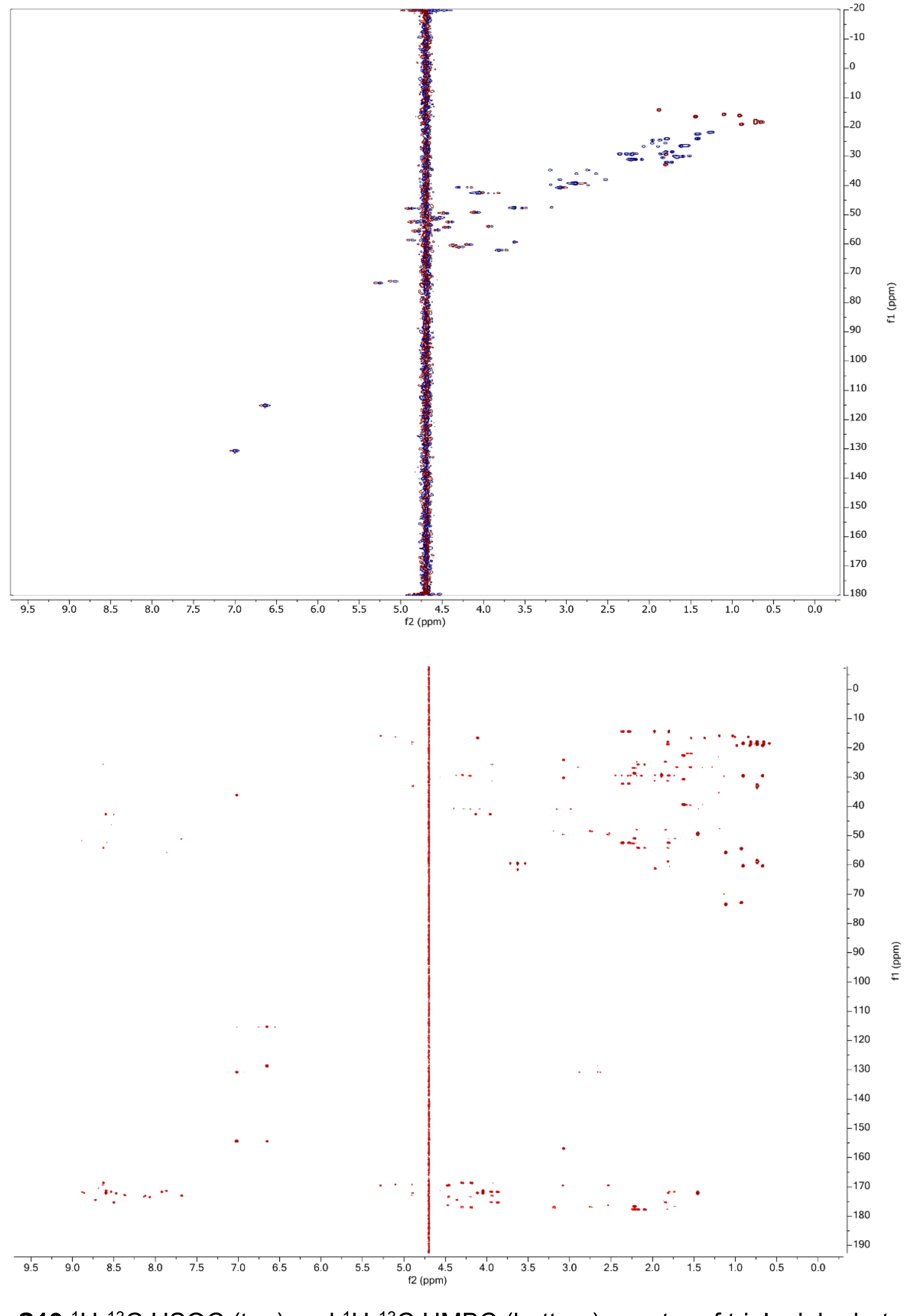


**Fig. S19** $^{1}H$-$^{13}C$ HSQC (top) and $^{1}H$-$^{13}C$ HMBC (bottom) spectra of triply dehydrated ThfA core peptide. The HMBC spectrum was optimized for 10 Hz.

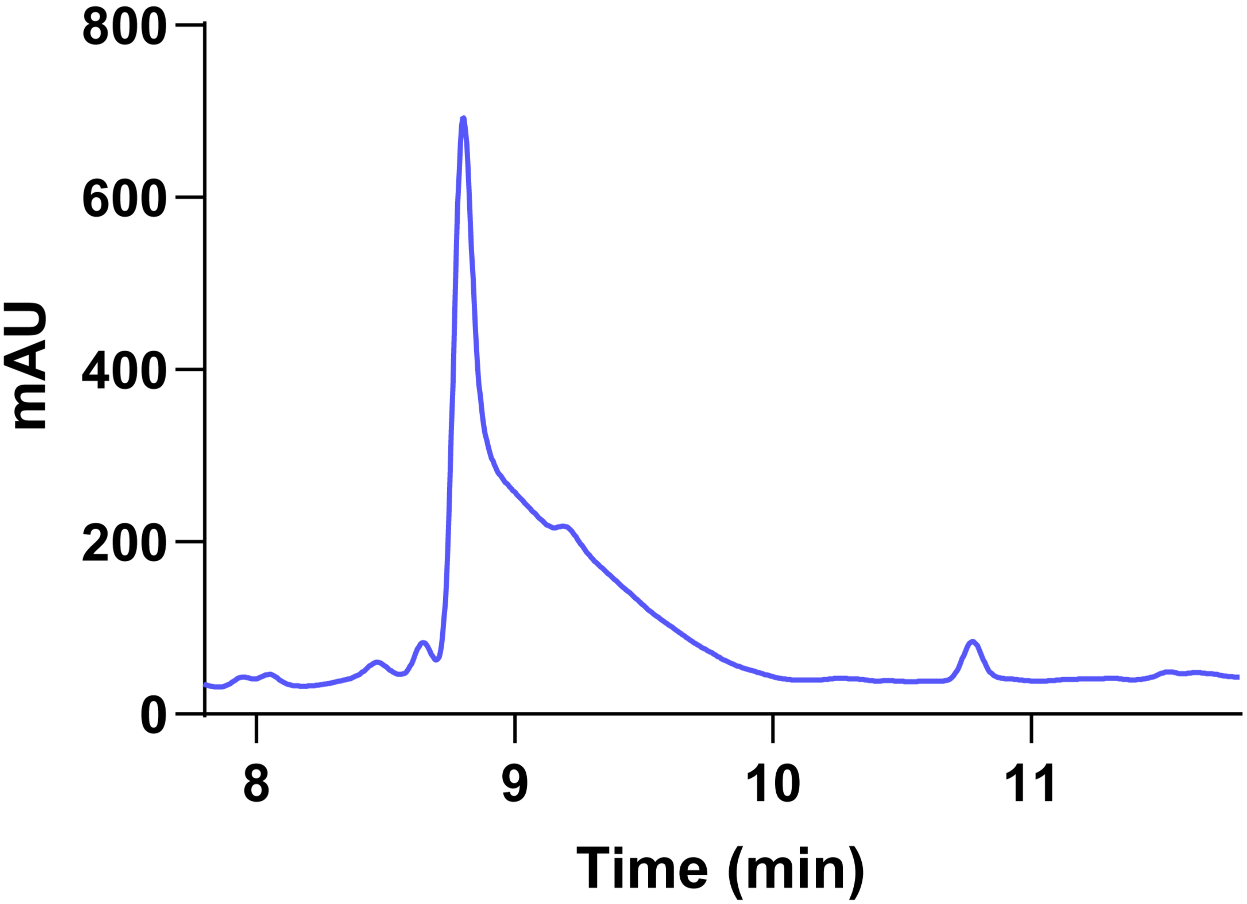


**Fig. S20** HPLC trace of trypsin digested doubly dehydrated ThfA. The peak at 8.8 min corresponds to the retention time of pre-fuscimiditide.

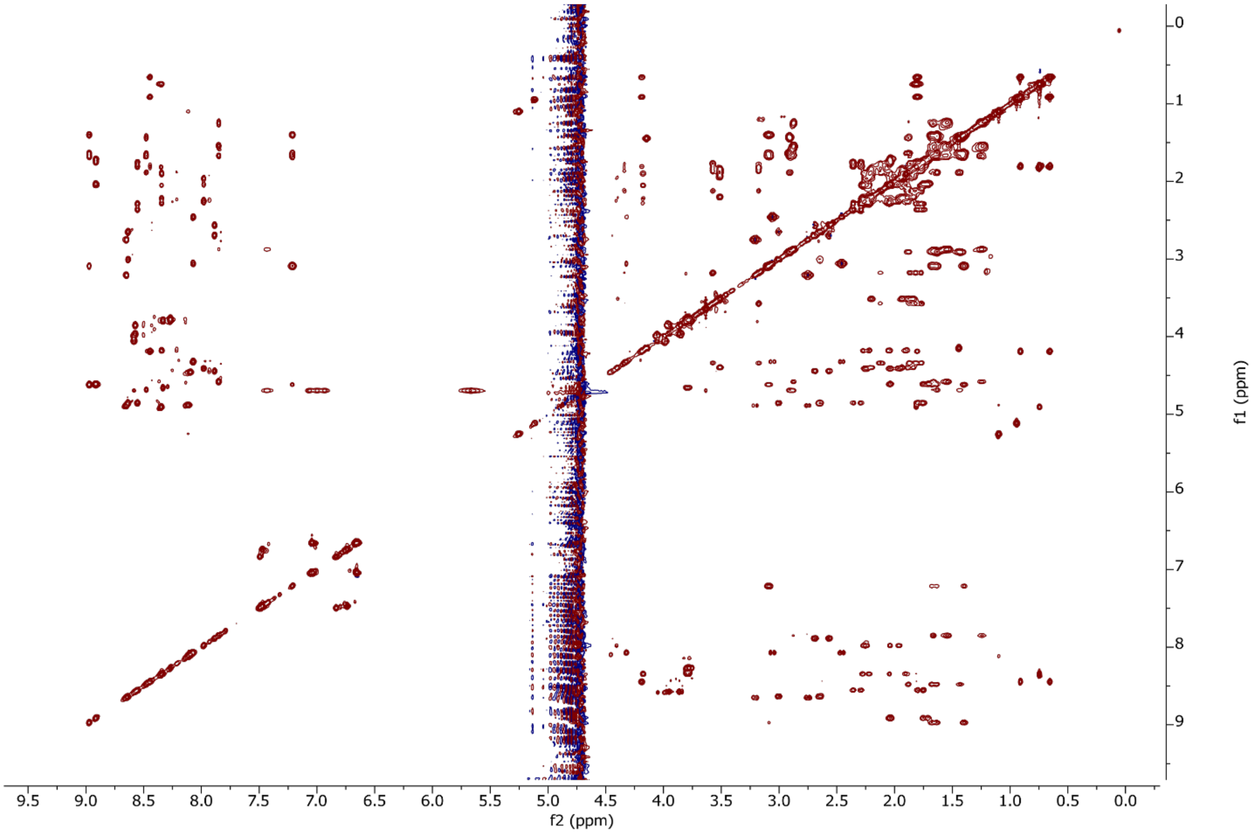


**Fig. S21** TOCSY spectrum of iso pre-fuscimiditide. Peak assignments are in Table S2.

**a**

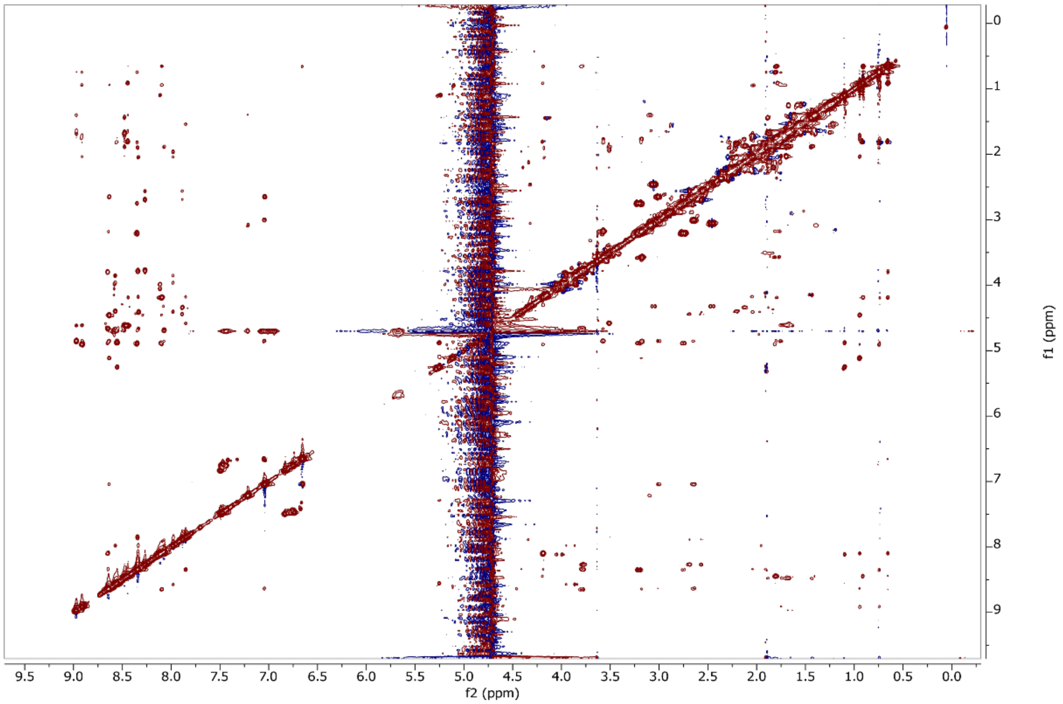


**b**

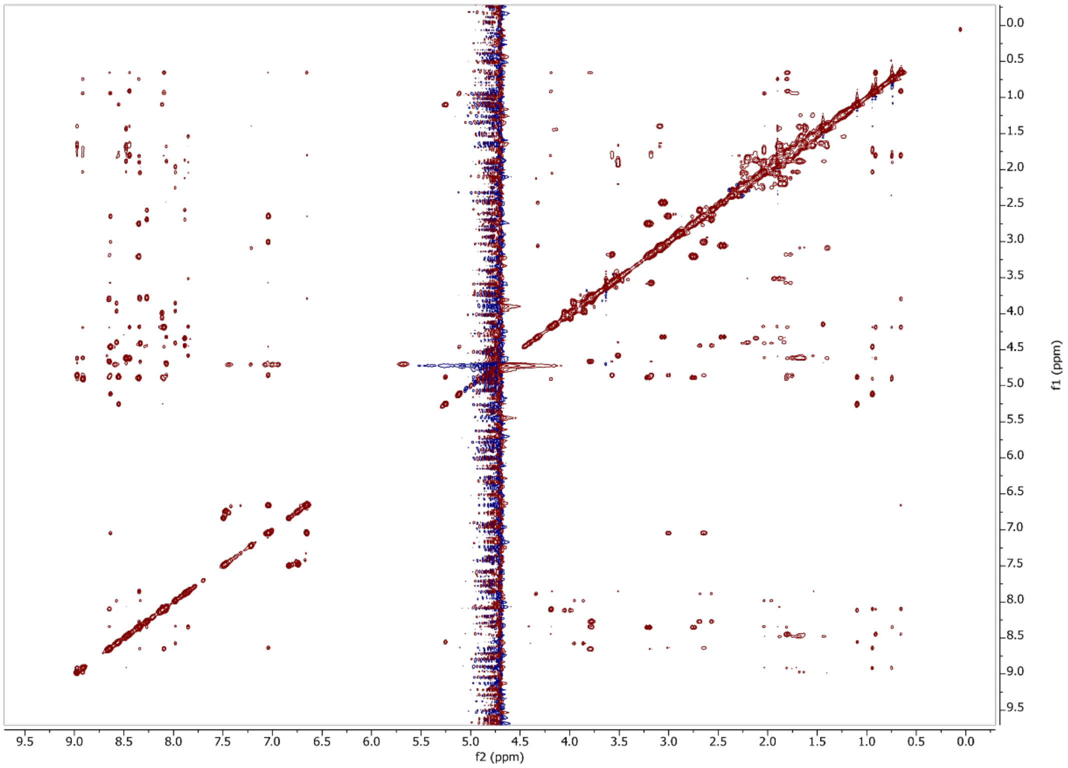


**Fig. S22** NOESY spectra of iso pre-fuscimiditide. a) 150 ms mixing time, b) 700 ms mixing time. Peak assignments are in Table S2.

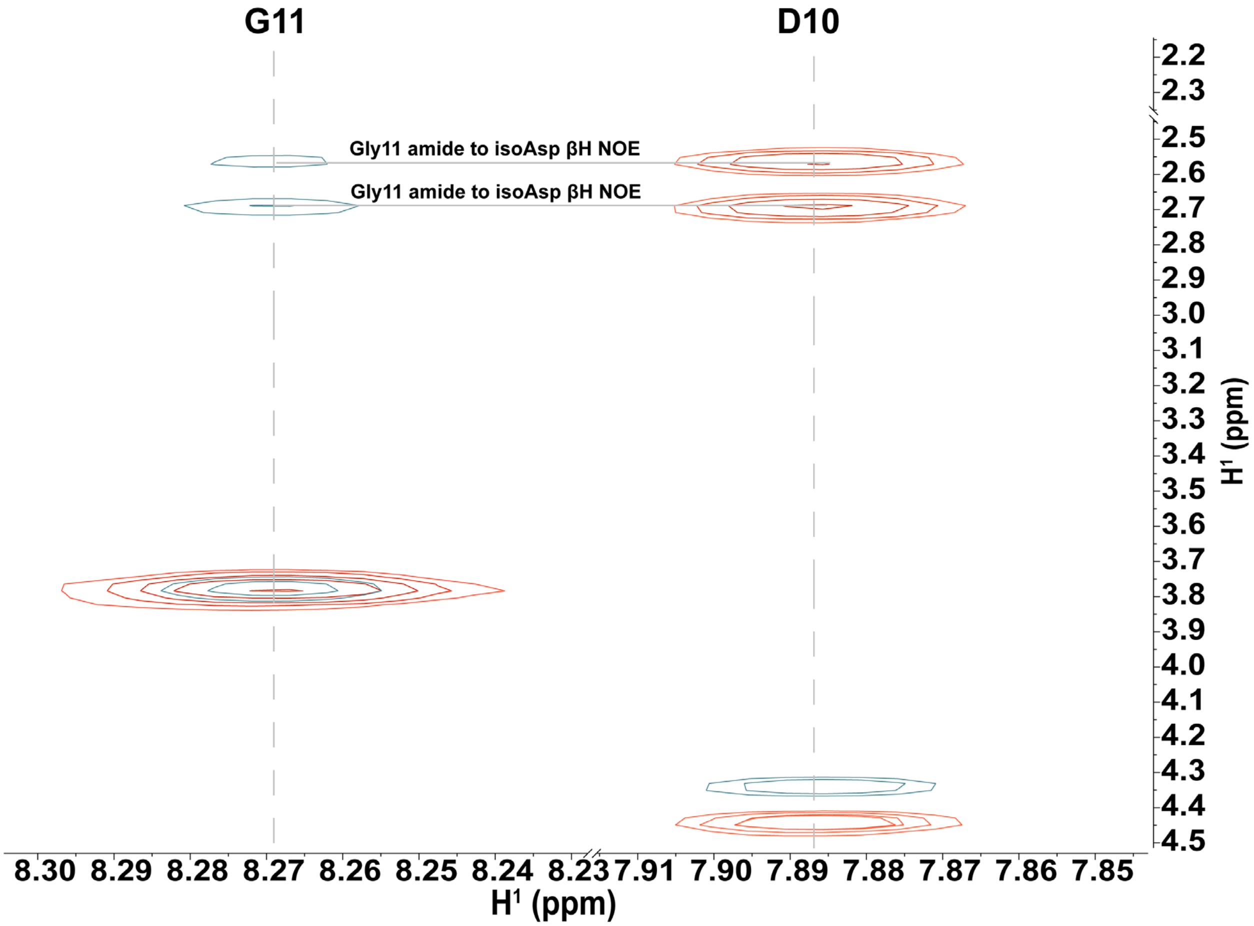


**Fig. S23** Overlay of TOCSY (orange) and NOESY (turquoise) spectra of pre-fuscimiditide. Crosspeaks showing the canonical backbone connectivity between D10 and G11 are highlighted. An analogous figure for iso pre-fuscimiditide is in Fig. 4d.

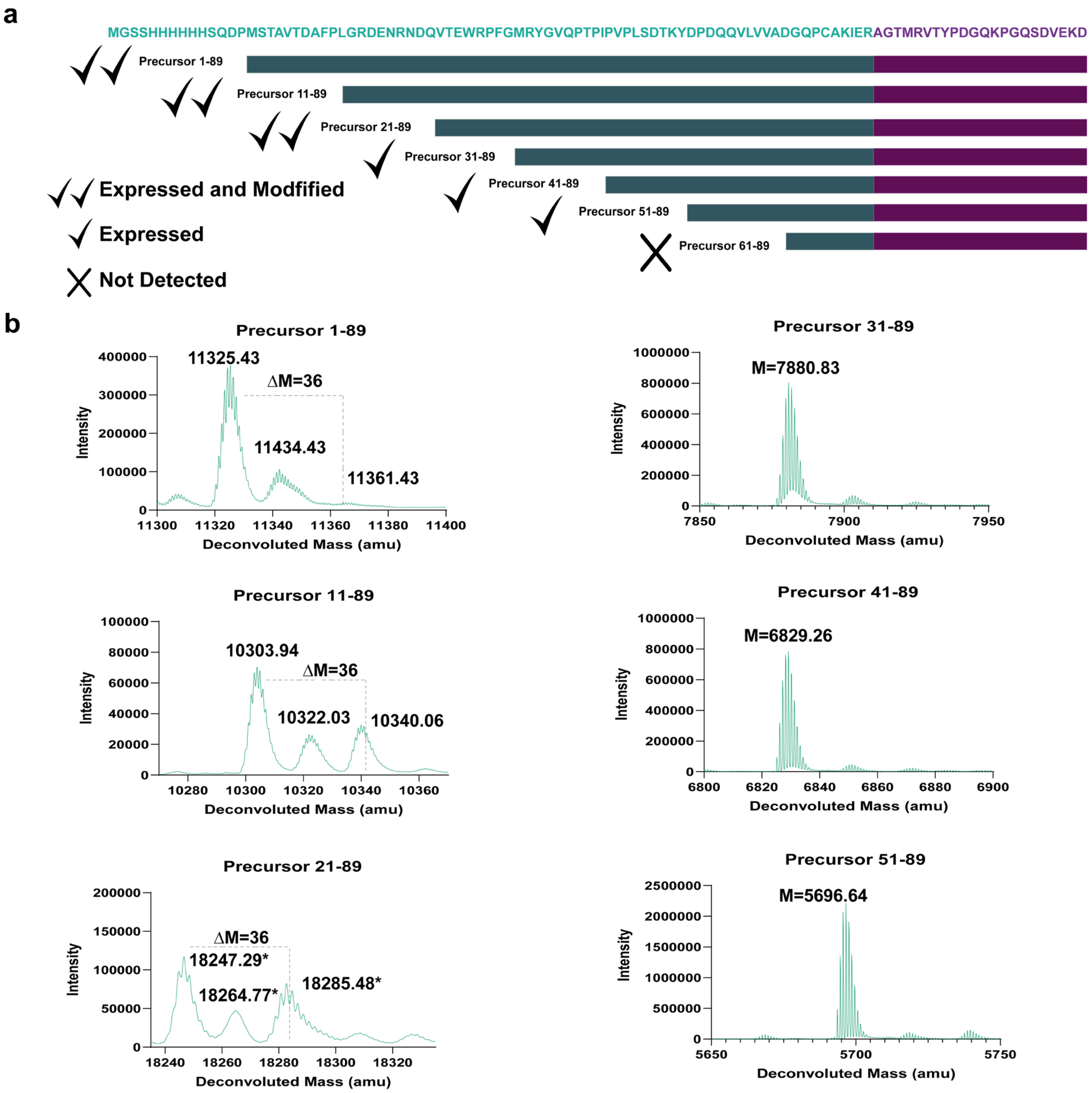


**Fig. S24** Leader peptide truncation studies on ThfA. a) The indicated truncation variants were cloned and coexpressed with ATP-grasp enzyme ThfB. While all variants except for the 60 aa truncate expressed, only the 10 and 20 aa truncates were still esterified. b) Deconvoluted mass spectra for each of the expressed variants showing dehydrations when present. For the Precursor 21-89 construct, only a disulfide bonded protein was observed, leading to a roughly doubled mass, indicated by asterisks.

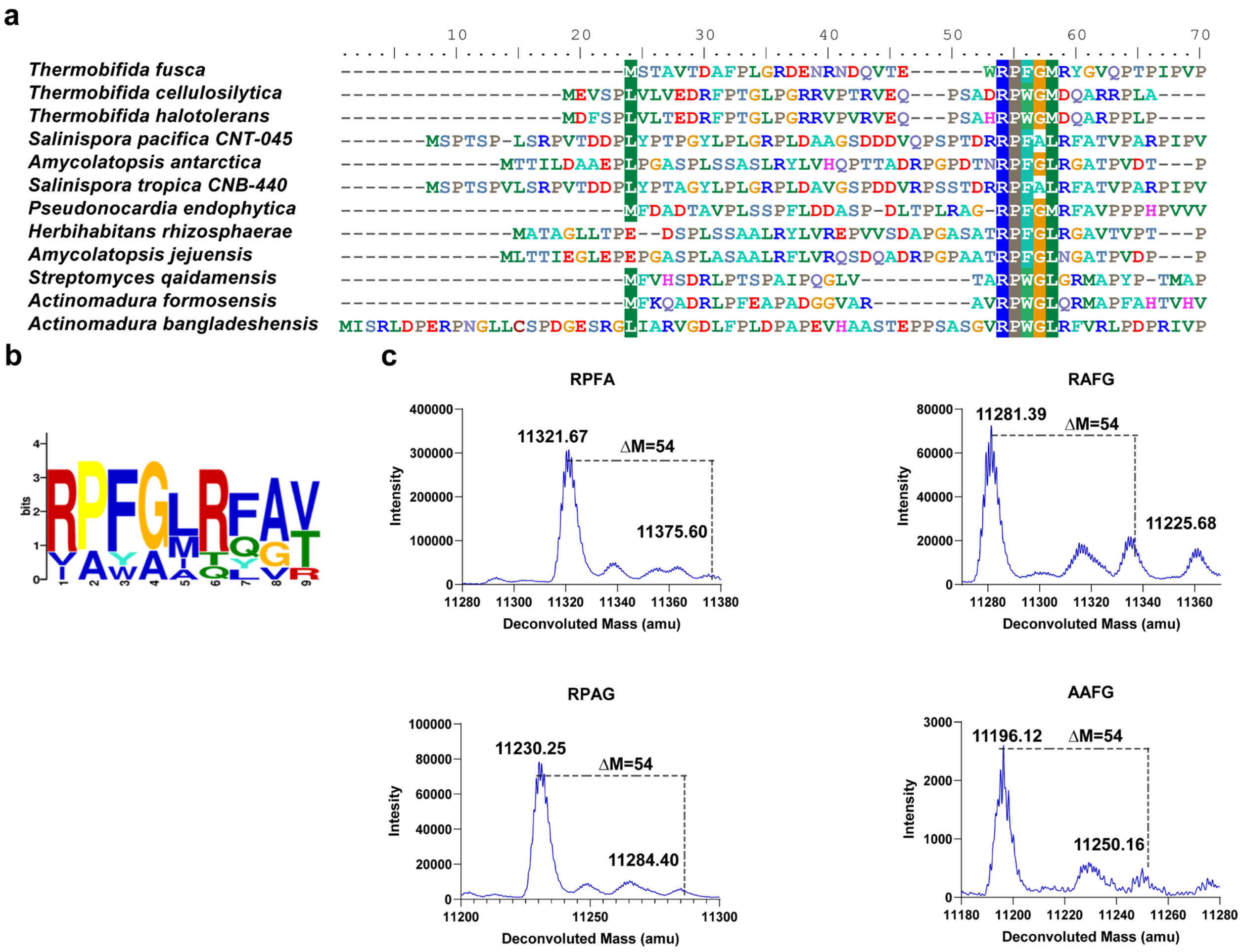


**Fig. S25** Alignment and mutagenesis of the ThfA leader peptide. a) Alignment of leader portions of several putative aspartimide modified peptides with fuscimiditide (labeled *Thermobifida fusca* in the alignment). b) Sequence logo of conserved motif from these leader peptides generated by MEME. The RPFG tetrapeptide at the beginning of this motif is strongly conserved. c) Mutagenesis of the putative RPFG motif does not lead to abrogation of either ATP-grasp or *O*-methyltransferase activity.

**a**

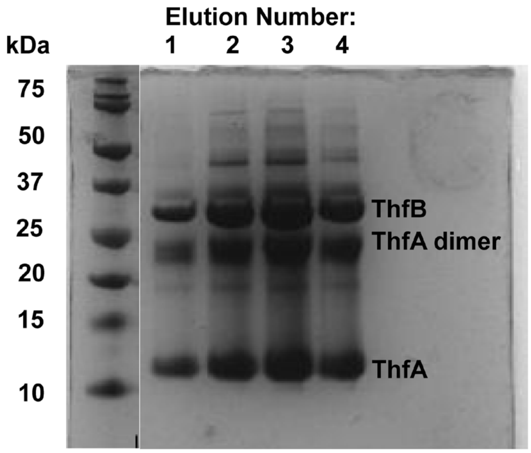


**b**

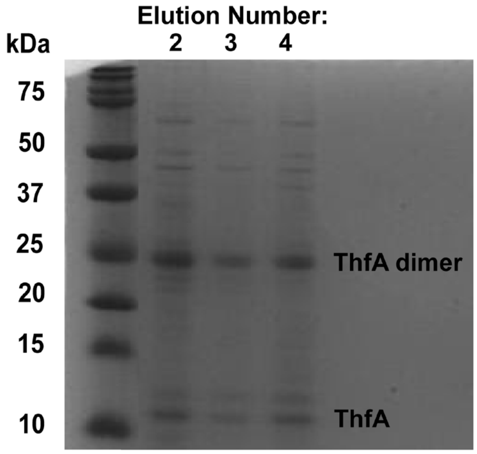


**Fig. S26** Native and denaturing purification of His-tagged ThfA coexpressed with untagged ThfB. a) When purified under native conditions (ie elution from Ni-NTA with imidazole), His-tagged ThfA coelutes with a stoichiometric amount of untagged ThfB. b) In order to isolate doubly dehydrated ThfA, the material from the native purification in part a) was repurified under denaturing conditions in 8 M urea to provide purified ThfA. In both purification conditions, disulfide bonding is observed, so DTT was added to all *in vitro* reactions.

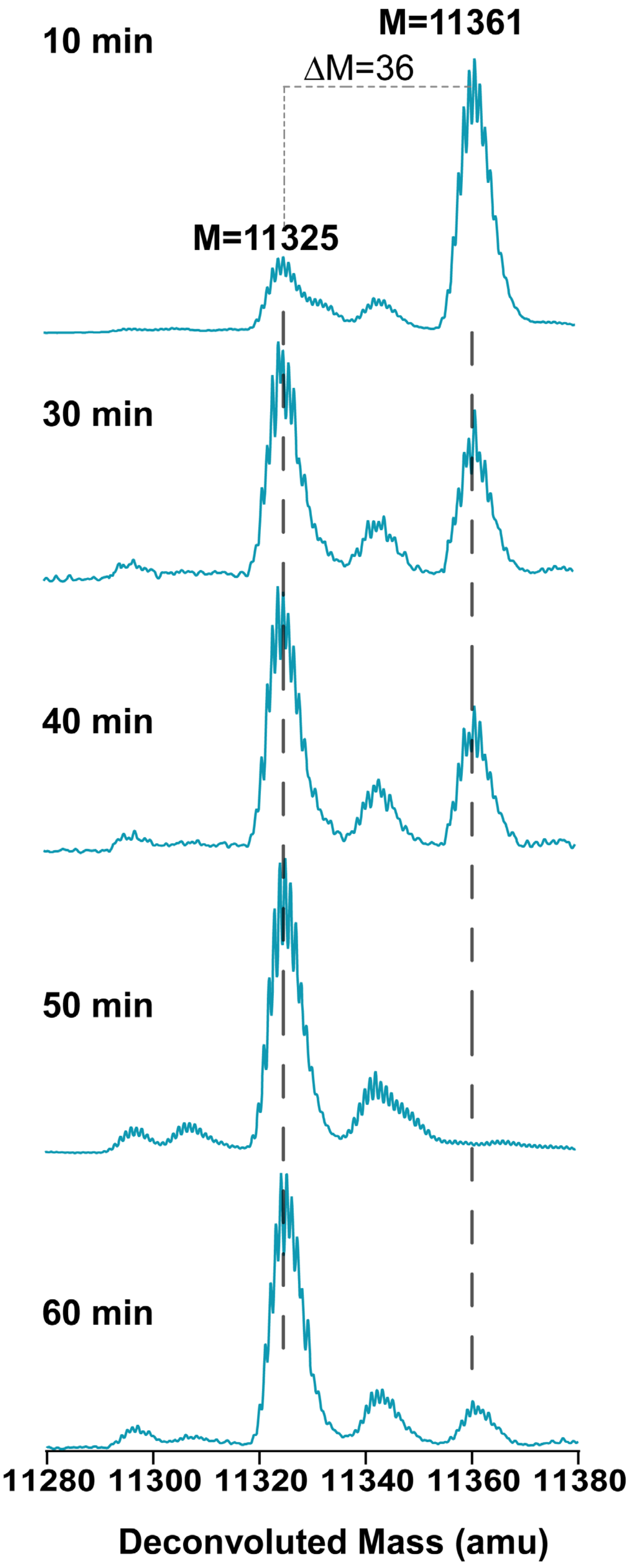


**Fig. S27** Time course of reaction of ThfA and ThfB with 100 mM KCl as an additive. As compared to the reaction lacking KCl (Fig. 5a) ester formation is more rapid.

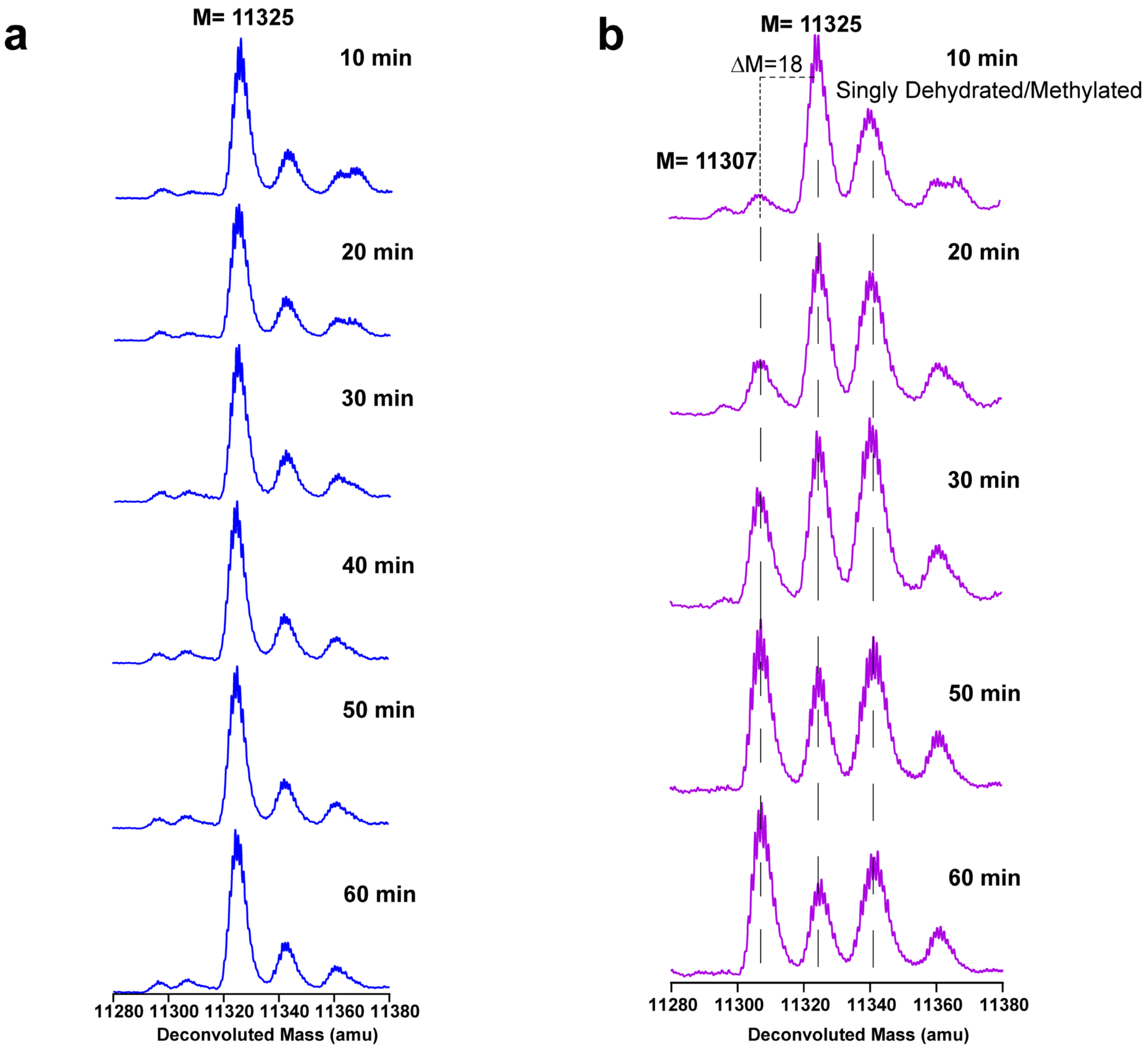


**Fig. S28** Time course of the reaction between denatured ThfA and ThfM in the presence and absence of SAM. a) In the absence of SAM, a third dehydration is not observed. b) Formation of the third dehydration is dependent on the presence SAM and proceeds to ~50% completion within 60 min. The isotopic distributions for singly-dehydrated ThfA and methylated doubly-dehydrated ThfA overlap.

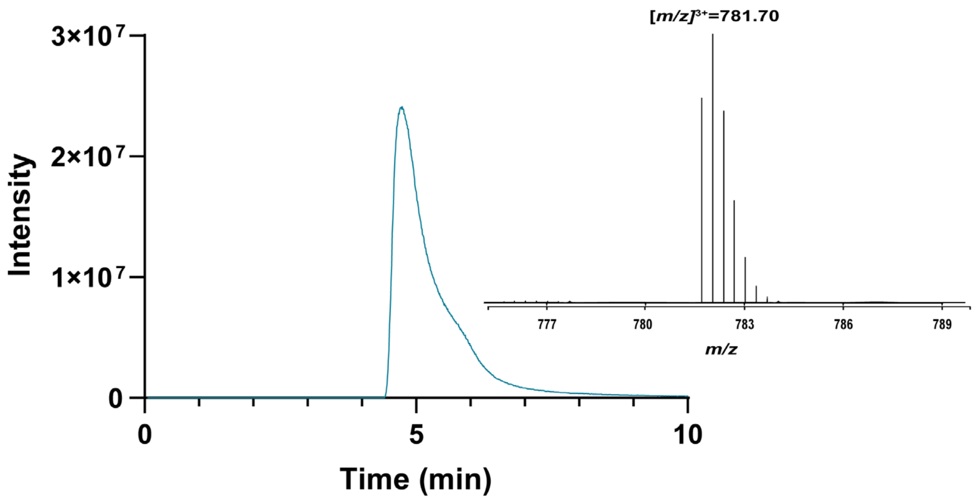


**Fig. S29** TIC chromatogram of the reaction between pre-fuscimiditide and ThfM. The inset is the extracted mass corresponding to unreacted pre-fuscimiditide. No masses corresponding to fuscimiditide were observed, demonstrating the requirement of at least part of the leader peptide for the function of ThfM.

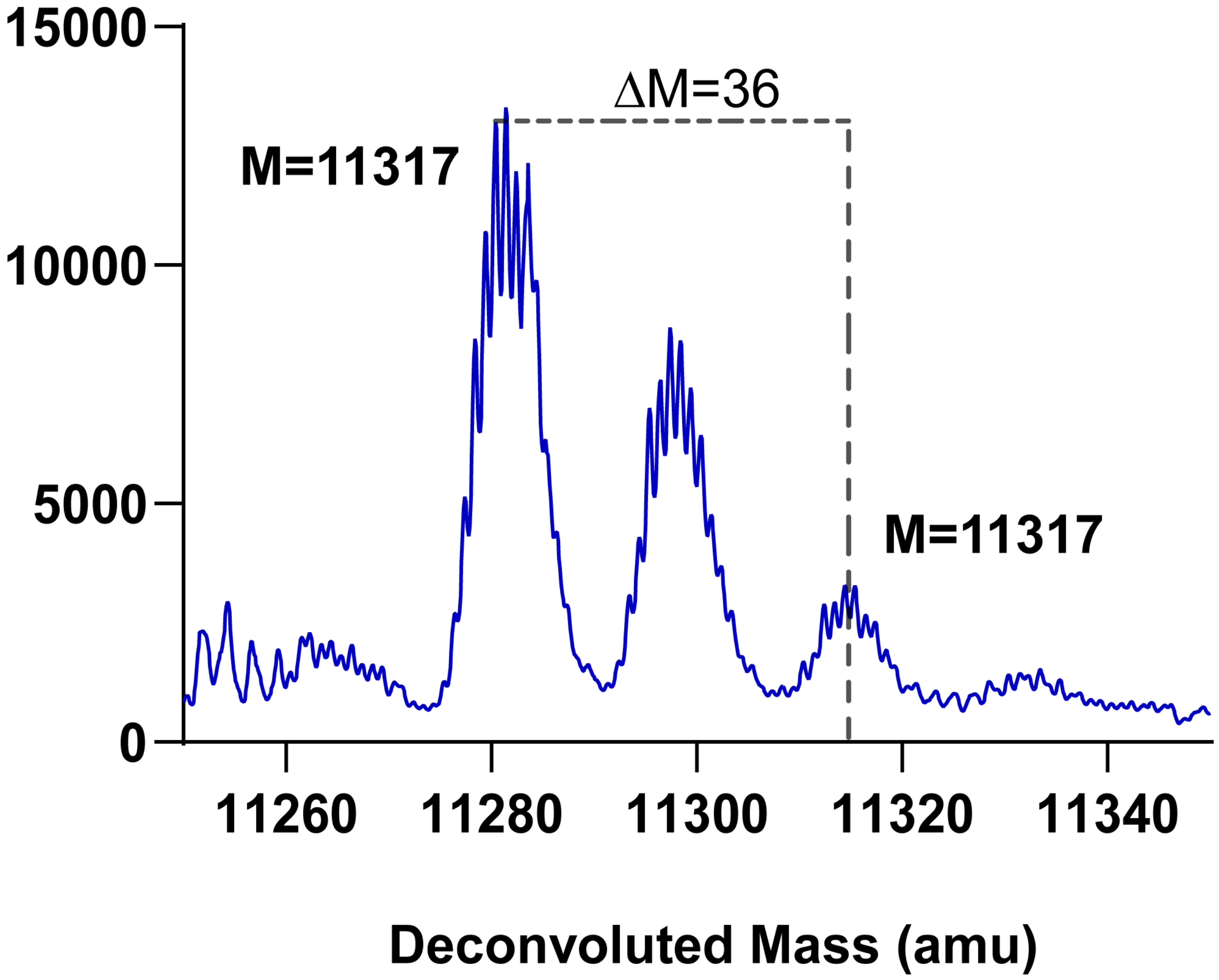


**Fig. S30** Deconvoluted mass spectrum of the products of an *in vitro* reaction comprised of ThfA D77A, ThfB, ThfM, ATP, and SAM. D77 in ThfA corresponds to D10 in fuscimiditide, the aspartate residue that is converted to aspartimide. Only two dehydrations are observed, presumably corresponding to ester linkages. This data further suggests that D10 is the site of aspartimidylation in fuscimiditide.

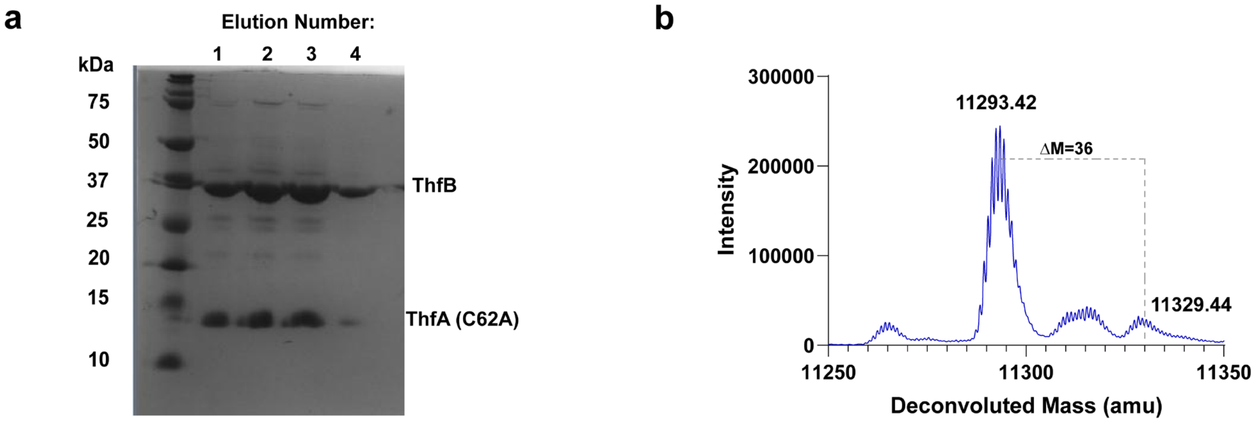


**Fig. S31** The ThfA C62A variant still binds to ThfB and can be doubly dehydrated. a) SDS-Page of His-tagged ThfA C62A coexpressed with untagged ThfB. ThfA still pulls down a stoichiometric amount of ThfB, though as expected, no ThfA dimers are observed (refer to Fig. S26). b) Deconvoluted mass spectrum of ThfA C62A coexpressed with ThfB, showing that this substrate variant is still doubly dehydrated.

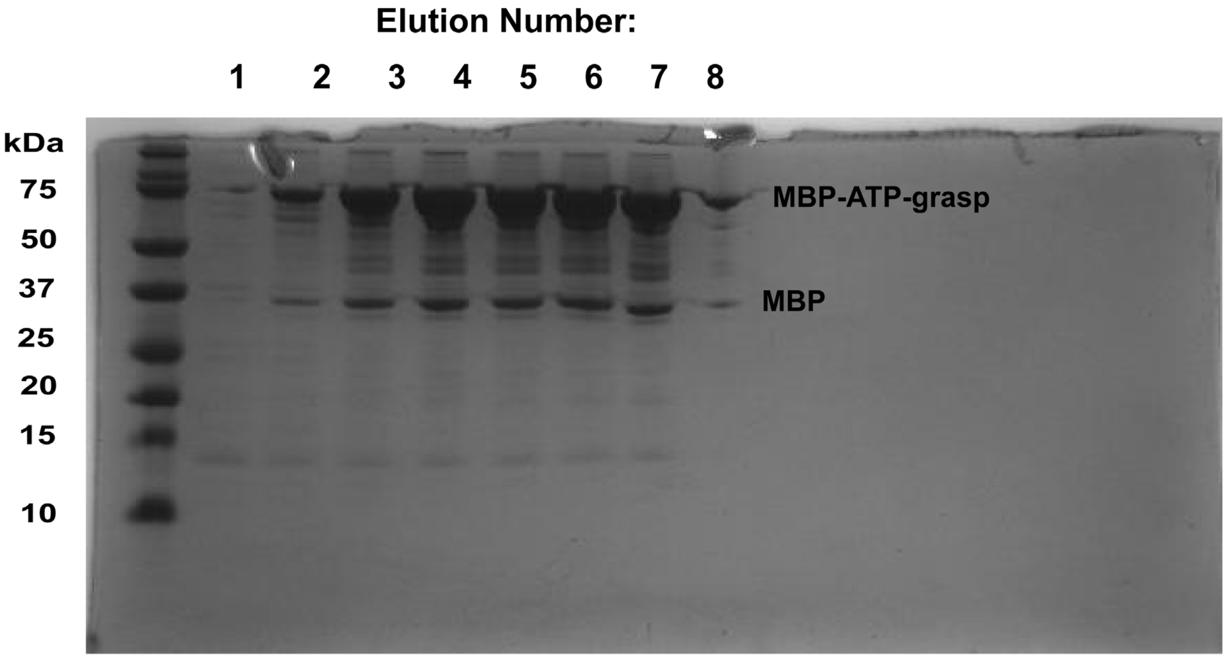


**Fig. S32** SDS-Page analysis of MBP-ThfB fusion protein for biolayer interferometry studies. The lanes correspond to elutions from an amylose column. Native MBP is a contaminant in these elutions.

**a**

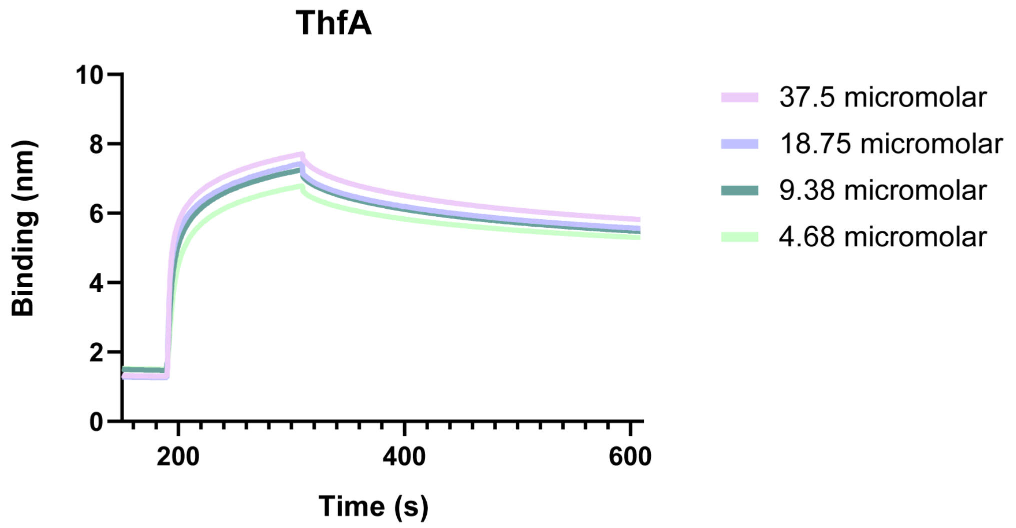


| $K_d$ (M) | $k_a$ ($M^{-1}s^{-1}$) | $k_d$($s^{-1}$) |
|---|---|---|
| $2.78 \times 10^{-8}$ | $5.89 \times 10^{4} \pm 0.09 \times 10^{4}$ | $1.63 \times 10^{-3} \pm 7.01 \times 10^{-5}$ |

**b**

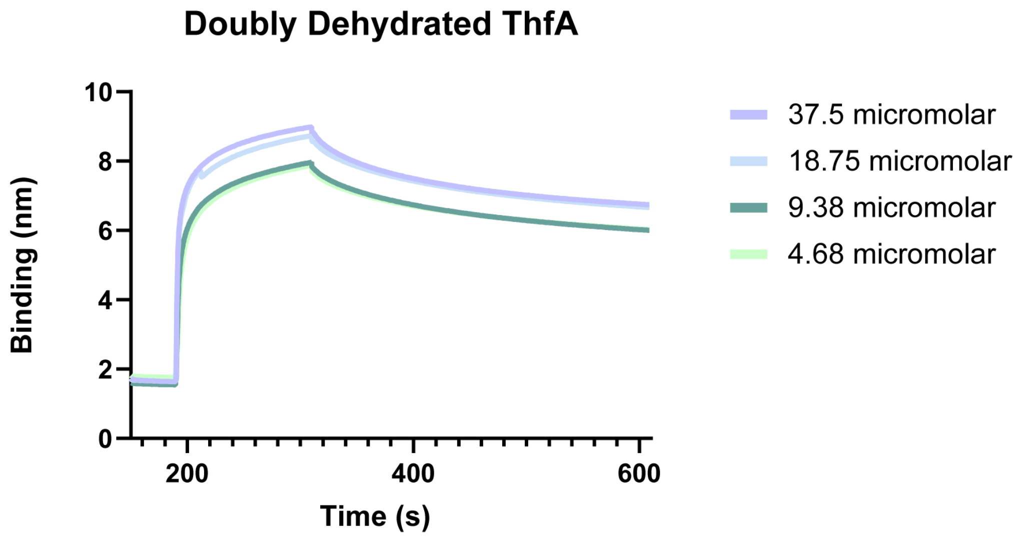


| $K_d$ (M) | $k_a$ ($M^{-1}s^{-1}$) | $k_d$($s^{-1}$) |
|---|---|---|
| $1.78 \times 10^{-8}$ | $9.28 \times 10^{4} \pm 0.17 \times 10^{4}$ | $1.65 \times 10^{-3} \pm 6.29 \times 10^{-5}$ |

**Fig. S33** Biolayer interferometry data for the ThfA/ThfB interaction. a) Kinetic data for the interaction between unmodified ThfA and ThfB. In these experiments, a constant amount of His-tagged ThfA C62A was loaded onto the biosensor, and different concentrations of the MBP-ThfB analyte were tested from 4.68 μM to 37.5 μM. b) As in part a) except that doubly dehydrated ThfA was loaded on the biosensor. Below the graphs are the global fits of the data, showing that both the substrate ThfA and product doubly dehydrated ThfA bind ThfB with nanomolar affinity.

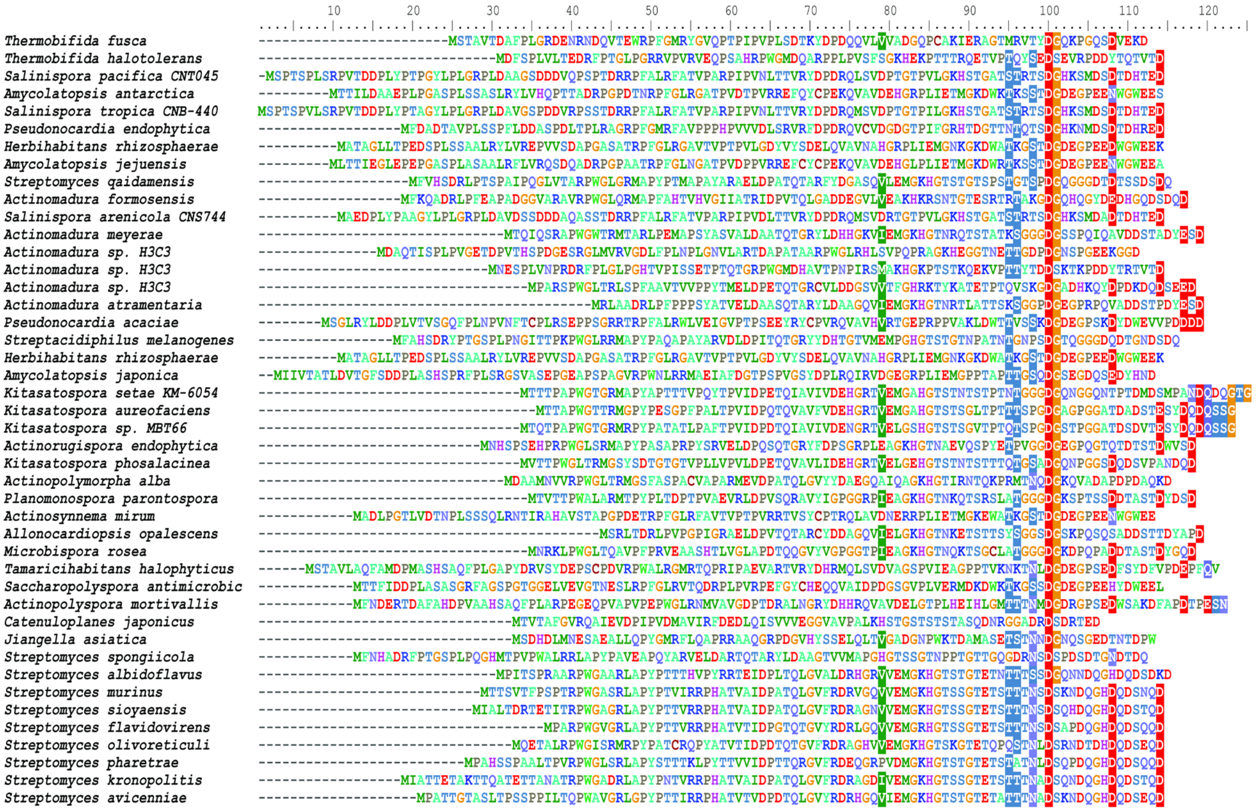


**Fig. S34** Sequence alignment of ThfA and 49 homologs. The sequences were manually aligned to show the highly conserved D residue that is the site of aspartimidylation in fuscimiditide. Note that residue following the conserved D is always G or S.

**Supplementary Tables:**

**Table S1: Primers used in this study.**

| | **Primer Sequence (5' to 3')** | **Description** |
|---|---|---|
| 1 | TATGGATCCGATGTCGACAGCGGTCACC | ThfA forward |
| 2 | ATAAAGCTTCTAGTCTTTTTCCACGTCGC | ThfA reverse |
| 3 | TATGAATTCATTAAAGAGGAGAAATTAACT ATGACCGTTCTCATCCTCACCAA | un-tagged ThfB forward |
| 4 | TATGGATCCATGACCGTTCTCATCCTCACC AA | His-tagged ThfB forward |
| 5 | TATAAGCTTAGTTTTCCTTCCTTGTGAGAG | ThfB reverse |
| 6 | ATAGGATCCACCACACAGCACACCTCAT | His-tagged ThfM forward |
| 7 | ATAAAGCTTTTAGAGCAGCATCGGCCA | His-tagged ThfM reverse |
| 8 | GAATTCATTAAAGAGGAGAAATTAACTATG ACCACACAGCACACCTCAT | Un-tagged ThM forward |
| 9 | GCATGGTCTCTAATTCATTAAAGAGGAGAA ATTAACTATG | Un-tagged ThM (bicistronic construct) forward |
| 10 | CGATGGTCTCTGCGGCCGCTTAGAGCAGG ATCGGCC | Un-tagged ThM (bicistronic construct) reverse |
| 11 | GCATGGTCTCTCCGCATTAAAGAGGAGAA ATTAACTATG | Un-tagged ThB (bicistronic construct) forward |
| 12 | CGATGGTCTCAAGCTTTTAGTTTTCCTTCC TTGTGAGAGC | Un-tagged ThfB (bicistronic construct) reverse |
| 13 | GGATCCGCTTGGCAGGGACGAGAACAG | Precursor 11-89 Forward |
| 14 | GGATCCGGTCACCGAGTGGCGCCCGTTC | Precursor 21-89 Forward |
| 15 | GGATCCGTACGGCGTACAGCCCACGCC | Precursor 31-89 Forward |
| 16 | CCCCTCAGCGACACCAAGTATG | Precursor 41-89 Forward |
| 17 | GGATCCGCAGCAGGTCCTCGTGGTGGC | Precursor 51-89 Forward |
| 18 | GGATCCGCCGTGCGCGAAAATCGAAAG | Precursor 61-89 Forward |
| 19 | AAGCTTCTAGTCTTTTTCCACGTCGC | Truncations Reverse |
| 20 | GGAGGGGCGGGATCCGATGTCGACAGCG | ThfA (mutagensis constructs) forward |

| | | |
|---|---|---|
| 21 | ATTATGCGGCCGTGTACAATACG | ThfA (mutagensis constructs) reverse |
| 22 | AGTGGCGCGCATTCGGAATGC | RAFG motif forward |
| 23 | GCATTCCGAATGCGCGCCACT | RAFG motif reverse |
| 24 | AGTGGGCAGCATTCGGAATGC | AAFG motif forward |
| 25 | GCATTCCGAATGCTGCCCACT | AAFG motif reverse |
| 26 | AGTGGCGCCCGTTCGCAATGC | RPFA motif forward |
| 27 | GCATTGCGAACGGGCGCCACT | RPFA motif reverse |
| 28 | AGTGGCGCCCGGCAGGAATGC | RPAG motif forward |
| 29 | GCATTCCTGCCGGGCGCCACT | RPAG motif reverse |
| 30 | ACGGCCAGGCTCCGGG | K13A forward |
| 31 | CTGGCCGTCGGGATAGGT | K13A reverse |
| 32 | TCCTGCACCAAGCTTCTAGTCTTTTTCCAC GTCTGCCTGCC | S17A reverse |
| 33 | CGCCCCTCCAGCAAGCTTCTAGTCTTTAG CCACGTCG | E20A reverse |
| 34 | GACGGCCAGCCGGCAGCGAAAAT | C62A forward |
| 35 | ACGGCTGGCCGTCTG | C62A reverse |
| 36 | AGTCACCTATCCCGCTGGCCAGAAG | D10A forward |
| 37 | GGGATAGGTGACTCTCATGGTGC | D10A reverse |
| 38 | GCACCATGAGAGTCGCATATCCCGAC | T7A forward |
| 39 | GACTCTCATGGTGCCGGCTCTTTC | T7A reverse |
| 40 | GAAAATCGAAAGAGCCGGCGCAATGAGAG T | T3A forward |
| 41 | GCCGGCTCTTTCGATTTTCGCG | T3A reverse |
| 42 | CCCCGCTCCAAGCTTCTAGTCTTTTTCCAC TGCGCTCTG | D18A reverse |
| 43 | CCCCGCTCCAAGCTTCTATGCTTTTTCCAC GTCGCTCTG | D22A reverse |
| 44 | CCCCGCTCCAAGCTTCTAGTCTGCTTCCA CGTCGCTCTG | K21A reverse |

| | | |
|---|---|---|
| 45 | GAAAATCGAAAGAGCCGGCGTTATGAGAGT | T3V forward |
| 46 | CCCCGCTCCAAGCTTCTAGTCTTTTTCCACTGCGCTCTG | D18E reverse |
| 47 | CCCCGCTCCAAGCTTCTATTCTTTTTCCACGTCGCTCTG | D22E reverse |

**Table S2. Chemical shift assignments for pre-fuscimiditide, fuscimiditide, and iso pre-fuscimiditide.**

| | | Pre-Fuscimiditide | Fuscimiditide | iso Pre-Fuscimiditide |
|---|---|---|---|---|
| **Residue** | **Atom** | **Chemical shift δ (ppm)** | **Chemical shift δ (ppm)** | **Chemical shift δ (ppm)** |
| Ala1 | HA | 4.133 | 4.113 | unresolved |
| Ala1 | HB | 1.441 | 1.441 | unresolved |
| Ala1 | CA | 49.469 | 49.149 | |
| Ala1 | CB | 16.596 | 16.437 | |
| Gly2 | H | 8.587 | 8.593 | 8.575 |
| Gly2 | HA2 | 4.046 | 4.034 | 3.96 |
| Gly2 | HA3 | 4.017 | | |
| Gly2 | C | | 171.977 | |
| Gly2 | CA | | 42.489 | |
| Thr3 | H | 8.111 | 7.861 | 8.114 |
| Thr3 | HA | unresolved | 4.811 | 4.874 |
| Thr3 | HB | 5.258 | 5.271 | 5.251 |
| Thr3 | QG2 | 1.103 | 1.072 | 1.105 |
| Thr3 | C | | 171.292 | |
| Thr3 | CB | 73.341 | 73.242 | |
| Thr3 | CG | 15.813 | 15.746 | |
| Met4 | H | 8.566 | 8.593 | 8.554 |
| Met4 | HA | 4.838 | 4.811 | 4.859 |
| Met4 | HB2 | 1.807 | 1.793 | 1.809 |
| Met4 | HB3 | 1.748 | 1.734 | 1.75 |
| Met4 | HG2 | 2.354 | 2.342 | 2.356 |
| Met4 | HG3 | 2.286 | 2.263 | 2.298 |
| Met4 | QE | 1.895 | 1.881 | unresolved |
| Met4 | CA | 46.729 | | |
| Met4 | CB | 32.252 | 32.107 | |
| Met4 | CG | 29.394 | 29.169 | |

| | | | | |
|---|---|---|---|---|
| Met4 | CE | 14.443 | 14.282 | |
| Arg5 | H | 8.91 | 8.88 | 8.972 |
| Arg5 | HA | 4.603 | 4.595 | 4.625 |
| Arg5 | HB2 | 1.66 | 1.636 | 1.672 |
| Arg5 | HB3 | 1.621 | 1.597 | 1.633 |
| Arg5 | QG | 1.416 | 1.421 | 1.398 |
| Arg5 | QD | 3.078 | 3.067 | 3.099 |
| Arg5 | HE | 7.028 | 6.974 | 7.214 |
| Arg5 | CB | 30.226 | 30.149 | |
| Arg5 | CG | 24.173 | 24.006 | |
| Arg5 | CD | 40.856 | 40.726 | |
| Val6 | H | 8.468 | 8.472 | 8.445 |
| Val6 | HA | 4.194 | 4.184 | 4.194 |
| Val6 | HB | 1.787 | 1.793 | 1.809 |
| Val6 | QG1 | 0.907 | 0.892 | 0.909 |
| Val6 | QG2 | 0.653 | 0.655 | 0.655 |
| Val6 | CA | 60.426 | | |
| Val6 | CB | 29.51 | 29.365 | |
| Val6 | CG | | 19.164 | |
| Thr7 | H | 8.13 | 8.147 | 8.095 |
| Thr7 | HA | 4.464 | 4.438 | 4.49 |
| Thr7 | HB | 5.111 | 5.096 | 5.113 |
| Thr7 | QG2 | 0.937 | 0.911 | 0.948 |
| Thr7 | C | | 169.334 | |
| Thr7 | CA | 54.36 | | |
| Thr7 | CB | 72.917 | 72.655 | |
| Thr7 | CG | 16.204 | 16.045 | |
| Tyr8 | H | 8.604 | 8.63 | 8.635 |
| Tyr8 | HA | 4.867 | 4.85 | 4.781 |

| | | | | |
|---|---|---|---|---|
| Tyr8 | HB2 | 2.951 | 2.871 | 3.002 |
| Tyr8 | HB3 | 2.618 | 2.636 | 2.5 |
| Tyr8 | QD | 7.032 | 7.016 | 7.043 |
| Tyr8 | QE | 6.666 | 6.652 | 6.659 |
| Tyr8 | CA | 55.73 | | |
| Tyr8 | CB | 35.865 | | |
| Tyr8 | CD | 130.755 | 130.678 | |
| Tyr8 | CE | 115.215 | 115.16 | |
| Pro9 | HA | 4.33 | 4.359 | 4.337 |
| Pro9 | HB2 | 2.159 | 2.146 | 1.876 |
| Pro9 | HB3 | 1.836 | 1.871 | 1.825 |
| Pro9 | HG2 | 1.797 | unresolved | unresolved |
| Pro9 | HG3 | 1.748 | 1.773 | 1.77 |
| Pro9 | HD2 | 3.586 | 3.615 | 3.573 |
| Pro9 | HD3 | 3.205 | 3.165 | 3.177 |
| Pro9 | C | | 174.423 | |
| Pro9 | CA | 60.734 | | |
| Pro9 | CB | 29.429 | 29.295 | |
| Pro9 | CG | 24.354 | | |
| Pro9 | CD | 47.607 | 47.472 | |
| Asp10 | H | 8.61 | 8.719 | 7.887 |
| Asp10 | HA | 4.682 | 4.673 | 4.449 |
| Asp10 | HB2 | 2.814 | 3.184 | 2.689 |
| Asp10 | HB3 | 2.657 | 2.734 | 2.552 |
| Asp10 | C | 172.997 | 176.674 | |
| Asp10 | CB | 36.578 | 34.739 | |
| Asp10 | CG | 174.346 | 177.066 | |
| Gly11 | H | 8.301 | unresolved | 8.268 |
| Gly11 | HA2 | 3.919 | 4.308 | 3.784 |

| | | | | |
|---|---|---|---|---|
| Gly11 | HA3 | 3.772 | 4.184 | |
| Gly11 | C | 171.648 | 168.6 | |
| Gly11 | CA | 43.041 | 40.53 | |
| Gln12 | H | 8.172 | 8.625 | 8.344 |
| Gln12 | HA | 4.124 | 3.929 | 4.175 |
| Gln12 | HB2 | 2.081 | 1.95 | 2.043 |
| Gln12 | HB3 | 1.836 | 1.793 | 2.906 |
| Gln12 | HG2 | 2.188 | 2.165 | 2.278 |
| Gln12 | HG3 | 2.081 | 2.087 | 2.219 |
| Gln12 | C | | 172.906 | |
| Gln12 | CA | 53.382 | | |
| Gln12 | CB | 26.039 | 25.478 | |
| Gln12 | CG | 31.285 | 31.128 | |
| Lys13 | H | 7.75 | 7.678 | 7.85 |
| Lys13 | HA | 4.525 | 4.556 | 4.586 |
| Lys13 | HB2 | 1.709 | 1.695 | 1.672 |
| Lys13 | HB3 | 1.582 | 1.519 | 1.535 |
| Lys13 | QG | 1.328 | 1.264 | 1.242 |
| Lys13 | QD | 1.582 | unresolved | unresolved |
| Lys13 | QE | 2.892 | 2.879 | 2.865 |
| Lys13 | QZ | unresolved | 2.641 | unresolved |
| Lys13 | C | | 172.955 | |
| Lys13 | CA | 51.425 | | |
| Lys13 | CB | 30.003 | 29.887 | |
| Lys13 | CG | 21.968 | | |
| Lys13 | CD | 26.575 | | |
| Lys13 | CE | 39.365 | 36.025 | |
| Pro14 | HA | 4.295 | 4.292 | 4.394 |
| Pro14 | HB2 | 2.19 | 2.165 | 2.2 |

| | | | | |
|---|---|---|---|---|
| Pro14 | HB3 | 1.826 | 1.871 | 1.848 |
| Pro14 | HG2 | 1.973 | 1.95 | 1.946 |
| Pro14 | HG3 | 1.905 | unresolved | unresolved |
| Pro14 | HD2 | 3.682 | 3.615 | 3.625 |
| Pro14 | HD3 | 3.537 | 3.517 | 3.51 |
| Pro14 | C | 174.878 | | |
| Pro14 | CA | 61.155 | | |
| Pro14 | CB | 29.146 | 29.176 | |
| Pro14 | CG | 24.814 | 24.468 | |
| Pro14 | CD | 47.886 | 47.659 | |
| Gly15 | H | 8.5 | 8.499 | 8.576 |
| Gly15 | HA2 | 4.002 | 3.929 | 3.982 |
| Gly15 | HA3 | 3.814 | 3.85 | |
| Gly15 | C | 171.199 | 171.683 | |
| Gly15 | CA | 42.725 | 42.685 | |
| Gln16 | H | 7.894 | 7.92 | 7.98 |
| Gln16 | HA | 4.457 | 4.399 | 4.41 |
| Gln16 | HB2 | 2.081 | 2.067 | 2.043 |
| Gln16 | HB3 | 1.905 | 1.891 | 1.965 |
| Gln16 | QG | 2.257 | 2.224 | 2.258 |
| Gln16 | CA | 52.599 | | |
| Gln16 | CB | 26.982 | 26.623 | |
| Gln16 | CG | 31.185 | 31.128 | |
| Ser17 | H | 8.294 | 8.118 | 8.334 |
| Ser17 | HA | 4.642 | 4.575 | 4.664 |
| Ser17 | HB2 | 3.831 | 3.815 | unresolved |
| Ser17 | HB3 | 3.772 | 3.792 | 3.784 |
| Ser17 | CB | 62.393 | 62.077 | |
| Asp18 | H | 8.665 | 8.689 | 8.649 |

| | | | | |
|---|---|---|---|---|
| Asp18 | HA | 4.897 | 4.889 | 4.879 |
| Asp18 | HB2 | 3.194 | 3.165 | 3.217 |
| Asp18 | HB3 | 2.745 | 2.734 | unresolved |
| Asp18 | CA | | | |
| Asp18 | CB | 39.865 | 34.739 | |
| Asp18 | CG | 169.063 | 169.138 | |
| Val19 | H | 8.359 | 8.368 | 8.348 |
| Val19 | HA | 4.897 | 4.889 | 4.899 |
| Val19 | HB | 1.817 | 1.793 | 1.809 |
| Val19 | QQG | 0.741 | 0.715 | 0.753 |
| Val19 | C | | 172.026 | |
| Val19 | CB | 33.032 | 32.891 | |
| Val19 | CG | 17.965 | 17.808 | |
| Glu20 | H | 8.881 | 8.858 | 8.914 |
| Glu20 | HA | 4.623 | 4.634 | 4.605 |
| Glu20 | HB2 | 1.807 | 1.793 | 1.75 |
| Glu20 | HB3 | 1.729 | 1.715 | 1.711 |
| Glu20 | QG | 2.191 | 2.205 | 2.043 |
| Glu20 | CB | 28.912 | 28.582 | |
| Glu20 | CG | 30.108 | | |
| Lys21 | H | 8.502 | 8.534 | 8.477 |
| Lys21 | HA | 4.674 | 4.654 | 4.683 |
| Lys21 | HB2 | 1.875 | 1.832 | 1.8872 |
| Lys21 | HB3 | 1.68 | 1.675 | unresolved |
| Lys21 | QG | 1.445 | 1.421 | 1.437 |
| Lys21 | QD | 1.622 | 1.617 | 1.672 |
| Lys21 | QE | 2.915 | 2.89 | 2.912 |
| Lys21 | CB | 30.747 | 30.492 | |
| Lys21 | CG | 22.679 | | |

| | | | | |
|---|---|---|---|---|
| Lys21 | CD | 26.575 | 26.427 | |
| Lys21 | CE | 39.355 | 39.159 | |
| Asp22 | H | 8.077 | 8.064 | 8.07 |
| Asp22 | HA | 4.328 | 4.477 | 4.331 |
| Asp22 | HB2 | 3.041 | 3.067 | 3.06 |
| Asp22 | HB3 | 2.477 | 2.518 | 2.454 |
| Asp22 | C | | 176.185 | |
| Asp22 | CA | 50.251 | | |
| Asp22 | CB | 38.578 | 37.984 | |
| Asp22 | CG | 169.513 | 169.432 | |

**Table S3. NOE correlations between non-adjacent residues for fuscimiditide.**

| **Residue** | **Atom** | **Residue** | **Atom** |
|---|---|---|---|
| Thr7 | H | Asp18 | HA |
| Thr7 | H | Val19 | HG |
| Asp18 | H | Val6 | HA |
| Asp18 | H | Thr7 | H |
| Glu20 | H | Thr7 | H |
| Glu20 | H | Thr7 | HB |
| Glu20 | H | Val6 | HA |
| Asp22 | H | Gly2 | HA |
| Asp22 | H | Thr3 | HA |

**Table S4. Chemical shift differences between pre-fuscimiditide and fuscimiditide/ iso pre-fuscimiditide.**

| Residue | Atom | Δ ppm Fuscimiditide | Δ ppm iso pre-Fuscimiditide |
|---|---|---|---|
| Ala1 | H | | |
| Ala1 | HA | -0.02 | |
| Ala1 | CA | -0.32 | |
| Ala1 | CB | -0.159 | |
| Gly2 | H | 0.006 | -0.012 |
| Gly2 | HA2 | -0.012 | -0.086 |
| Thr3 | H | -0.25 | 0.003 |
| Thr3 | HB | 0.013 | -0.007 |
| Thr3 | QG2 | -0.031 | 0.002 |
| Thr3 | CB | -0.099 | |
| Thr3 | CG | -0.067 | |
| Met4 | H | 0.027 | -0.012 |
| Met4 | HA | -0.027 | 0.021 |
| Met4 | HB2 | -0.014 | 0.002 |
| Met4 | HB3 | -0.014 | 0.002 |
| Met4 | HG2 | -0.012 | 0.002 |
| Met4 | HG3 | -0.023 | 0.012 |
| Met4 | QE | -0.014 | |
| Met4 | CB | -0.015 | |
| Met4 | CG | -0.025 | |
| Met4 | CE | -1.16 | |
| Arg5 | H | -0.03 | 0.062 |
| Arg5 | HA | -0.008 | |
| Arg5 | HB2 | -0.024 | 0.012 |
| Arg5 | HB3 | -0.024 | 0.012 |
| Arg5 | QG | 0.005 | -0.018 |

| | | | |
|---|---|---|---|
| Arg5 | QD | -0.011 | 0.021 |
| Arg5 | HE | -0.054 | 0.186 |
| Arg5 | CB | -0.077 | |
| Arg5 | CG | -0.167 | |
| Arg5 | CD | -0.13 | |
| Val6 | H | 0.004 | -0.023 |
| Val6 | HA | -0.01 | 0 |
| Val6 | HB | 0.006 | 0.022 |
| Val6 | QG1 | -0.015 | 0.002 |
| Val6 | QG2 | 0.002 | 0.002 |
| Val6 | CB | -0.145 | |
| Thr7 | H | 0.017 | -0.035 |
| Thr7 | HA | -0.026 | |
| Thr7 | HB | -0.015 | 0.002 |
| Thr7 | QG2 | -0.026 | 0.011 |
| Thr7 | CB | -0.262 | |
| Thr7 | CG | -0.159 | |
| Tyr8 | H | 0.026 | |
| Tyr8 | HA | -0.017 | -0.086 |
| Tyr8 | HB2 | -0.08 | |
| Tyr8 | HB3 | 0.018 | -0.118 |
| Tyr8 | QD | -0.016 | 0.011 |
| Tyr8 | QE | -0.014 | -0.007 |
| Tyr8 | CD | -0.077 | |
| Tyr8 | CE | -0.055 | |
| Pro9 | HA | 0.029 | 0.007 |
| Pro9 | HB2 | -0.013 | -0.283 |
| Pro9 | HB3 | 0.035 | -0.011 |
| Pro9 | HG3 | 0.025 | 0.022 |

| | | | |
|---|---|---|---|
| Pro9 | HD2 | 0.029 | -0.013 |
| Pro9 | HD3 | -0.04 | -0.028 |
| Pro9 | CB | -0.134 | |
| Pro9 | CD | -0.135 | |
| Asp10 | H | 0.109 | -0.723 |
| Asp10 | HA | -0.009 | -0.233 |
| Asp10 | HB2 | 0.37 | -0.125 |
| Asp10 | HB3 | 0.077 | |
| Asp10 | C | 3.677 | |
| Asp10 | CB | -1.839 | |
| Asp10 | CG | 2.72 | |
| Gly11 | H | | -0.033 |
| Gly11 | HA2 | 0.389 | -0.135 |
| Gly11 | HA3 | 0.412 | |
| Gly11 | C | -3.048 | |
| Gly11 | CA | -2.511 | |
| Gln12 | H | 0.453 | 0.172 |
| Gln12 | HA | -0.195 | 0.051 |
| Gln12 | HB2 | -0.131 | -0.038 |
| Gln12 | HB3 | -0.043 | |
| Gln12 | HG2 | -0.023 | 0.09 |
| Gln12 | HG3 | 0.006 | 0.138 |
| Gln12 | CB | -0.561 | |
| Gln12 | CG | -0.157 | |
| Lys13 | H | -0.072 | |
| Lys13 | HA | 0.031 | 0.061 |
| Lys13 | HB2 | -0.014 | -0.037 |
| Lys13 | HB3 | -0.063 | -0.047 |
| Lys13 | QG | -0.064 | -0.086 |

| | | | |
|---|---|---|---|
| Lys13 | QE | -0.013 | -0.027 |
| Lys13 | CB | -0.116 | |
| Pro14 | HA | -0.003 | 0.099 |
| Pro14 | HB2 | -0.025 | 0.01 |
| Pro14 | HB3 | 0.045 | 0.022 |
| Pro14 | HG2 | -0.023 | -0.027 |
| Pro14 | HD2 | -0.067 | -0.057 |
| Pro14 | HD3 | -0.02 | -0.027 |
| Pro14 | CB | 0.03 | |
| Pro14 | CG | -0.346 | |
| Pro14 | CD | -0.227 | |
| Gly15 | H | -0.001 | 0.076 |
| Gly15 | HA2 | -0.073 | -0.02 |
| Gly15 | HA3 | 0.036 | |
| Gly15 | C | 0.484 | |
| Gly15 | CA | -0.04 | |
| Gln16 | H | 0.026 | 0.086 |
| Gln16 | HA | -0.058 | -0.047 |
| Gln16 | HB2 | -0.014 | -0.038 |
| Gln16 | HB3 | -0.014 | 0.06 |
| Gln16 | QG | -0.033 | 0.001 |
| Gln16 | CB | -0.359 | |
| Gln16 | CG | -0.057 | |
| Ser17 | H | -0.176 | 0.04 |
| Ser17 | HA | -0.067 | 0.022 |
| Ser17 | HB2 | -0.016 | |
| Ser17 | HB3 | 0.02 | 0.012 |
| Ser17 | CB | -0.316 | |
| Asp18 | H | 0.024 | -0.016 |

| | | | |
|---|---|---|---|
| Asp18 | HA | -0.008 | -0.018 |
| Asp18 | HB2 | -0.029 | 0.023 |
| Asp18 | HB3 | -0.011 | |
| Asp18 | CG | 0.075 | |
| Val19 | H | 0.009 | -0.011 |
| Val19 | HA | -0.008 | 0.002 |
| Val19 | HB | -0.024 | -0.008 |
| Val19 | QQG | -0.026 | 0.012 |
| Val19 | CB | -0.141 | |
| Val19 | CG | -0.157 | |
| Glu20 | H | -0.023 | 0.033 |
| Glu20 | HA | 0.011 | -0.018 |
| Glu20 | HB2 | -0.014 | -0.057 |
| Glu20 | HB3 | -0.014 | -0.018 |
| Glu20 | QG | 0.014 | -0.148 |
| Glu20 | CB | -0.33 | |
| Lys21 | H | 0.032 | |
| Lys21 | HA | -0.02 | 0.009 |
| Lys21 | HB2 | -0.043 | 0.0122 |
| Lys21 | HB3 | -0.005 | |
| Lys21 | QG | -0.024 | -0.008 |
| Lys21 | QD | -0.005 | 0.05 |
| Lys21 | QE | -0.025 | -0.003 |
| Lys21 | CB | -0.255 | |
| Lys21 | CD | -0.148 | |
| Lys21 | CE | -0.196 | |
| Asp22 | H | -0.013 | -0.007 |
| Asp22 | HA | 0.149 | 0.003 |
| Asp22 | HB2 | 0.026 | 0.019 |

| | | | |
|---|---|---|---|
| Asp22 | HB3 | 0.041 | -0.023 |
| Asp22 | CB | -0.594 | |
| Asp22 | CG | -0.081 | |
